# Universal prediction of cell cycle position using transfer learning

**DOI:** 10.1101/2021.04.06.438463

**Authors:** Shijie C. Zheng, Genevieve Stein-O’Brien, Jonathan J. Augustin, Jared Slosberg, Giovanni A. Carosso, Briana Winer, Gloria Shin, Hans T. Bjornsson, Loyal A. Goff, Kasper D. Hansen

**Affiliations:** Department of Biostatistics, Johns Hopkins Bloomberg School of Public Health; Department of Genetic Medicine, Johns Hopkins School of Medicine; Department of Neuroscience, Johns Hopkins School of Medicine; Kavli Neurodiscovery Institute, Johns Hopkins University; Division of Biostatistics and Bioinformatics, Department of Oncology, Johns Hopkins School of Medicine; Department of Pediatrics, Johns Hopkins University School of Medicine; Faculty of Medicine, University of Iceland; Landspitali University Hospital

**Author notes:** Correspondence to (LAG), (KDH).

## Abstract

**Background:** The cell cycle is a highly conserved, continuous process which controls faithful replication and division of cells. Single-cell technologies have enabled increasingly precise measurements of the cell cycle both as a biological process of interest and as a possible confounding factor. Despite its importance and conservation, there is no universally applicable approach to infer position in the cell cycle with high-resolution from single-cell RNA-seq data.

**Results:** Here, we present tricycle, an R/Bioconductor package, to address this challenge by leveraging key features of the biology of the cell cycle, the mathematical properties of principal component analysis of periodic functions, and the use of transfer learning. We estimate a cell cycle embedding using a fixed reference dataset and project new data into this reference embedding; an approach that overcomes key limitations of learning a dataset dependent embedding. Tricycle then predicts a cell-specific position in the cell cycle based on the data projection. The accuracy of tricycle compares favorably to gold-standard experimental assays, which generally require specialized measurements in specifically constructed *in vitro* systems. Using internal controls which are available for any dataset, we show that tricycle predictions generalize to datasets with multiple cell types, across tissues, species and even sequencing assays.

**Conclusions:** Tricycle generalizes across datasets, is highly scalable and applicable to atlas-level single-cell RNA-seq data.

## INTRODUCTION

The cell cycle is the biological process which controls faithful replication and division of cells across all species of life. Despite existing as a continuous process, cell cycle has historically been characterized as having four discrete stages during which the cell performs growth and maintenance (G1), replicates its DNA (S), increases further in size and prepares for mitosis (G2), and undergoes mitosis and cytokinesis (M). Cell cycle is a highly conserved mechanism with and integral role in generating the diversity of cell types within multicellular organisms. As a result, maladaptive modifications of the cell cycle can have devastating consequences in development and disease (McConnell and Kaznowski, 1991; Ambros, 1999; Ohnuma and Harris, 2003). Despite its importance, many of the molecular mechanisms regulating and interacting with cell cycle remain poorly understood.

High-throughput expression data has been utilized for studying the cell cycle since the seminal work on the yeast cell cycle by Spellman et al. (1998) and Cho et al. (1998) at the dawn of the microarray era. This work used various approaches to synchronize cells in specific cell cycle stages followed by assaying cells in bulk. The data from Spellman et al. (1998) were later used by Alter et al. (2000) to show that principal component analysis reveals a circular pattern which represents the cyclical nature of the cell cycle; widely cited as one of the first examples of the use of principal component analysis and singular value decomposition in analysis of high-throughput expression data. Subsequent work sought to systematically identify both periodically expressed genes and cell cycle marker genes and deposited these into widely used databases (Whitfield et al., 2002; Gauthier et al., 2008).

Single-cell technologies have enabled the ability to study the effects of cell cycle in multicellular organisms with a degree of sensitivity and accuracy only previously available in monocelluar or clonal systems. Thus, cell cycle has been the subject of substantial interest, both as a biological variable of interest and as a possible confounding feature for other comparisons of interest (Buettner et al., 2015). A number of methods have been developed to estimate cell cycle state from single-cell expression data (Leng et al., 2015; Scialdone et al., 2015; Liu et al., 2017; Stuart et al., 2019; Hsiao et al., 2020; Schwabe et al., 2020); some of these approaches are related to more general methods for finding topological structure in single-cell data (Rizvi et al., 2017). These methods differ broadly in the definition of cell cycle state (discrete stages vs. continuous pseudotime) as well as the use of special training data. Most of these methods have been demonstrated to be effective on datasets consisting of a single cell type. Despite the conservation of the cell cycle process, none of these methods have been shown to be applicable across single-cell technologies and mammalian tissues.

## RESULTS

### Transfer learning

To develop a universal method for estimating a continuous cell cycle pseudotime for a single-cell expression data set independent of technology, cell type, or species, we leverage transfer learning via dimensionality reduction (Pan et al., 2008; Pan and Yang, 2009). We define a reference cell cycle embedding (or latent space) into which we project a new data set; an approach originally advocated for in Stein-O’Brien et al. (2019). After projection, we infer cell cycle pseudotime as the polar angle around the origin. This pseudotime variable takes values in [0, 2*π*] and is unrelated to wall time (time measured by a clock in SI units), but rather represents progression through the cell cycle phases. We refer to this pseudotime variable as cell cycle position to avoid confusion with wall time and to emphasize its periodic nature.

To define a reference cell cycle embedding, we leverage key features of principal component analysis of cell cycle genes. Previous work has found that principal component analysis on expression data sometimes yields an ellipsoid pattern. This was first described by Alter et al. (2000); it has later been observed independently in multiple data sets (Schwabe et al., 2020; Liu et al., 2017; Mahdessian et al., 2021). Here, we draw attention to the fact that the ellipsoid pattern is a consequence of a link between Fourier analysis of periodic functions and principal component analysis which links the progression through the cell cycle process with angular position on the ellipsoid. This is similar in spirit to previous observations of principal component analysis of genotype data (Novembre and Stephens, 2008) and connected to mathematical results on circulant matrices (Gray, 2005).

We use the first two principal components to define a reference embedding representing the cell cycle. Because this reference embedding is a lowdimensional linear space, we obtain an orthogonal projection operator allowing us to project any new data set into the reference embedding. We show that projecting new data into the reference cell cycle embedding overcomes technical and biological challenges posed by data sets where substantial variation is explained by one or more factors different from cell cycle, such as cellular differentiation.

### Principal component analysis and periodic functions

To gain insight into gene expression dynamics over the cell cycle, we start by analyzing principal component analysis of periodic functions. Our model is a collection of periodic functions with a single peak, taking the form

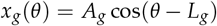

with a gene-specific amplitude (*A_g_*) and location of the peak (*L_g_*) with 0 ≤ *θ* < 2*π* representing the unknown cell cycle position. Figure 1a,b depicts the unobserved (true) time ordering, observed on a discrete grid of time points, together with a random permutation of these time points; this represents the observed data which is not ordered by time. A key insight is the fact that the first two principal components are the same for the observed and the unobserved data (Figure 1c), when performed on a discrete set of observation times. The unknown time order can be inferred from the principal component plot as the angle of each point, making it possible to fully reconstruct the unobserved time order (Figure 1d), i.e., the first two principal components form an orthogonal projection into a twodimensional space representing the periodic time. For this result to hold, at least two peak locations (which are not separated by exactly *π*) are required. Our Methods section contains a constructive proof in the case of single peak genes, which are particular relevant for cell cycle expression.

**Figure 1.**
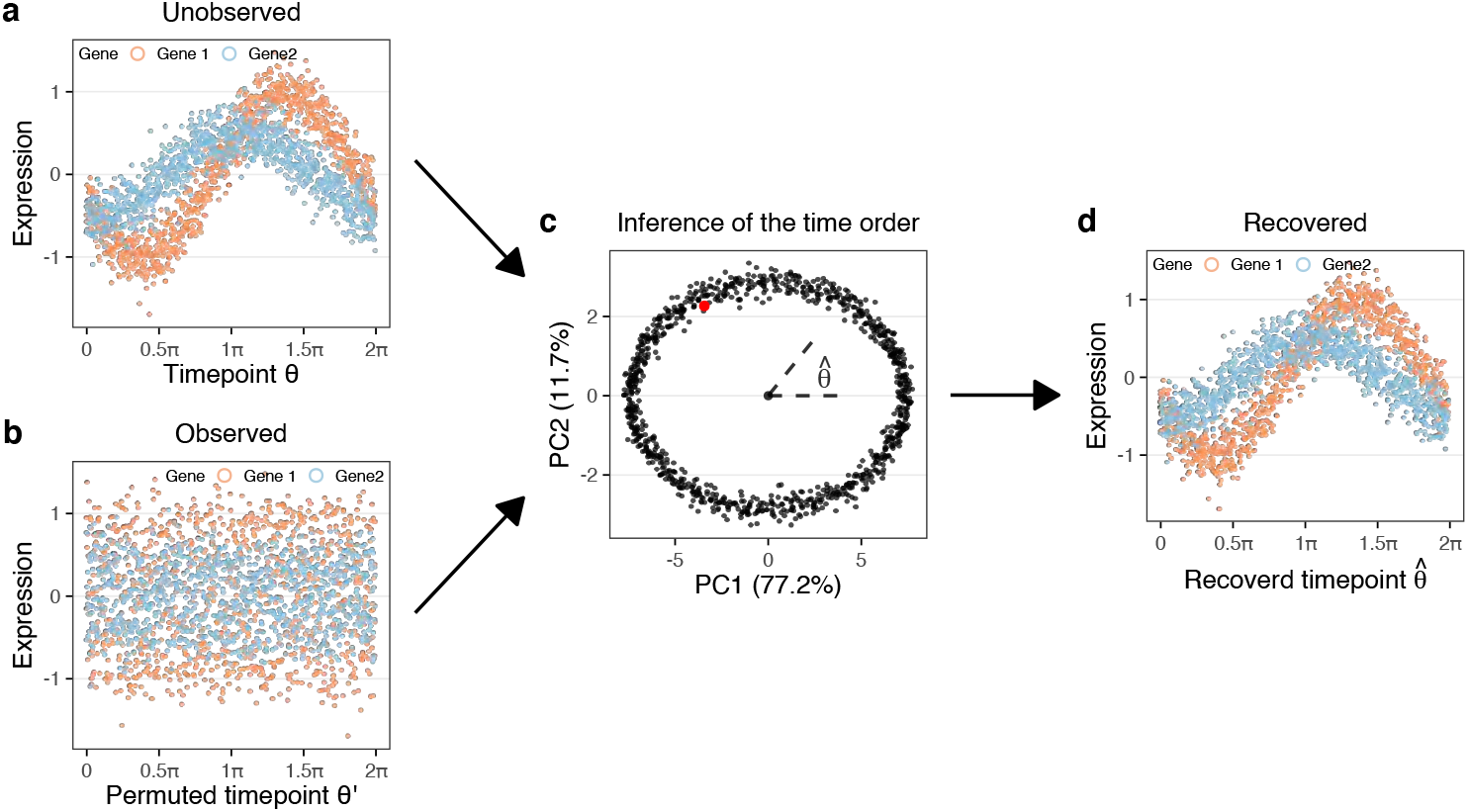
Principal component analysis recovers time ordering in simulations. Simulations are based on cosine functions with Gaussian noise (Methods). **(a)** Expression vs. time for 2 genes with different peak locations and amplitudes. Each of the two gene peaks are replicated 50 times for a total of 500 genes and 1,000 time points (cells). **(b)** Expression vs. permuted time, representing the unknown time order of observed data which obscures the periodicity of the functions. **(c)** Principal component analysis of the data from (b) and (a); the two datasets have equivalent principal components. We infer cell cycle position 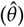 by the angle of the ellipsoid. The red dot indicates *θ* = 0. **(d)** Expression vs. inferred cell cycle position.

The simulated data depicted in Figure 1 has Gaussian noise, but we have verified that the result holds for data generated using the negative binomial distribution with an associated mean-variance relationship. Using the negative binomial distributed data required more than 2 distinct peaks to be stable (Supplementary Figures S1, S2). For both distributions, this approach is robust to downsampling of the data similar to what is seen with the increased sparsity from droplet based sequencing technology. In simulations, we can recover cell cycle position with as little as 10 total counts per cell across 100 genes (depending on noise levels and heights of the peaks) (Supplementary Figure S3).

### Recovering cell cycle position using principal component analysis on cell cycle genes

We next assess our model on experimental data, and learn an embedding representing cell cycle. We use 10x Genomics Chromium single-cell RNA-sequencing (scRNA-seq) data on two replicate cultures of E14.5 mouse cortical neurospheres (Methods), integrated using Seurat 3 and transformed to log_2_-scale. The use of an alignment method (CCA in Seurat3) to integrate the two samples is important for the quality of the ellipsoid, by maximizing the correlation structure between the two samples. Since neurospheres are maintained in a proliferative state, we expect that cell cycle phase is an important contributor to the variation in expression within this single-cell dataset. To confirm this expectation, we consider a UMAP representation of the data based on all variable genes (Supplementary Figure S4) colored according to the predictions from two separate cell cycle stage estimation utilities (cyclone and a modification of Schwabe et al. (2020) we call SchwabeCC, see Methods); this analysis demonstrates that the cell cycle is a major source of transcriptional variation in the neurosphere dataset.

We then perform principal component analysis of the top 500 most variable genes amongst the roughly 1700 genes annotated with the Gene Ontology cell cycle term (*GO:0007049*, Methods) (Ashburner et al., 2000). As suggested by our model, the first two principal components form an ellipsoid with a sparse/empty interior (Figure 2a). Using the SchwabeCC cell cycle stage predictor, we observe a strong relationship between polar angle on the ellipsoid and predicted cell cycle stage.

**Figure 2.**
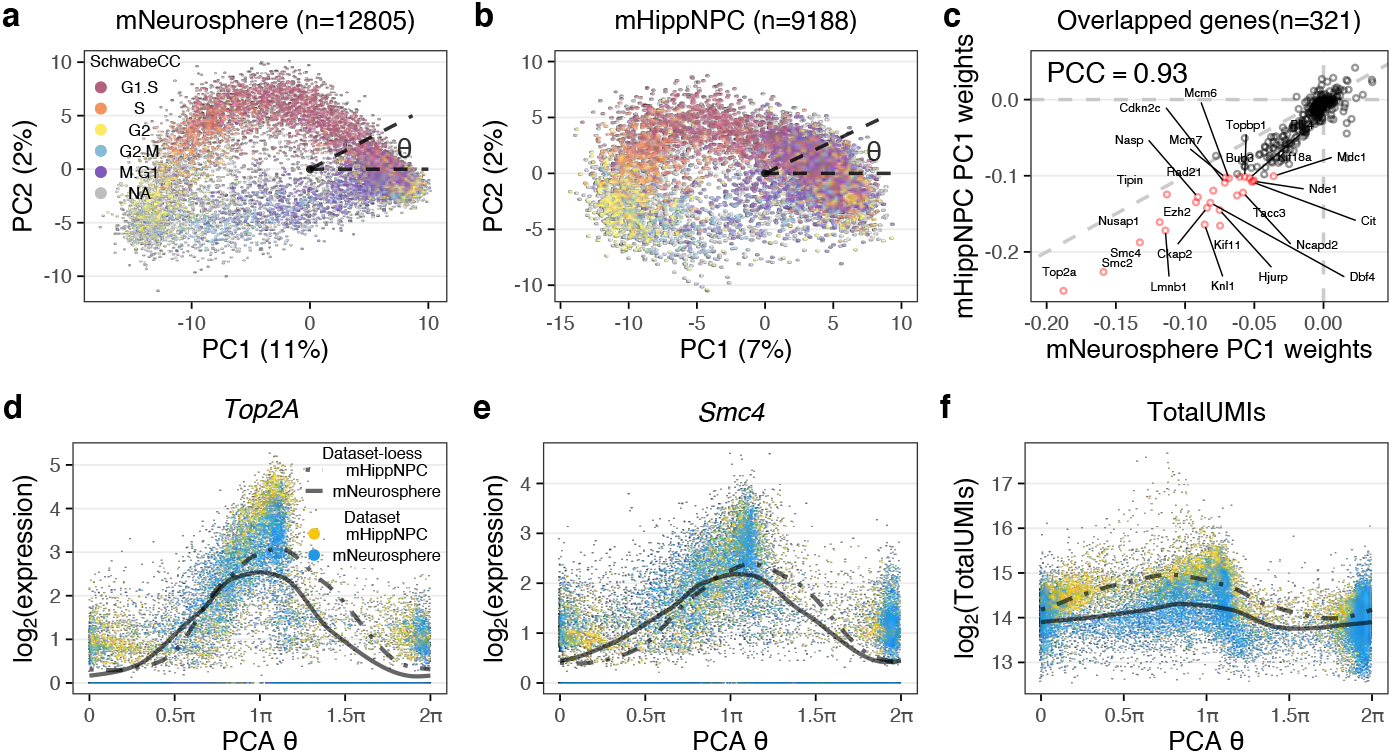
The cell cycle ellipsoid and cell cycle position. **(a)** Top 2 principal components of GO cell cycle genes from E14.5 primary mouse cortical neurospheres, in which the variation is primarily driven by cell cycle. Each point represents a single cell, which is colored by 5 stage cell cycle representation, inferred using the SchwabeCC method (Schwabe et al., 2020). The cell cycle position *θ* (with values in [0, 2*π*); sometimes called cell cycle pseudotime) is the polar angle. **(b)** As in (a), but for a dataset of primary mouse hippocampal progenitor cells from both a mouse model of Kabuki syndrome and a wildtype. **(c)** A comparison of the weights on principal component 1 between the cortical neurosphere and hippocampal progenitor datasets. Genes with high weights (|score| > 0.1 for either vector) are highlighted in red. PCC: Pearson Correlation Coefficient. **(d,e)** The expression dynamics of (d) Top2A and (e) Smc4 using the inferred cell cycle position, with a periodic loess line (Methods). **(f)** The dynamics of total UMI using the inferred cell cycle position, with a periodic loess line, illustrating the high agreement of the dynamics between datasets.

The strong relationship between polar angle on the ellipsoid and predicted cell cycle stage was also observed on an independent dataset on cultured primary mouse hippocampal progenitors from a wild-type mouse as well as from a *Kmt2d^+/βgeo^* mouse, a previously described model of Kabuki syndrome (Carosso et al., 2019). The data were processed similarly to the neurosphere data. Again, we select the top 500 most variable cell cycle genes and perform a principal component analysis (Figure 2b) which reveal an ellipsoid pattern. The shape of the principal component plot differs between the two datasets, but the weights used to form the first two principal components are highly concordant (Figure 2c, Supplementary Figure S5 for PC2) for the 321 genes present in both cell cycle embeddings. Almost all of the highly ranked genes (absolute weights > 0.1, highlighted in red and labelled with gene name) represent important regulators of, or participants in, the cell cycle. For example, the highest ranked gene is Topoisomerase 2A *Top2A* which controls the topological state of DNA strands and catalyzes the breaking and rejoining of DNA to relieve supercoiling tension during DNA replication and transcription (Lodish et al., 2008). Also highly ranked are *Smc2* and *Smc4* which compose the core subunits of condensin, which regulates chromosome assembly and segregation (Ono et al., 2003; Wei-Shan et al., 2019).

Given our mathematical analysis as well as the strong empirical relationship between polar angle on the ellipsoid and cell cycle stage predictions, we define a method to learn cell cycle position as the polar angle around the origin on the coordinate plane which we denote by *θ*. We center the coordinate plane on (0, 0) whose location corresponds to cells with zero expression for all 500 variable cell cycle genes.

To demonstrate that cell cycle position reflects the true biological cell cycle progression, we consider expression dynamics of specific cell cycle genes. For *Top2A* and *Smc2* the peak expressions are observed at G2 stage around *π* (Figure 2e), consistent with their known increased expression through S phase and into G2 (Heck et al., 1988; Belluti et al., 2013; Wei-Shan et al., 2019). Furthermore, the dynamics are highly similar between the independently analyzed cortical neurosphere and hippocampal NPC datasets, which supports the observation that the two different embeddings yield concordant cell cycle positions (despite each including dataset-specific genes). These observations hold for all genes with high weights (Supplementary Figure S6). This approach serves as an internal control in any single-cell RNA-seq data set and can be used to assess the quality of any continuous ordering.

Next, we directly relate *θ* to the measured transcription values. Figure 2f shows the log_2_ transformed total UMI numbers against *θ*, with a periodic loess smoother for each dataset. In both datasets, the maximum level is reached around *π* and the minimum around 1.5*π*, which corresponds to the end of G2 and the middle of M stage respectively. We observe the total UMI number begins to increase at the beginning of G1/S phase and to decrease sharply as cells progress through M phase. The difference between the maximum and minimum of the periodic loess line is 1, corresponding to a two-fold difference in total UMI, which is known to be proportional to cell size (Marguerat and Bähler, 2012; Padovan-Merhar et al., 2015). This observation, and the timing with respect to cell cycle position, is consistent with the approximate reduction in cellular volume by one half as a result of cytokinesis in M phase and the formation of two daughter cells of roughly equal size.

This approach of using expression dynamics and (if available) log_2_-totalUMI can be used to evaluate whether *any* continuous position is related to the cell cycle. We name these “internal controls” and we note that they are available for *any* single-cell expression dataset. These internal controls will be used extensively throughout this manuscript.

Note that these principal component analyses are differentiating G2/M cells from G1/G0 cells on the first principal component. This contrasts with the mathematical analysis where the starting point (*θ* = 0) can be any location (red point in Figure 1) as there is no clear starting point for a periodic function. That the first principal component differentiates G2M from G1/G0 can be explained by the nature of principal component analysis. Before principal component analysis, we subtract each gene’s mean expression. However, genes marking G2/M usually have very high expression compared to other stages, with G0/G1 being the lowest (Supplementary Figure S7), ensuring that this becomes the first principal component. A clustering analysis of the expression patterns provides further evidence that cell cycle genes have a single peak pattern of expression (Supplementary Figure S7). Thus, the observed behavior of the cell cycle genes in these data sets fits the theoretical requirements of our model.

In summary, principal component analysis of the cell cycle genes predicts cell cycle progression for the mNeurosphere and mHippNPC datasets with a high degree of similarity between the cell cycle position inferred independently in the two datasets as predicted by our mathematical model.

### When principal component analysis fails to reflect cell cycle position

A principal component analysis does not always yield an ellipsoid pattern; a requirement for this to work is for the first principal component to be dominated by cell cycle. To illustrate this, we used an existing mouse developing pancreas dataset, with cell type labels (Bastidas-Ponce et al., 2019). A major source of variation in this dataset is cellular differentiation as demonstrated by a standard UMAP embedding (based on all variable genes) illustrating the previously described (Bastidas-Ponce et al., 2019) differentiation trajectories (Figure 3a). When we perform principal component analysis using only the variable cell cycle genes, the resulting PCA plot still reflects the differentiation trajectory and does not resemble the ellipsoid pattern observed in the previous section (Figure 3b). Note that PC1 has some relationship with cell cycle since the differentiation path goes from cycling to non-cycling cells, but it also reflects the progression from cycling multipotent cells to terminally differentiated cells. This result strongly suggests that some of the cell cycle genes may participate in biological processes other than the cell cycle and demonstrates that PCA of cell cycle genes does not always exclusively capture cell cycle variance.

**Figure 3.**
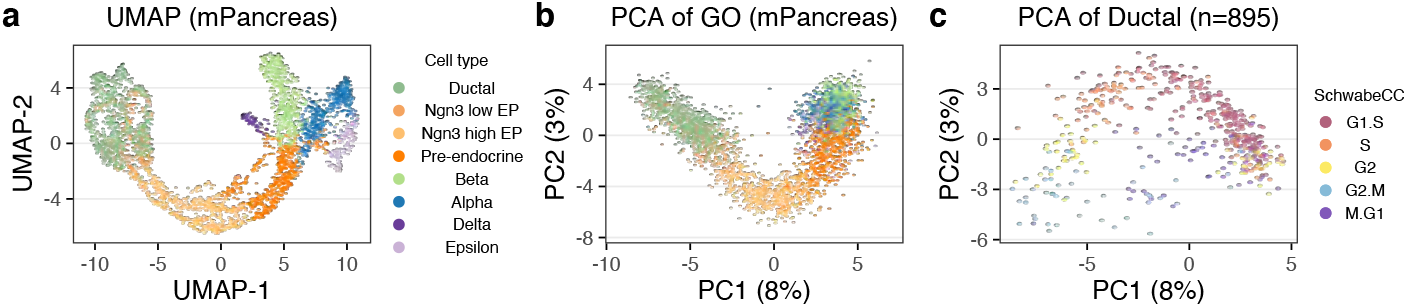
When principal component analysis fails to describe the cell cycle. Data is from the developing mouse pancreas. **(a)** UMAP embedding using all variable genes. Cells are colored by cell type. **(b)** PCA plot of the cell cycle genes; this reflects the differentiation path in (a). **(c)** PCA plot of the cell cycle genes for ductal cells only; this plot reflects cell cycle.

However, when we perform principal component analysis only on a subset of cells from a single, proliferating progenitor cell type, the ellipsoid pattern returns (Figure 3c and Supplementary Figure S8a,b). This highlights the challenge of inferring cell cycle for datasets that contain many different cell types, including postmitotic cells.

### Transfer learning through projection

To overcome the challenges of inferring cell cycle position in arbitrary datasets, we propose a simple, yet highly effective transfer learning approach we term tricycle (transferable representation and inference of cell cycle). In short, we first construct a reference embedding representing the cell cycle process using a fixed dataset where cell cycle is the primary source of transcriptional variation. For the remainder of this manuscript we will use the cortical neurosphere data as this reference. We show that the learned reference embedding generalizes across all datasets we have examined. Because our reference embedding is a linear subspace, we benefit from an orthogonal projection operator which allows us to map new data into the reference embedding, with well-understood mathematical properties. Finally, we infer cell cycle position by the polar angle around the origin of each cell in the embedding space. The robustness of this approach is demonstrated by the ability of this projection to estimate cell cycle position in multiple independent and disparate datasets; evidence of which is provided below. Specifically, using the cortical neurosphere dataset as a fixed reference, our transfer learning approach generalizes across cell types, species (human/mouse), sequencing depths and even singlecell RNA sequencing protocols.

As a demonstration, we consider a diverse selection of single-cell RNA-seq datasets representing different species (mouse and human), cell types and technologies (10x Chromium, SMARTer-Seq, Drop-seq and Fluidigm C1) (Table 1). We project these datasets into the cell cycle embedding learned from the neurosphere data (Figure 4a, Supplementary Figure S9a). To effectively visualize cell cycle position defined as the polar angle, we use a circular color scale to account for the fact that position “wrap around” from 2*π* to 0. Although the shape of the projection varies from dataset to dataset, the cells of the same stage always appear at a similar position of *θ*, such as cells at S stage centering at 0.75*π*.

**Figure 4.**
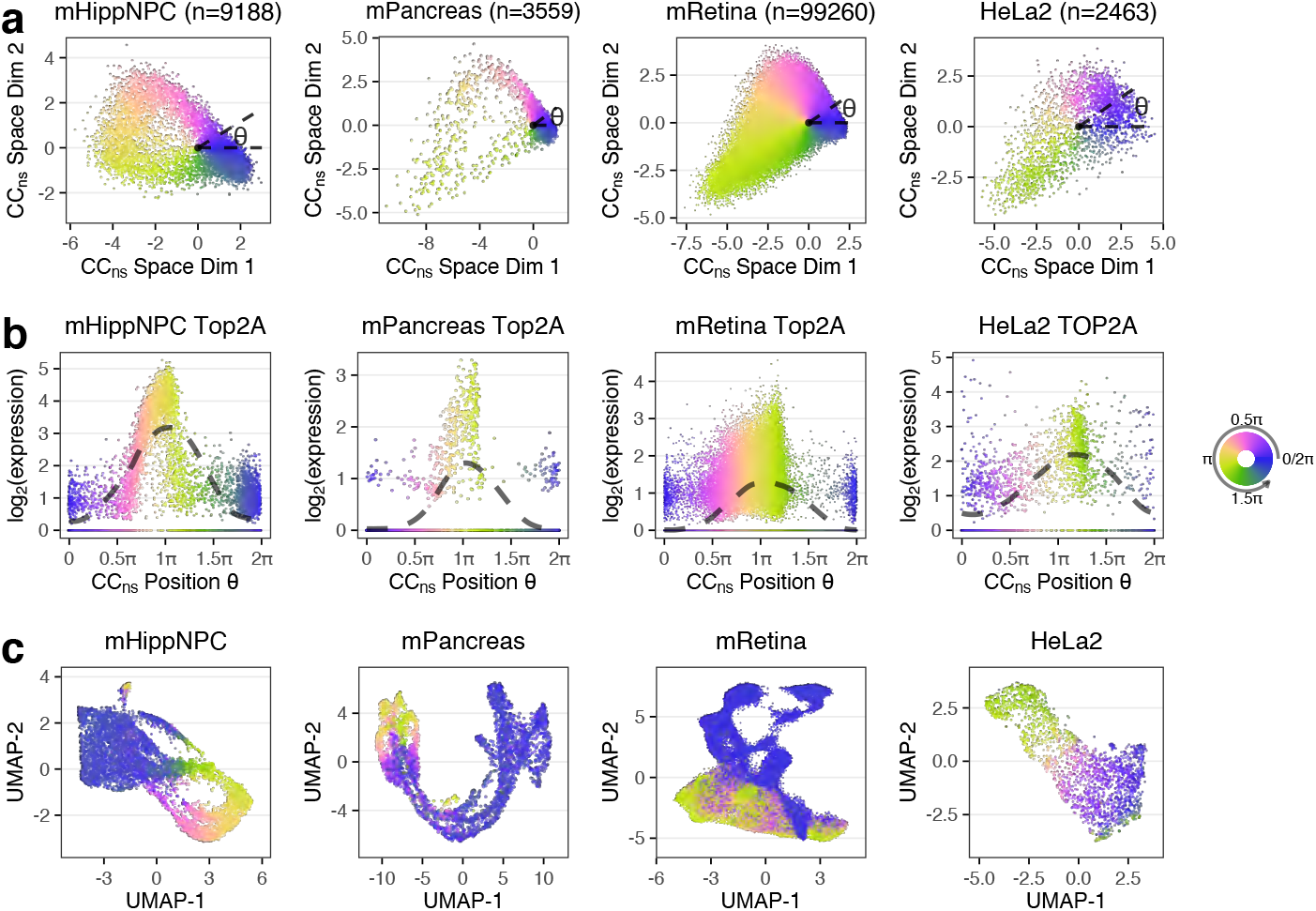
A pre-learned weights matrix learned from proliferating cortical neurospheres enables cell cycle position estimation in other proliferating datasets. **(a)** Different datasets (hippocampal NPCs, mouse pancreas, mouse retina and HeLa set 2) projected into the cell cycle embedding defined by the cortical neurosphere dataset. Cell cycle position *θ* is estimated as the polar angle. **(b)** Inferred expression dynamics of *Top2A* (*TOP2A* for human), with a periodic loess line (Methods). **(c)** UMAP embeddings of top variable genes. All the cells are colored by cell cycle position using a circular color scale.

**Table 1.** Datasets.

| Dataset | Species | Platform | # cells | Note | Reference |
| --- | --- | --- | --- | --- | --- |
| mNeurosphere | Mouse | 10x | 12805 |  |  |
| mHippNPC | Mouse | 10x | 9188 |  |  |
| mPancreas | Mouse | 10x | 3559 |  | Bastidas-Ponce et al. (2019) |
| mHSC | Mouse | SMARTer | 1343 |  | Kowalczyk et al. (2015) |
| mRetina | Mouse | 10x | 99260 |  | Clark et al. (2019) |
| HeLa 1 | Human | Drop-seq | 1398 |  | Schwabe et al. (2020) |
| HeLa 2 | Human | Drop-seq | 2463 |  | Schwabe et al. (2020) |
| mESC | Mouse | Fluidigm C1 | 279 | FACS | Buettner et al. (2015) |
| hESC | Human | Fluidigm C1 | 226 | FACS | Leng et al. (2015) |
| hU2OS | Human | SMART-seq2 | 1114 | FUCCI | Mahdessian et al. (2021) |
| hiPSCs | Human | Fluidigm C1 | 888 | FUCCI | Hsiao et al. (2020) |
| Fetal tissue atlas | Human | sci-RNA-seq3 | Varies |  | Cao et al. (2020) |

To verify our cell cycle ordering, we again use the internal controls as they exist in all datasets. Specifically, we show the expression dynamics of *Top2A* and *Smc4* as a function of *θ* (Figure 4b, Supplementary Figure S9b, Supplementary Figure S10). In contrast, PCA plots of the GO cell cycle genes for each dataset illustrates the advantage of using a fixed embedding to represent cell cycle (Supplementary Figure S11). Together, these results strongly support that tricycle generalizes across data modalities.

Having inferred cell cycle position, we can visualize the cell cycle dynamics on a UMAP plot representing the full transcriptional variation, as is standard in the scRNA-seq literature (Figure 4c, Supplementary Figure S9c). Doing so reveals the smooth behaviour of the tricycle predictions (despite not using smoothing or imputation) and argues for representing cell cycle in gene expression data as a continual progression rather than discrete states.

We draw attention to two specific datasets in these figures: the mPancreas and mRetina datasets (Figure 4) and the HCA (human cell atlas) Pancreas data (Supplementary Figure S9). These three datasets contain many different cell types and strong drivers of gene expression in addition to the cell cycle, such as differentiation. We have previously seen how principal component analysis fails to be ellipsoid on the mPancreas dataset (Figure 3) and this example shows how tricycle – by projecting the data into a fixed reference embedding – overcomes the limitations of principal component analysis. Finally, note that the HCA Pancreas dataset is sparse with a median of 892 total UMIs per cell.

### Cell cycle position estimation on gold-standard datasets

We validated tricycle on multiple datasets containing “gold-standard” cell cycle measurements, including measurements by proxy using the fluorescent ubiquitination-based cell-cycle indicator (FUCCI) system and by fluorescence-activated cell sorting (FACS) of cells into discrete cell cycle stages. Both of these approaches allow for assignment to or selection of cells from discrete phases of the cell cycle. The FUCCI system uses a dual reporter assay in which the reporters are fused to two genes with dynamic and opposing regulation during the cell cycle (Sakaue-Sawano et al., 2008), allowing for a quantitative assessment of whether cells are in G1 or S/G2/M phase. In contrast to FACS, FUCCI systems, combined with an appropriate quantification method, make it possible to continuously measure cell cycle progression by placing the 2 protein measurements in a 2-dimensional space. Cell cycle pseudotime needs to be inferred from these 2-dimensional measurements, which is usually done by a variant of polar angle (Hsiao et al., 2020; Mahdessian et al., 2021).

Mahdessian et al. (2021) measured human U-2 OS cells to derive a FUCCI-based pseudo-time scoring. Their FUCCI measurements form a distinct horseshoe shape with the left side of the horseshoe representing time post-metaphase-anaphase transition with a continuous progression through G1, S, G2 and ending pre-metaphase-anaphase transition (Figure 5; this depiction mirrors other data presentations (Sakaue-Sawano et al., 2008; Sakaue-Sawano et al., 2017)). Cell cycle is a continuous process which is not immediately reflected in the horseshoe form because of the large gap (in the x-axis) between the two ends of the horsehoe. The x-axis reflects the protein levels of geminin (GMNN) which is degraded during the metaphase-anaphase transition (McGarry and Kirschner, 1998) and the two ‘open’ ends of the horseshoe are closely connected in time despite the visual gap in the scatterplot. This fact gives the FUCCI system the ability to assess whether a cell in M phase is before or after this transition, or said differently, a high temporal resolution around this transition despite the relatively short wall time compared to the rest of the cell cycle. We observe a close correspondence between tricycle cell cycle position and FUCCI pseudotime (circular correlation coefficient *ρ* = 0.70). The only cells for which there is a apparent disagreement are placed in M phase by tricycle (cell cycle position around 0.9*π*) and are split between pre-metaphase-anaphase transition and post-metaphase-anaphase transition by FUCCI pseudotime, for this particular transition the FUCCI system has higher temporal resolution than tricycle; adding a small offset to these cells results in a remarkable concordance between the two systems (Figure 5). Elsewhere in the cell cycle, there is no evidence of better temporal resolution with FUCCI; examining expression dynamics suggests that tricycle does at least as well as FUCCI at ordering key cell cycle genes. We can use tricycle to examine the expression dynamics of *GMNN* and *CDT1* which reveals that *GMNN* expression is stable across the cell cycle (Supplementary Figure S12), suggesting the protein is predominantly regulated post-transcriptionally during mitosis.

**Figure 5.**
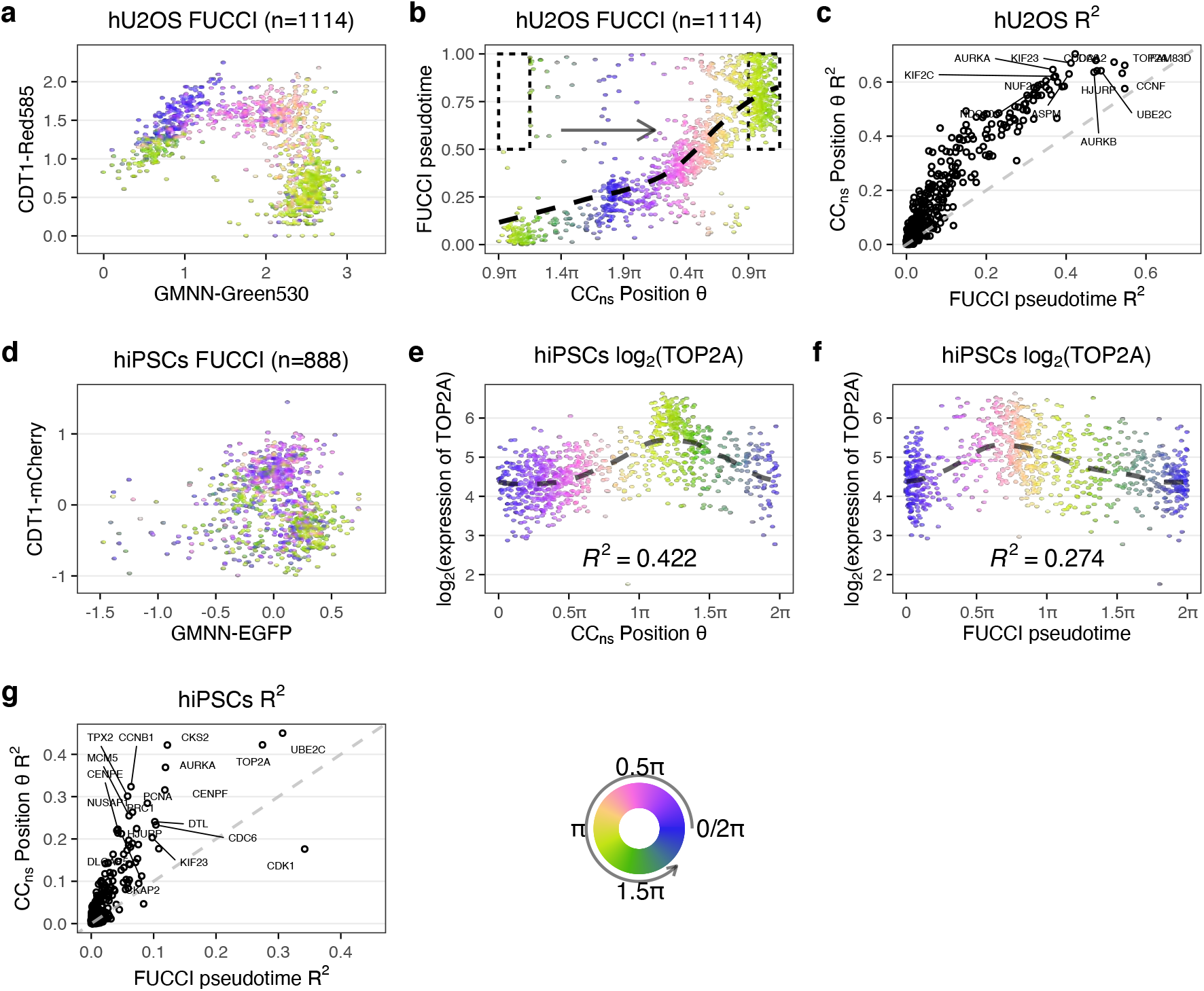
Evaluation of tricycle on FUCCI datasets. **(a-c)** Data from Mahdessian et al. (2021). **(a)** FUCCI scores colored by tricycle cell cycle position. **(b)** Comparison between FUCCI pseudotime and tricycle cell cycle position with a periodic loess line. We are displaying the data on [0.9*π*, 0.9*π* + 2*π*] (compared to [0, 2*π*] elsewhere) because FUCCI pseudotime of 0 roughly corresponds to a tricycle position of 0.9*π*. Cells in the dotted rectangle were moved for display purposes in this panel by adding 2*π* (one period) to tricycle *θ* to reflect the higher temporal resolution around the anaphase-metaphase transition for FUCCI pseudotime (see Results). **(c)** *R*^2^ values of periodic loess line of all projection genes when using tricycle *θ* and FUCCI pseudotime as the predictor. The dashed line represents *y* = *x*. **(d-g)** Data from Hsiao et al. (2020). **(d)** FUCCI scores colored by tricycle cell cycle position. **(e,f)** Expression dynamics of *Top2A* with a periodic loess line using either (e) tricycle cell cycle position or (f) FUCCI pseudotime inferred by Hsiao et al. (2020). Cells are colored by corresponding x-axis values. **(g)** Similar to (c), but for the data from Hsiao et al. (2020).

Hsiao et al. (2020) used FUCCI on human induced pluripotent stem cells (iPSC) followed by scRNA sequencing using Fluidigm C1. While the Mahdessian et al. (2021) FUCCI data look like a horseshoe, the Hsiao et al. (2020) FUCCI data are more akin to a cloud (the data differ in quantification and normalization of the FUCCI scores). These data are used to estimate a continuous cell cycle position (which we term “FUCCI pseudotime”) based on polar angle of the FUCCI scores. Compared with the data in Mahdessian et al. (2021), there are larger differences between FUCCI pseudotime and tricycle cell cycle position. However, we can directly compare the associated expression dynamics of key cell cycle genes (Figure 5 for TOP2A, Supplementary Figure S13 for 8 additional genes). These results suggests that tricycle cell cycle position is at least as good or better as the FUCCI pseudotime at ordering the cells along the cell cycle; the *R*^2^ for *TOP2A* is 0.42 for tricycle compared with 0.27 for peco.

In contrasts to FUCCI measurements, FACS sorting and enrichment of cells yields groups of genes in (supposedly) distinct phases of the cell cycle. We consider 2 different datasets where FACS has been combined with single-cell RNA-seq. Buettner et al. (2015) assays mouse embryonic stem cells (mESC) using Hoechst 33342-staining followed by cell isolation using the Fluidigm C1. They use very conservative gating for G1 and G2M at the cost of less conservative gating for S phase. Leng et al. (2015) uses FACS on FUCCI labeled H1 human embryonic stem cells (hESC) followed by cell isolation using the Fluidigm C1. In both experiments, cells largely appear as expected in the cell cycle embedding defined by the cortical neurosphere reference embedding (Supplementary Figure S14). For the mESC, we note that some cells labeled S (but not G1 or G2M) appear outside the position expected for this stage, consistent with the gating strategy used for these data.

Summarizing this evidence, we conclude that tricycle recapitulates and refines the cell cycle ordering consistent with current “state of the art” experimental methods. Tricycle cell cycle position is competitive with FUCCI based measurements, except for cells in the metaphase to anaphase transition during mitosis.

### Comparison with existing tools for cell cycle position inference

We next sought to compare tricycle cell cycle position estimates with those obtained from other available methods. Existing methods for cell cycle assessment can be divided into those which infer a continuous position and those which assign a discrete stage. We have evaluated the following methods: peco (Hsiao et al., 2020), Revelio (Schwabe et al., 2020), Oscope (Leng et al., 2015), reCAT (Liu et al., 2017), cyclone (Scialdone et al., 2015), Seurat (Stuart et al., 2019), the original Schwabe Schwabe et al., 2020, and the SchwabeCC 5 stage assignment method. Each method differs in which datasets it works well on and which issues it might have; a detailed comparison is available in the Supplement (Supplemental Methods, Supplementary Figures S15-S21).

Issues with existing methods include (a) the ability to work on datasets with multiple cell types, (b) the ability to scale to tens of thousands of cells or more, and (c) the ability to work on less information-rich datasets such as those generated by droplet-based or *in situ* scRNA-seq methods. Oscope requires data on many genes due to its use of pairwise correlations, and therefore does not work on less information rich platforms (e.g. 10x Chromium or Drop-Seq). peco works better on less sparse and information-rich data (e.g. Fluidigm C1), but even on data from this platform, it is outperformed by tricycle. reCAT is critically dependent on the extent to which a principal component analysis of the cell cycle genes reflects cell cycle and only infers a cell ordering; it is not straightforward to interpret the reCAT ordering, especially across datasets. Revelio is primarily a visualization tool, which appears to fail on datasets where substantial variation is driven by processes other than the cell cycle. Of the discrete predictors, Seurat agrees well with tricycle (and is very scalable) but is limited by only predicting a 3 stage cell cycle representation (G1/S/G2M). Cyclone appears to do poorly in labelling cells in S phase and only predicts 3 stages. The SchwabeCC predictor assigns 5 stages, but has many missing labels and mis-assigns cells from G0/G1 to other stages.

Additionally, we benchmarked the computational speed and performance of tricycle against other cell cycle estimation algorithms. We briefly compared the running time of several methods using subsets of the mRetina dataset (Supplementary Figure S22). To compute continuous estimates using tricycle takes a mean of about 0.58,0.86 and 1.48 seconds when the number of cells is 5000, 10000, and 50000 respectively. In contrast, to compute finite discrete stages, Seurat takes a mean of about 1.10, 1.22 and 4.95 seconds for a three-stage estimation and cyclone takes a mean of about 7.96, 11.50 and 50.66 minutes for a three-stage estimation, when the number of cells is 5000, 10000, and 50000 respectively. Other methods (peco, Oscope, reCAT) are not capable of processing large (10k-100k+) datasets. All of the comparisons were run on Apple Mac mini (2018) with 3.2 GHz 6-Core Intel Core i7 CPU, 64GB RAM, and operating system macOS 11.2. Thus, tricycle is able to scale with the increasing size of datasets.

### Application of tricycle to a single-cell RNA-seq atlas

To demonstrate the scalability and generalizability of tricycle we applied it to a recent dataset of ≈ 4 million cells from the developing human (Cao et al., 2020). The data were generated using combinatorial indexing (sci-RNA-seq3) and are relatively lightly sequenced with a median of 429 – 892 total UMIs for 4 single-cell profiled tissues and 354 – 795 for 11 single-nuclei profiled tissues (Supplementary Figure S23). Using tricycle, we are able to rapidly and robustly annotate cell cycle position for each of the cells/nuclei in this atlas (Figure 6a, Supplementary Figure S24). Within a global UMAP embedding, tricycle annotations enable immediate visual identification of proliferating and/or progenitor cell populations for most cell types and tissues. The rapid annotation of cell cycle position on this reference dataset further allowed us to examine the relative differences in the proportion of cells actively proliferating across different tissues and cell types in the developing human. To quantify this, we discretized all cells along *θ* into two bins corresponding to actively proliferating (0.25*π* < *θ* < 1.5*π*; S/G2/M) or non-proliferating (G1/G0). We next ranked each tissue by the relative proportion of actively proliferating cells to identify the tissues and cell types with the highest proliferative index (Figure 6b). To examine cell-type specific differences in proliferation potential, we computed the cell cycle embedding as well as the proliferative index for the 9 most abundant cell types within each tissue (Supplementary Figures S25 and S26).

**Figure 6.**
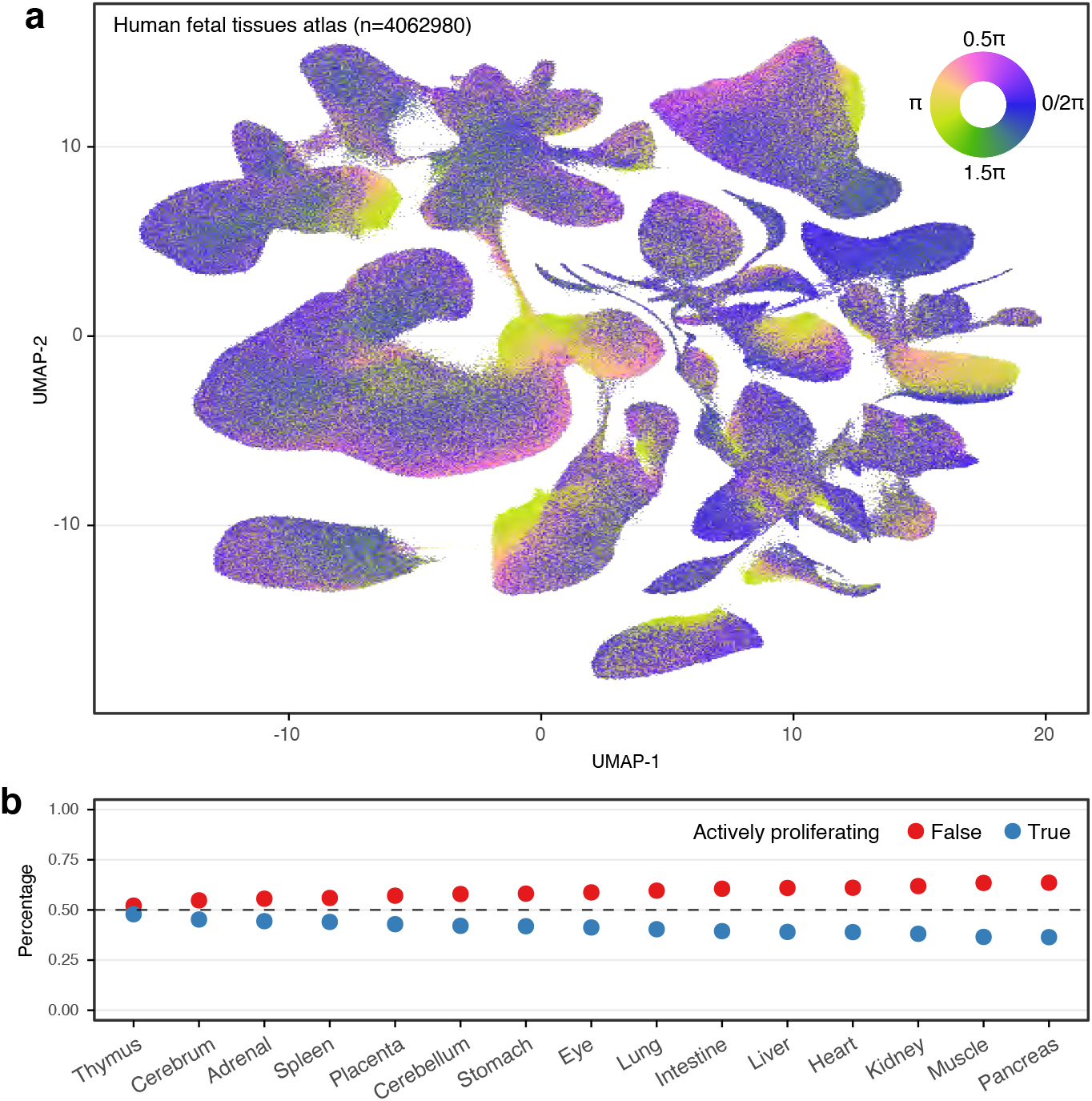
Application of tricycle on a human fetal tissue atlas. Data is from from Cao et al. (2020). **(a)** UMAP embedding of human fetal tissue atlas data colored by cell cycle position *θ* estimated using mNeurosphere reference. **(b)** The percentage of actively proliferating cells in human fetal tissue atlas. Tissues are ordered decreasingly with the percentage. Tissue and cell type annotations are available in Supplementary Figure S24.

Tissue-level proliferation indexes identified thymus, cerebrum, and adrenal gland as having the highest overall proportions of dividing cells across the sampled fetal timepoints. Within the thymus, thymocytes represent both the most abundant cell type and the most ‘prolific’ cell types as a function of the proportion of mitotic cells. Thymocytes exhibit a circular embedding in UMAP space that effectively recapitulates the estimated cell cycle position predictions from tricycle (Supplemental Figure S26k). Within this circular embedding, there is a gap of cells with cell cycle position estimates at ≈ *π*, consistent with dropout of cells and lower information content in M-phase. Comparison of tricycle cell cycle annotations to SchwabeCC cell cycle phase calls in this embedding suggests that tricycle more accurately estimates cell cycle position even on cell types with a mean total UMI of 354 (Supplementary Figure S27).

Within tissues, lymphoid cells are often the cell type with the highest proliferation index (Supplementary Figures S26, S25); often with a greater number of actively proliferating cells than not. Within the fetal liver and spleen – both sites of early embryonic erythropoiesis during human development (Cumano and Godin, 2007) – erythroblasts represent the cell type with the highest fraction of proliferating cells. Across developmental time, most tissues maintain relatively monotonic proliferation indices, however several (liver, placenta, intestine) exhibit dynamic changes across the sampled timepoints. This application illustrates the utility of tricycle to atlas-level data.

### Stability of the cell cycle position assignments

To test the robustness of tricycle we performed in silico experiments to determine the stability of cell cycle position assignments. We evaluated three different types of stability wrt. (a) missing genes, (b) sequencing depth, and (c) data preprocessing.

When projecting new data into the cell cycle reference embedding, it is common that the feature mapping between the two data sets contains only a subset of the 500 genes used in the embedding. The number of genes available for feature mapping has an impact on the shape of the resulting embedding; the mNeurosphere and mHippNPC datasets have almost the same shape when restricted to a set of common genes (Supplementary Figure S28). To establish the stability of tricycle, we randomly removed genes from the neurosphere dataset and computed tricycle cell cycle positions; we used the neurosphere dataset as a positive control to ensure all genes are present. We used the circular correlation coefficient to assess the similarity between the tricycle cell cycle position for the full dataset vs. the dataset with randomly pruned genes (Supplementary Figures S29, S30). This reveals excellent stability (circular *ρ* > 0.8) using as little as 100 genes.

To examine the impact of sequencing depth, we downsampled the mHippNPC dataset (Supplementary Figures S31, S32), and used the circular correlation coefficient to quantify to similarity to the cell cycle position inferred using the full sample. Originally, the median of library sizes (total UMIs) is 10,000 for mHippNPC data. Downsampling to 20% of the original depth(approximate median of library sizes 2,000) kept circular *ρ* > 0.8. This is congruent with the observed robustness of the method to the varying sequencing depth of the various datasets examined above.

Next, we examined the stability of tricycle wrt. the choice of reference embedding. Above, we show a cell cycle space estimated separately for the mNeurosphere and the mHippNPC datasets (Figure 2). We observe that the inferred expression dynamics are more alike in the two datasets if we project the mHippNPC into the mNeurosphere embedding compare to using its own embedding. To quantify this, we pick key cell cycle genes (previously examined in Supplementary Figure S6) and compare the location of peak expression in the mNeurosphere dataset to that of the mHippNPC dataset with cell cycle position estimated using these two approaches (Supplementary Figure S33). For the vast majority of genes, the highest expression appear at a closer position when we estimate cell cycle position by projecting the mHippNPC dataset into the mNeurosphere embedding.

To examine the impact of preprocessing data prior to projection, we compared cell cycle position inferred using data processed with and without Seurat. Note that when we estimate the cell cycle space, we use Seurat to align the different biological samples. But this is not done when we project new data using the prelearned reference. We observe negligible differences, whether or not Seurat is used (Supplementary Figure S34). We also conformed the direction of cell cycle position *θ* is consistent with the direction of RNA velocity projections (Supplementary Figure S35).

These results demonstrate the high sensitivity of tricycle to accurately estimate the cell cycle position across a high dynamic range of number of detectable genes within the feature map as well as depth of the information content in the target cells.

## DISCUSSION

Here, we have demonstrated the ability of tricycle to infer cell cycle position in 26 datasets across species, cell types, and assay technologies. To do so – as is common in the field – we have made extensive use of gold standard datasets, with a particular emphasis on the FUCCI assay. We show tricycle compares favorably to FUCCI-based pseudotime, specifically the tricycle inferred cell cycle position is a better predictor of expression dynamics of key cell cycle genes compared to FUCCI-based pseudotime; however, FUCCI pseudotime has higher temporal resolution during the metaphase to anaphase transition, a very specific point during cell division.

An important limitation of existing gold standard datasets is that the measurements are done on cell lines where the main driver of expression can be assumed to be cell cycle. In contrast, many common applications of single-cell expression contain multiple cell types (e.g. tissue samples) and/or other strong drivers of expression such as differentiation. Predicting cell cycle position in such datasets is much harder than predicting cell cycle position on homogeneous cell lines. For this reason, it is not enough to merely assess a method on gold standard datasets.

To address the limitation of gold-standard measurements on cell lines, we have made extensive use of internal controls. Specifically, we use inferred cell cycle position to assess whether key cell cycle genes (and log totalUMI if available) exhibit the expected expression dynamics across the learned *θ* progression. These internal controls are available in *any* single-cell expression dataset, including complex tissue samples. These internal controls do not by themselves give a clear answer to how precise the predictions are, but they undoubtedly carry some information on whether the inferred cell cycle position is at all associated with cell cycle phases. By using these internal controls, we overcome the limitations of the available gold standard datasets – which we can think of as having “external controls” – and show that tricycle performs well on differentiation datasets and datasets with multiple cell types. Additionally, we are able to ascertain the generalizability of the method. The cell cycle genes we use for evaluation are also used to construct the reference embedding and for projection. However, at the projection stage, the weights are fixed without any dataset dependent optimization. This removes – in our opinion – any circular reasoning. We note that internal controls are useful to assess any continuous prediction of cell cycle position.

Here, we use a fixed reference embedding to represent cell cycle, defined using the mouse cortical neurosphere dataset. This raises the question: is there a single best embedding? One part of this question is whether there is a minimal best set of genes to construct the embedding? Empirically, when we project the neurosphere data into itself and remove genes, we get good performance with around 100 genes. However, this experiment does not measure generalizability, and we have anecdotally observed that reducing our gene list this much impacts performance in some datasets. Another part of this question is whether we can optimize the embedding to be as circular as possible. We observe that, despite different shapes, embeddings based on the cortical neurosphere and the primary hippocampal NPC datasets result in similar cell cycle position estimates. One interpretation of these observations is that the robustness of the approach is derived from the structure created by the relationship of the genes to each other rather than the behavior of any individual marker gene, as described in Stein-O’Brien et al. (2019).

In many single-cell experiments, cell cycle is often considered a confounding factor and as such, methods exist to remove this effect from the data prior to analysis. We caution against *removing* cell cycle progression blindly as it can be intimately intertwined with other sources of variation of interest. Taking the mPancreas data as an example, there is a clear relationship between the number of cycling cells and differentiation as the multi-potent ductal cells advance to be terminally differentiated alpha and beta cells. If correction for cell cycle progression is warranted, our analysis of the mPancreas data suggests that the common approach of regressing out principal components of cell cycle genes may remove additional biological variation of interest.

It is currently unknown what will happen if tricycle is applied to a dataset without cycling cells. In this case, all cells belong to the G0/G1 cloud, and we hypothesize that the G0/G1 cloud will be centered on the (0, 0) origin and tricycle inferred cell cycle position will be wrong. We expect to be able to diagnose this situation using internal controls, but we caution against using tricycle or other cell cycle inference methods on a dataset without cycling cells.

Tricycle is a locked-down prediction procedure. There are no tuning parameters, neither explicitly set nor implicitly set through the use of crossvalidation or alternatives. And in our applications, we have not aligned different samples to each other. That being said, we observe that different datasets exhibit small discrepancies. An example is that the precise location of peak expression in Top2A differs slightly from dataset to dataset. Possible sources of dataset-to-dataset variation include both biological and technical candidates such as technology and batch effects.

There are two specific sources of variation we want to highlight. First is the variation in nonzero expression. Specifically, we observe that 300-500 genes – out of a total of 500 genes – have nonzero expression in a given dataset, and we show that differences in these genes can cause changes in the shape of the embedding (Supplementary Figures S28, S29, and S30). These differences in nonzero expression are also connected to the utility of using a larger set of genes to define the embedding space, as discussed above.

Second, the variation in the proportion of cells in G0 and G1, which is associated with the actual wall clock length of the cell cycle. Reflecting the biology of the system, this ratio also affects the placement of the origin of the projected data. However, this also potentially complicates across data set comparisons, as the only normalization currently performed in tricycle is the mean centering of each gene, which is susceptible to differences in this ratio. Lastly, we note that the peak expression location variations might be a consequence of different cell cycle dynamics in different systems. We hope the biological implications will be examined closely in the future. Methods to expand tricycle to allow cross data comparisons are currently an active area of research.

We anticipate that the ability to model cell cycle as the continuous process that it is, will enable considerable advancements in the modeling of developmental and disease processes in which it plays a major role.

## CONCLUSIONS

We have explained why the cell cycle – in datasets where the primary source of variation is cell cycle – is visible as an ellipsoid shape in the principal components of the data. We have shown that principal component analysis of cell cycle genes sometimes reflects other processes such as differentiation. We have proposed to use projections into a reference embedding to isolate the specific cell cycle signal in a dataset with many sources of variation, and we have shown that this approach allows us to isolate a specific, pre-specified signal.

We have shown that tricycle is capable of inferring continuous cell cycle position and can be applied to datasets with multiple cell types, across species and a variety of single-cell technologies from relatively deeply sequenced plate based technologies to shallower sequenced droplet-based technologies, including very sparse data. As part of applying tricycle, a user can use internal controls to assess the validity of the inferred cell cycle positions on their own dataset. Tricycle is highly scalable and available in an open-source implementation from the Bioconductor project.

## METHODS

### Using principal component analysis to recover time ordering

We will consider the following statistical model. The mean expression of each gene is modelled as

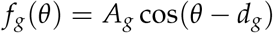

Here, *A_g_* is a gene-specific amplitude and *d_g_* is a mean-specific displacement (location of the peak). In this formulation, the mean function has a single peak and is periodic. We have *G* genes and each gene has its own (but not necessarily unique) (*A_g_, d_g_*).

Basic trigonometry yields the identity

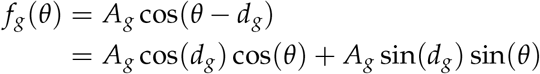

which we can write as

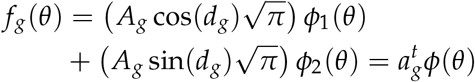

using the orthonormal functions

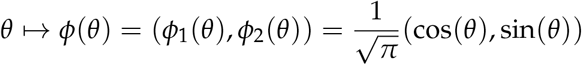

Our derivation is based on Ramsay and Silverman (2005) section 8.4. This section shows that the variance-covariance operator is given by

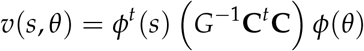

where the inner matrix (which turns out to determine the principal components) is a 2 × 2 matrix equal to

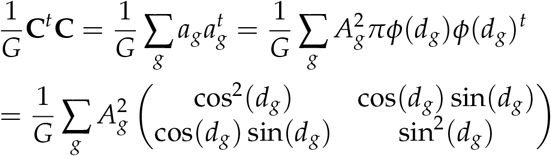

The principal component analysis is given by the Eigen-functions and -values of the variance-covariance operator. Such an Eigen-function and -value pair *ξ, ρ* takes the form

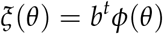

for a vector *b* which satisfies

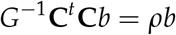

ie. *b, ρ* are Eigen-vectors and -values for the G^−1^ **C*^t^*C** matrix. Specifically, if *q*_1_, *q*_2_, *λ*_1_, *λ*_2_ are two such Eigen-vectors- and -values, then the two first principal components are given by

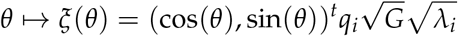

### Simulations

For Figure 1 we performed the following simulation. 50 realization of a cosine function with a location of 0.2 and an amplitude of 0.5 as well as 50 realizations of a cosine function with a location of 1.2 and an amplitude of 1. Each function was evaluated on an equidistant grid of 1000 points and independent Gaussian noise with a standard deviation of 0.2 was added. The depictions in Figure 1a,b were each one of the realizations of the two different cosine functions.

For Supplementary Figures S1, S2 and S3 we simulated data using the negative binomial distribution, inspired by the setup in Splatter (Zappia et al., 2017). In addition to a gene-specific amplitude (*A_g_*) and location of the peak (*L_g_*), we also consider different library size (*l*), which is an approximate as we still have some cell-to-cell variance. For a cell, we let 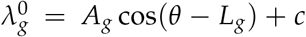, with *c* a constant to ensure positivity of 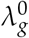. Then the cell mean is 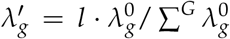. The trended cell mean is simulated from a Gamma distribution as 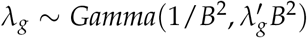, with *B* the biological coefficient of variation (we fix *B* as 0.1 in our simulations). Thus, the counts for gene *g* is given as *y_g_* ~ *Pois*(*λ_g_*). We always simulate a 100 genes times 5000 cells count matrix, with cell timepoint *θ* uniformed distributed between 0 and 2*π*. We only vary one of *L_g_*, *A_g_* and *l* in Supplementary Figures S1, S2 and S3. Specifically, in Supplementary Figures S1, we used different number of distinct peak locations across 100 genes, and fixed the amplitudes (across 100 genes) as 3 and library size as 2000. In Supplementary Figures S2, we used different numbers of distinct amplitudes across 100 genes, and fixed the number of distinct peak locations (across 100 genes) to 100 and library size to 2000. In Supplementary Figures S3, we changed the library size l, and fixed the number of distinct peak locations (across 100 genes) as 100 and the amplitudes (across 100 genes) as 3. PCA was performed on the library size normalized and log_2_ transformed matrix after we got the count matrix.

### Generation of mouse primary hippocampal NPC scRNA-seq dataset

Hippocampal neural stem/progenitor cells (NPCs) were isolated by microdissection from E17 day embryos (offspring of male *Kmt2d^+/βgeo^* and female C57Bl/6J) and cultured on Matrigel as described in Carosso et al. (2019). We verified neuronal lineage by demonstrating Nestin, Calbindin, and Prox1 expression (not shown). Cells were maintained in an undifferentiated state with growth factor inhibition (EGF, FGF2) in Neurobasal media. In a prior publication, we have demonstrated that the *Kmt2d^+/βgeo^* cells exhibit defects in proliferation (Carosso et al., 2019). Following isolation we collected cells from both genotypes at the undifferentiated state (day 0) and then after growth factor removal on days 4, 7, 10 and 14, capturing cells that were ever more differentiated. scRNA-seq libraries were created with a Chromium Single-Cell 3′ library & Gel Bead Kit v2 (10x Genomics) according to manufacturer protocol. Only cells from day 0 are analyzed here.

### Generation of mouse E14.5 Neurosphere scRNA-seq dataset

Cortical neurospheres were generated from the dissociated telencephalon of embryonic day 14.5 (E14.5) wild type embryos. Embryos were harvested and the dorsal telencephalon was dissected away and collected in 1X HBSS at RT temperature. The dorsal telencephalon was gently triturated using p1000 pipette tips and the resultant cell suspension was spun at 500G for 5min and the media was aspirated off. The cell pellets were resuspended in complete neurosphere media 7ml (CNM) and plated in ultra-low adherence T25 flasks. CNM is made by combining 480ml DMEM-F12 with glutamine, 1.45g of glucose, 1X N2 supplement, 1X B27 supplement without retinoic acid, 1x penicillium/streptomycin and 10ng/ml of both epidermal growth factor (EGF) and basic fibroblast growth factor (bFGF). The cell pellets were cultured for 3-5 days, or until spheroids have formed. The neurospheres were then collected and spun at 100G for 5min and the supernatant was removed. Neurospheres were resuspeneded in 5ml TrypLE and incubated for a maximum of 5min at 37° C with gentle trituration every 1.5min with a p1000 until the neurospheres are mostly a single-cell suspension. The cells were spun down at 500G for 5min and the supernatant was removed. The cells were resuspended in 15ml of CNM and gently passed through a 40uM filter to remove large cell clumps. The resultant cell suspension was then plated in T75 flasks for another 2-5 days or until spheres began to have dark centers. This process was repeated two more times before cells were collected for 10X Genomics single-cell library prep. Before single-cell library preparation, the neurospheres were dissociated as described above and passed through a 40uM filter to ensure a single-cell suspension. Approx. 7000 cells were selected from each sample for input to the scRNA-seq library prep. scRNA-seq libraries were created using the Chromium Single-Cell 3’ library & Gel Bead Kit v2 (10x Genomics) according to manufacturer protocol.

### Reference genome and mapping index building

For mouse, GRCm38 reference genome fasta file and primary gene annotation GTF file (v25) were downloaded from GENCODE (https://www.gencodegenes.org). Similarly, GRCh38 reference genome fasta file and primary gene annotation GFT file(v35) were downloaded for human. We built a reference index for use by alevin as described by Soneson (2020) using R package eisaR(v1.2.0), which we use to quantify both spliced and unspliced counts of annotated genes.

### scRNA-seq preprocessing

#### Mouse Neurosphere (mNeurosphere) dataset

fastqs files were used to quantify both spliced and unspliced counts by Alevin (Salmon v1.3.0) with default settings as described by Soneson (2020). Abundances matrices were read in by R package tximeta (v1.8.1). The spliced counts were treated as the expression counts. We removed cells with less than 200 expressed genes, and cells flagged as outliers (deviating more than triple median absolute deviations(MAD) from the median of log_2_(TotalUMIs), log_2_(number of expressed genes), percentage of mitochondrial gene counts, or log_10_(doublet scores)). system.

The doublet scores were computed using doubleto Cells function in R package scran (v1.18.1). All mitochondrial genes and any genes which were expressed in less than 20 cells were further excluded from all subsequent analyses. Expression abundances were then library size normalized and log_2_ transformed by function normalizeCounts in R package scuttle (v1.0.2). The biological samples were integrated by Seurat (v3.2.2). We then run PCA on the top 2000 highly variable genes of the integrated log_2_(expression) using the runPCA function with default parameters, followed by running the runUMAP function on the resulting top 30 principal components with default parameters. Note that we did not restrict genes to cell cycle genes in this step, as we would like to see the overall variation of the data. Cell types were inferred by SingleR package v1.4.0 using built-in MouseRNAseqData dataset as the reference.

#### Mouse primary hippocampal NPC (mHipp-NPC) dataset

All preprocessing are the same as for the mouse Neurosphere (mNeurosphere) dataset.

#### Mouse developing pancreas (mPancreas) dataset

We obtained the spliced and unspliced count matrices of the Mouse developing pancreas dataset from the python package scvelo (v0.2.1). The spliced counts were treated as the expression counts. We removed cells with less than 200 expressed genes, and any cells flagged as outliers (deviating more than triple median absolute deviations(MAD) from the median of log_2_(TotalUMIs), log_2_(number of expressed genes), percentage of mitochondrial gene counts, or log_10_(doublet scores)). Here, the doublet scores were computed using doubletCells function in R package scran (v1.18.1). All mitochondrial genes and any genes which were expressed in less than 20 cells were further excluded from all subsequent analyses. Expression abundances were then library size normalized and log_2_ transformed by function normalizeCounts in R package scuttle (v1.0.2). We run PCA on the top 500 highly variable genes using the runPCA function with default parameters, followed by running the runUMAP function on the resulting top 30 principal components. When running the UMAP, we set *min_dist* to 0.5 instead of default value 0.01 to replicate the UMAP figure shown in Bergen et al. (2020) with other parameters default. Of note, the single-cell libraries of the data was generated using 10x Genomics’ Chromium v2

#### Mouse Hematopoietic Stem Cell (mHSC) dataset

We downloaded processed log_2_ transform TPM matrix directly from GEO under accession number GSE59114 (Kowalczyk et al., 2015). We only used the cells from C57BL/6 strain, of which contains more cells, as the number of overlapped genes between xlsx file of C57BL/6 strain and DBA/2 strain is too small. Because the data was already processed and filtered, we did not perform any other processing. Unlike the above mentioned dataset, the SMARTer protocol was applied during library preparation.

#### Mouse Retina (mRetina) dataset

This dataset is available at https://github.com/gofflab/developing_mouse_retina_scRNASeq. We removed cells flagged as outliers (deviating more than triple median absolute deviations(MAD) from the median of log_2_(TotalUMIs), log_2_ (number of expressed genes), percentage of mitochondrial gene counts, or log_10_(doublet scores)). As the total UMIs depend on cell type, we filtered the cells by blocking for each cell type. The doublet scores were computed using doubletCells function in R package scran (v1.18.1). All mitochondrial genes and any genes which were expressed in less than 20 cells were further excluded from all subsequent analyses. Expression abundances were then library size normalized and *log*_2_ transformed by function normalizeCounts. We used the cell type annotations as the *new_CellType* column in the provided phenotype file. The single-cell libraries of the data was generated using 10x Genomics’ Chromium v2 system.

#### HeLa cell lines datasets

The spliced and unspliced count matrices of HeLa Set 1 (HeLa1) and HeLa Set 2 (HeLa2) were downloaded from GEO website with accession number GSE142277 and GSE142356. Both datasets were generated by the same lab under the same protocol, while the sequencing depth of Set 2 is only about half that of Set 1 (Schwabe et al., 2020). For each dataset, we only used the genes existing in both spliced and unspliced count matrices. The spliced counts were treated as the expression counts. We removed cells with less than 200 expressed genes, and cells flagged as outliers (deviating more than triple median absolute deviations(MAD) from the median of log_2_(TotalUMIs), log_2_ (number of expressed genes), percentage of mitochondrial gene counts, or log_10_(doublet scores)). All mitochondrial genes and any genes which were expressed in less than 20 cells were further excluded from all subsequent analyses. Expression abundances were then library size normalized and log_2_ transformed by the function normalizeCounts. The single-cell libraries of the data were generated using Drop-seq system.

#### Mouse embryonic stem cell (mESC) dataset

The processed count matrix was downloaded from ArrayExpress website under accession number E-MTAB-2805 (https://www.ebi.ac.uk/arrayexpress/experiments/E-MTAB-2805/). We only retained 279 cells with *log*_2_(counts) greater than 15. The count matrix was library size normalized across cells and *log*_2_ transformed by function normalizeCounts. The RNA-seq data was generated using Fluidigm C1 system in this dataset.

#### Human embroyonic stem cells (hESC) dataset

The processed count matrix was downloaded from GEO under accession number GSE64016. We only retained FACS sorted cells. The count matrix were library size normalized across cells and log_2_ transformed by function normalizeCounts. The RNA-seq data was generated using Fluidigm C1 system in this dataset.

#### Human U-2 OS cells (hU2OS) dataset

The TPM matrix was downloaded from GEO under accession number GSE146773. We only retained FACS sorted cells with log_2_(counts) greater than the 3 times MAD range. Genes which were expressed in less than 20 cell were removed. The left TPM matrix were library size normalized across cells and log_2_ transformed by function normalizeCounts. The RNA-seq data was generated using SMART-seq2 chemistry in this dataset. We got the FUCCI coordinates and FUCCI pseudotime directly from the authors of Mahdessian et al. (2021) (version 1.2).

#### Human induced pluripotent stem cells (hiP-SCs) dataset

The processed FUCCI intensity and RNA-seq data was downloaded from https://github.com/jdblischak/fucci-seq/blob/master/data/eset-final.rds?raw=true. The preprocessing was described in Hsiao et al. (2020). The count matrix were library size normalized across cells and log_2_ transformed by function normalizeCounts. The RNA-seq data was generated using Fluidigm C1 system in this dataset.

#### Fetal tissue dataset

We got the loom file containing gene counts of all tissue from GEO under accession number GSE156793. We then processed and analyzed each tissue separately. For each tissue type, cells of which log_2_(TotalUMIs) is lower than median – 3 × MAD, and genes expressed in less than 20 cells were excluded from further analyses. The count matrix was library size normalized across cells and log_2_ transformed by function normalizeCounts. All 4 tissues profiled using singlecell and 9 tissues profiled using single-nuclei were generated on sci-RNA-seq3 system.

### 5 stage cell cycle assignments

The 5 stage (G1S, S, G2, G2M, and MG1) cell cycle assignments were adapted from Schwabe et al. (2020) with some modifications. Briefly, the assignments use the high expression genes list for each stage, curated by Whitfield et al. (2002). Let *k* represent one of the 5 stages, and 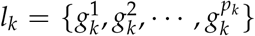 represent the gene list with *p_k_* genes. For each stage *k*, we could calculate the mean expression across genes in the gene list *l_k_* for the *j*th cell as 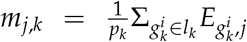 with 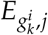 as the log_2_ transformed expression value of gene 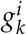 and cell *j*. Then we assess how well a gene in a gene list correlates to the mean expression level of that gene list as 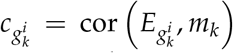. For each stage, the gene list is pruned to genes with 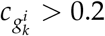. (For the fetal tissues dataset, we used 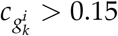 since the extremely shallowly sequenced data shows less co-expression patterns and the threshold 0.2 could leave us with no genes.) We label this pruned new gene list as 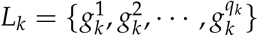 with *q_k_* the number of genes. The stage assignment score for cell *j* and stage *k* is given as

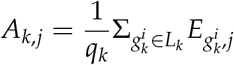

The 5-by-n matrix **A**, of which the number of columns equals to the number of cells, follows *z*-score transformations w.r.t. first rows and then columns, resulting the 5-by-n matrix 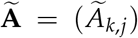. For each cell, we compute the preliminary stage assignment as 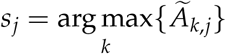.

As in the Schwabe et al. (2020), we also apply two filtering steps. The first filtering, which is unchanged from the original method, is as follows. We require 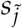, the stage with the second largest assignment score, to be the neighboring stage to *s_j_*. This requirement corresponds to the 5 stages being continuously cyclic processes.

As for the second filtering step, the original method discards all cells with the second largest assignment score 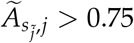. We found the threshold of 0.75 to some extent not applicable, as in some datasets it leads to losing 90% of cells. Therefore, we use a more adaptive threshold by requiring 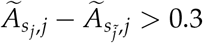.

If the cell passes two filtering steps, it will be assigned to a stage *s_j_*. Otherwise, it would be assigned as *NA* w.r.t. 5 stages of the cell cycle. To mitigate the batch effect on the 5 stage assignments, the assigning procedures are done for each sample/batch separately within each dataset, as recommended in Revelio package (Schwabe et al., 2020).

### PCA of GO cell cycle genes

For each dataset, we subsetted the preprocessed log_2_ transformed expression matrix to genes in the GO term cell cycle (GO:0007049). If there are clear batches defined in the dataset, such as sample or batch, we use Seurat3 to remove batch effect. In the case of using Seurat3, we used a library size normalized count matrix as input instead of log_2_ transformed values. The integration anchors were searched in the space of the top 30 PCs. The output integrated matrix is a log_2_ transformed matrix of the top 500 most variable genes. We then performed principal component analysis on the genewise mean centered expression matrix. In the case of no batch exiting, we also restricting to the top 500 variable genes among GO cell cycle genes.

### Projection of new data to cell cycle embedding and calculation of cell cycle position *θ*

The projection using pre-learned weights matrix during PCA of GO cell cycle genes is straight forward, given by

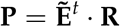

where **R** represents the o-by-2 reference matrix (*o* ≤ 500), contains the weights of top 2 PCs learned from PCA of GO cell cycle genes; 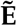 is a o-by-n matrix, subsetted from **E** (the log_2_ transformed expression matrix) with genes in the weights matrix and rowmeans centered. The resulting n-by-2 **P** is the cell cycle embedding projected by the reference. The calculation of the cell cycle position *θ* is given by

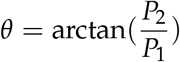

where *P_i_* is the *i*th column of matrix **P**. When mapping the genes between weights matrix and the data that we want to project, the Ensemble ID is given higher priority than the gene symbol for mouse. For across species projection, we only consider the homologous genes of the same gene symbols.

### Periodic loess

As *θ* is a circular variable bound between 0 to 2*π*, fitting a traditional loess model *y* ~ *θ*, with *y* as any response variable, such as the gene expression of gene, or log_2_(TotalUMIs), has problems around the boundaries 0 and 2*π*. Hence, we concatenate triple *y* and triple *θ* with one period shift to form [*y, y, y*] and [*θ* – 2*π, θ, θ* + 2*π*], on which the loess line is fitted. We then only use the fitted value 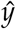 when *θ* is between 0 and 2*π* for visualization purpose.

The calculation of the coefficient of determination *R*^2^ of the fitted loess model is given by

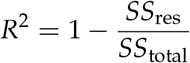

Here 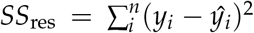 and 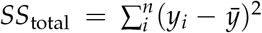. Note that instead of using all three copies of data points, we restrict the calculation of *SS*_res_ and *SS*_total_ on the original data points (the middle copy) The residuals are not the same for the three copies, especially at the beginning and end of [−2*π*, 2*π*].

### The circular correlation coefficient *ρ*

We use the circular correlation coefficient *ρ* defined by Jammalamadaka and Sarma, 1988 to evaluate concordance between two polar vectors *θ*_1_ and *θ*_2_ It is defined as follows

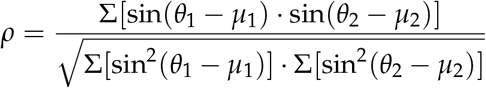

*μ*_1_ and *μ*_2_ represent the mean of *θ*_1_ and *θ*_2_ respectively, and are estimated by maximum likelihood estimation under von Mises distribution assumption.

### Running other methods

For other cell cycle inference methods, we use all default parameters and its built-in reference (if needed) in the following packages: cyclone in scran (v1.18.5), CellCycleScoring in Seurat (v4.0.0.9015), Revelio (v0.1.0), peco (v1.1.21), and reCAT (v1.1.0).

### Silhouette index on angular separation distance of tricycle cell cycle position *θ*

For cyclone and Seurat, we could use Silhouette index to describe consistency between discretized cell cycle stage and tricycle cell cycle position *θ*. We use angular separation distance metric to quantify the distance between cell *i* and cell *j* as

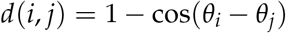

For a cell *i* ∈ *S_k^(i)^_* ∋ *k^(i)^* ∈ {G1, S, G2*M*}. The mean distance between cell *i* and all other cells assigned to the same stage

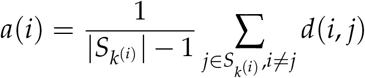

with |*S_k^(i)^_*| the cardinality of *S_k^(i)^_*. Specially, *a*(*i*) = 0 if |*S_k^(i)^_*| = 1. The mean distance from cell *i* to all cells assigned to other stage *k′* such that *k′* ≠ *k^(i)^* Λ *k′* ∈ {G1, S,G2M} is

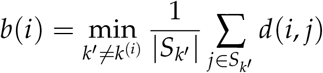

The Silhouette index for cell *i* is given as

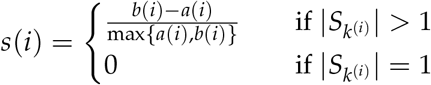

For any cell *i*, the Silhouette index *s*(*i*) is bound between −1 to 1 (−1 ≤ *s*(*i*) ≤ 1). An *s*(*i*) close to 1 means the cell is consistently assigned to its neighbors w.r.t. its cell cycle position *θ_i_*. An *s*(*i*) close to −1 means the cell is closer to the other stage. An *s*(*i*) equals to 0 means the cell is on the border of two stages. The mean Silhouette index on all cells measures how tight the stage assignments are. In this context, this value must be interpreted carefully as it is different from traditional clustering which might put hard boundaries and gaps between clusters. As the cell cycle process is continuous in nature, there must be cells assigned on the boundaries and ambiguous to either stage, and no gap should appear between stages. Thus, the mean silhouette index greater than 0 might be appropriate to conclude the agreement between tricycle cell cycle position *θ* and discretized cell cycle stages.

## Data and code availability

The mouse E14.5 Neurosphere data is available under GEO accession number GSE171636. The mouse primary hippocampal NPC data is being submitted to NCBI GEO. Get in touch if you want it sooner!

All code to analyze the data and generate figures is available at https://github.com/hansenlab/tricycle_paper_figs. The processed data is available at https://doi.org/10.5281/zenodo.5519841.

## Software availability

The tricycle method is implemented in the R package tricycle containing the mNeurosphere reference, which is available on https://github.com/hansenlab/tricycle and on Bioconductor (https://www.bioconductor.org/packages/tricycle).

## Disclaimer

The content is solely the responsibility of the authors and does not necessarily represent the official views of the National Institutes of Health or National Science Foundation.

## SUPPLEMENTARY MATERIALS

### SUPPLEMENTARY METHODS

#### Comparison with existing cell cycle tools

##### Oscope

Oscope poses significant challenges when run on shallow data (10X, sci-RNA-seq3, or DropSeq), since the method requires quantification of a high number of genes in every cell. For this reason, we do not evaluate Oscope.

##### peco

Peco supplies 2 models: one trained on 101 genes and one trained on 5 genes. We used the 101 gene model to be robust to some genes not being measurable in all datasets. We applied peco to all dataset described in Table 1, except mRetina and human fetal tissues. For human fetal tissues, we only use a subset of 2000 random cells selected from human fetal intestine data (termed “hfIntestineSub”).

We assess the expression dynamics of 4 genes highlighted in Hsiao et al. (2020): *CDK1, TOP2A, UBE2C* and *H4C3* (Supplementary Figure S15); not all datasets have these genes measured in which case they are absent from the figure. To systematically compare tricycle and peco we use the *R^2^* associated with two different cell cycle positions. This is a comparison between *R*^2^ for the same data, but using the same periodic loess approach with two different position variables. For these genes, across all dataset, tricycle cell cycle position has a higher *R*^2^ than peco cell cycle position (Supplementary Figure S15). Generally, information-rich Fluidigm C1 data does better with peco compared to information-poor 10X, Drop-Seq.

##### Revelio

Revelio is designed to search for an ellipsoid pattern amongst (rotated) principal components, by finding the directions having the strongest association with 5 discrete cell cycle stages. The output of Revelio is therefore supposed to be an ellipsoid. Revelio by itself does not quantify cell cycle position, although it seems natural to do so by the angle. When we use Revelio, we do indeed observe an ellipsoid in 4 datasets (Supplementary Figure S16a, b, f, g, i and j), but it clearly fails in 3 datasets: mPancreas dataset, mRetina dataset, and mHSC dataset (Supplementary Figure S16c, d, and e). These 3 datasets all have substantial variation which is not associated with cell cycle, such as cell types and differentiation, which we believe explains the non-ellipsoidal embedding. For example, in the mPancreas data, some of the differentiation effect is perfectly confounded with cell cycle as the terminally differentiated cells stop cycling. It is not clear that simply rotating the principal components will help us find a better cell cycle exclusive dimension. Additionally, Revelio removes any cell which does not have a prediction using the Schwabe stage predictor; in the mRetina dataset only 30k out of more than 90k cells are retained.

##### reCAT

reCAT starts with a principal component analysis of the cell cycle genes, and infers an ordering by solving a traveling salesman problem on this representation. This produces an ordering, but this ordering is hard to interpret because it is not directly linked to cell cycle stage. To address this, the authors provide two different stage predictors. Because the method requires the solution of a traveling salesman problem, it scales poorly. Due to these issues, we only ran reCAT on data with less than 5000 cells. The orderings inferred by reCAT are largely consistent with our cell cycle position *θ* using mNeurosphere reference for all datasets except the most shallow sequenced hfIntestineSub data (Supplementary Figure S17 last sub-panel in each panel). And the expression dynamics of Top2A on the time series also confirms the appropriate ordering of cells (Supplementary Figure S17 the third subpanel in each panel). However, the two-stage predictors given by reCAT yield different predictions on stages. For example, for the mPancreas dataset (Supplementary Figure S17a), the majority of cells are at S stage based on Bayes scores but are at G1 stage based on mean scores. Note that the reCAT function requires the user to feed an approximate cutoff position to assign a cell cycle stage based on Bayes scores. However, in all the datasets, we are unable to assign cutoff position to let each stage have its own highest scores interval. Without a useful stage assignment, the ability to make use of the cell orders is substantially restricted as the percentage of each stage is different across dataset.

##### Cyclone

We observe a general agreement between the 3 stage predictions of cyclone and tricycle cell cycle position, as the cyclone stages cluster together (Supplementary Figure S18). We note that cyclone assigns very few cells to the S stage. We believe this is caused by the assignment strategy (cells are assigned to S stage if both G1 and G2M scores are below 0.5). To expand on this comparison, we computed silhouette index with a distance defined by the tricycle cell cycle position (Methods). For cyclone, the underrepresentation of S stage drags down the silhouette index for both G1 and S stages, as cells at S stages are usually mixed with G1 cells, making the mean distance to all cells at G1 stage and to all cells at S stage not that differentiable. We note that cyclone works best on the last two FACS datasets, with one of them (mESC) being the training dataset for cyclone gene list.

##### Seurat

We observe good agreement between the 3 stage predictions of Seurat and tricycle cell cycle position, better than cyclone (Supplementary Figure S19). Compared to cyclone, we have a much higher silhouette index for Seurat; the highest observed mean is 0.74 for the mHSC dataset, which confirms the highly visual agreement between Seurat assignments and tricycle. The main disadvantage of Seurat is the inherent limitation of a 3 stage prediction.

##### SchwabeCC

The SchwabeCC method assigns cells to 5 different stages. Because of the higher resolution, it is the main predictor we use in our work. By default, the Schwabe method as reported in Schwabe et al. (2020) produces a substantial amount of missing labels, and we have therefore modified the method to address this (Methods); we used this modified Schwabe predictor unless specified otherwise (named as SchwabeCC).

Broadly, the SchwabeCC predictor agrees with tricycle, with one specific type of disagreement. These inconsistencies are examined in Supplementary Figure S20. Some cells with a tricycle cell cycle position of 0/2*π* (G0/G1) are assigned to other stages by SchwabeCC (Supplementary Figure S20 second subpanel of each row). It is well appreciated that there are many more genes specifically expressed at S, G2 or M stage as compared to G0/G1 stage (Dolatabadi et al., 2017). For each dataset, we plot out the percentage of nonexpressed genes over all projection genes in the first sub-panels, which show that the dynamics of percentages are captured by cell cycle position *θ* using mNeurosphere reference. We plot the percentage of non-expressed genes conditioned on stage and whether tricycle cell cycle position is around 0/2*π* (Supplementary Figure S20 third sub-panel of each row), which confirms that for each stage there exist two distinct groups. This is reinforced by the different expression patterns of *Top2A* and *Smc4* between flagged cells and non-flagged cells in the last two sub-panels. Thus, we conclude the cells around 0/2*π* are likely to be wrongly assigned to other stages, probably due to low information content.

To assess whether these inconsistencies are caused by our modification of Schwabe, we repeat the comparison using the original Schwabe assignments and arrive at the same conclusion (Supplementary Figure S21). This assessment highlights the large number of missing labels from the original Schwabe predictor, for example only 30k out of 90k cells in the mRetina dataset are labelled.

### SUPPLEMENTARY FIGURES

**Supplementary Figure S1.**
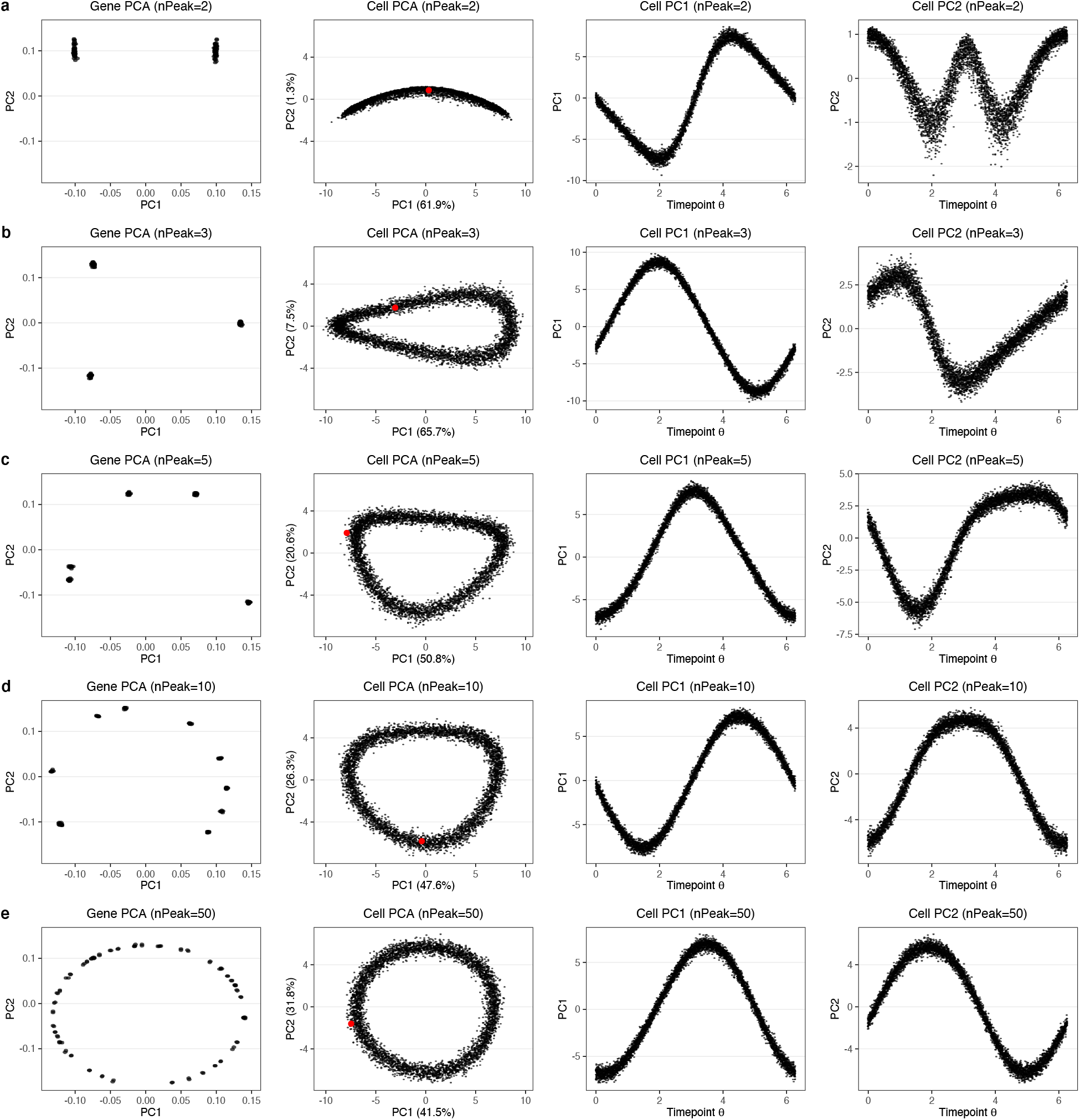
Simulations using negative binomial distribution with different number of distinct peak locations. We used different number of distinct peak locations across 100 genes, and fixed the amplitudes (across 100 genes) as 3 and library size as 2000. The number of distinct peak locations across 100 genes is **(a)** 2, **(b)** 3, **(c)** 5, **(d)** 10, and **(e)** 50. As long as we have more than 2 distinct peak locations, we get an ellipsoid.

**Supplementary Figure S2.**
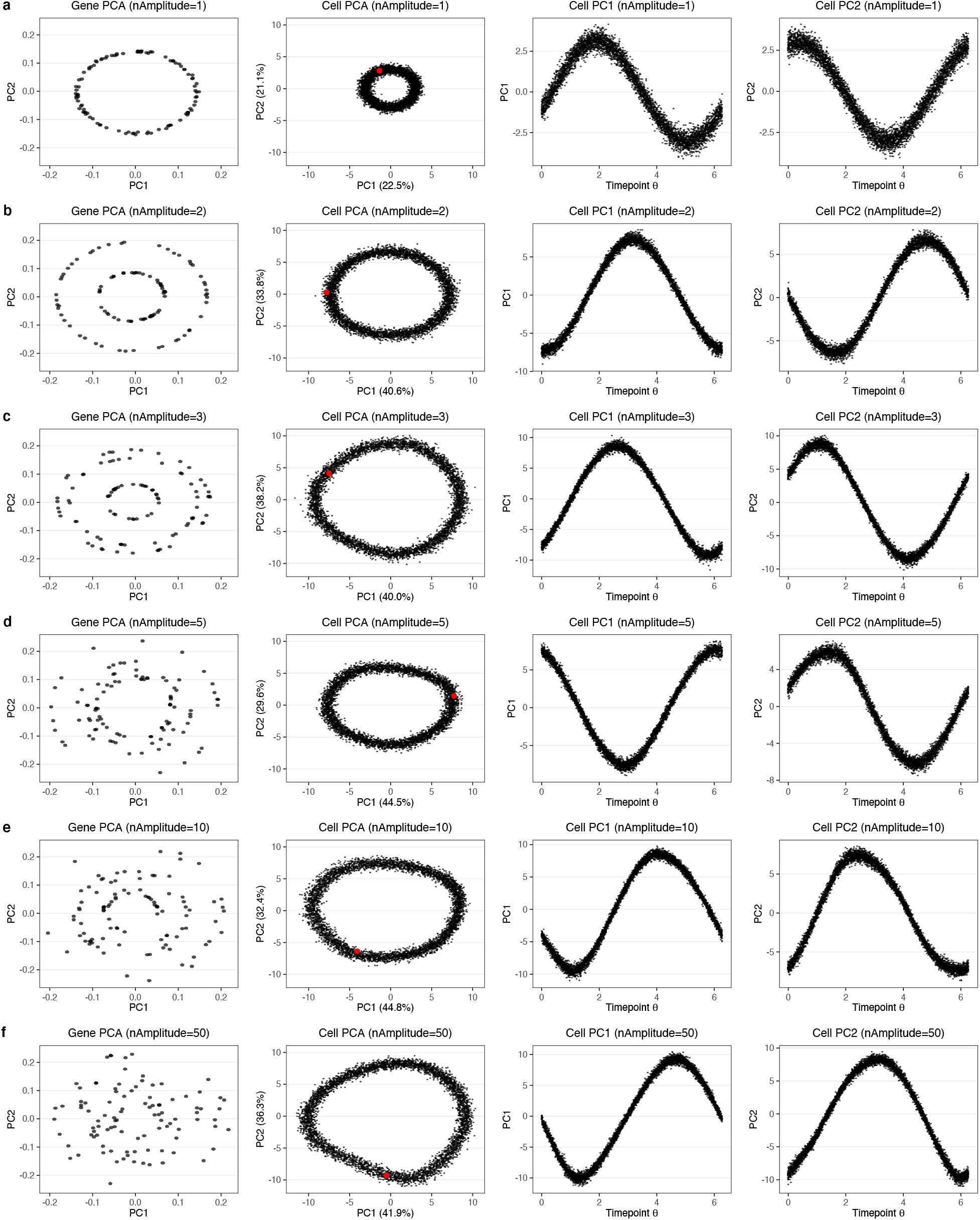
Simulations using negative binomial distribution with different number of distinct amplitudes. We used different numbers of distinct amplitudes across 100 genes, and fixed the number of distinct peak locations (across 100 genes) as 100 and library size as 2000. The number of distinct amplitudes across 100 genes is **(a)** 1, **(b)** 2, **(c)** 3, **(d)** 5, **(e)** 10, and **(f)** 50. No matter what the number of distinct amplitude(s) is, we always get an ellipsoid.

**Supplementary Figure S3.**
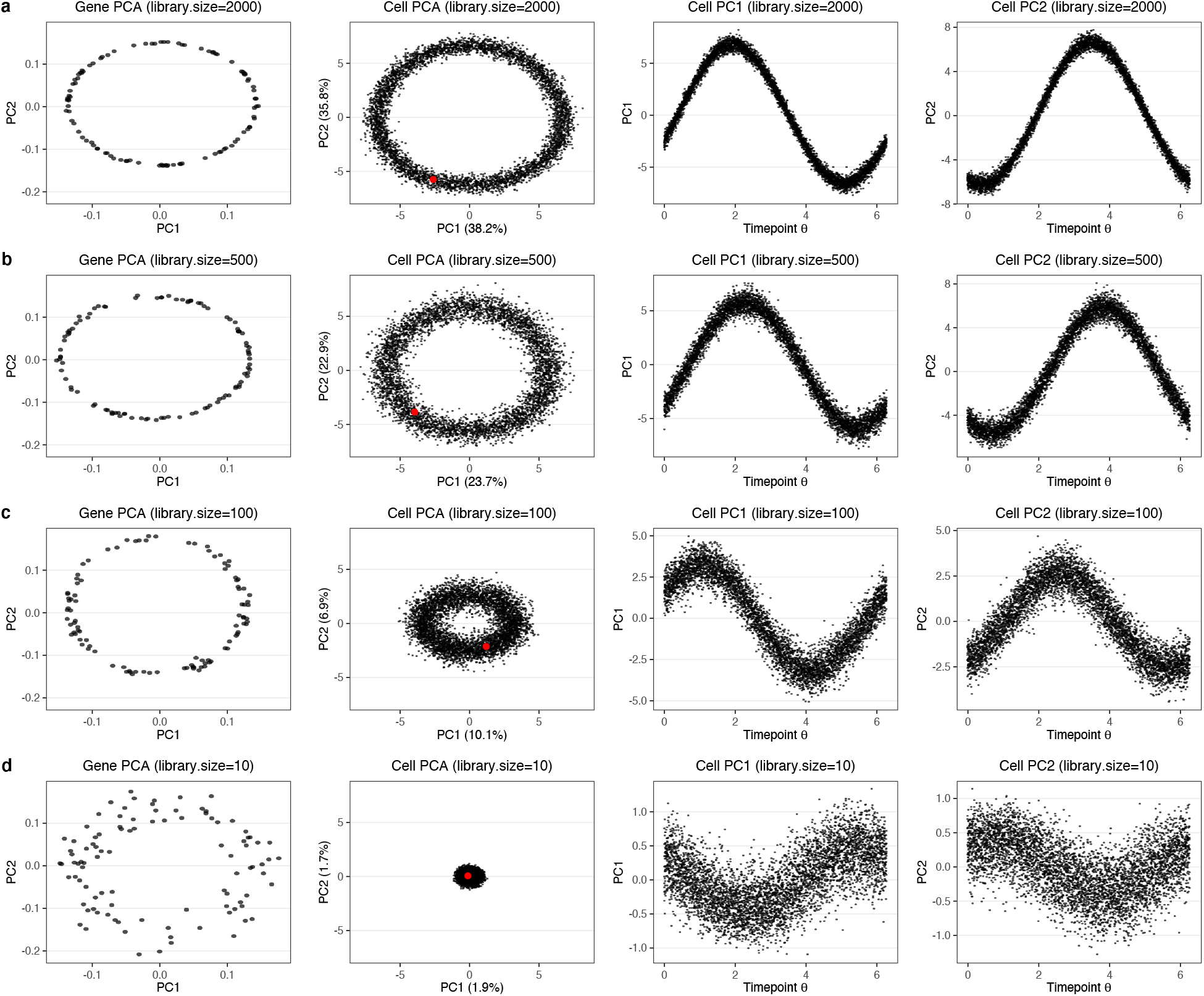
Simulations using negative binomial distribution with different library size. We changed the library size l, and fixed the number of distinct peak locations (across 100 genes) as 100 and the amplitudes (across 100 genes) as 3. The library size is **(a)** 2000, **(b)** 500, **(c)** 100, and **(d)** 10. The range of x-axis and y-axis of the first two sub-panels are fixed across (a)-(d). With library size decreasing, the ellipsoid shrinks to the (0, 0). However, the orders of cell can still be recovered.

**Supplementary Figure S4.**
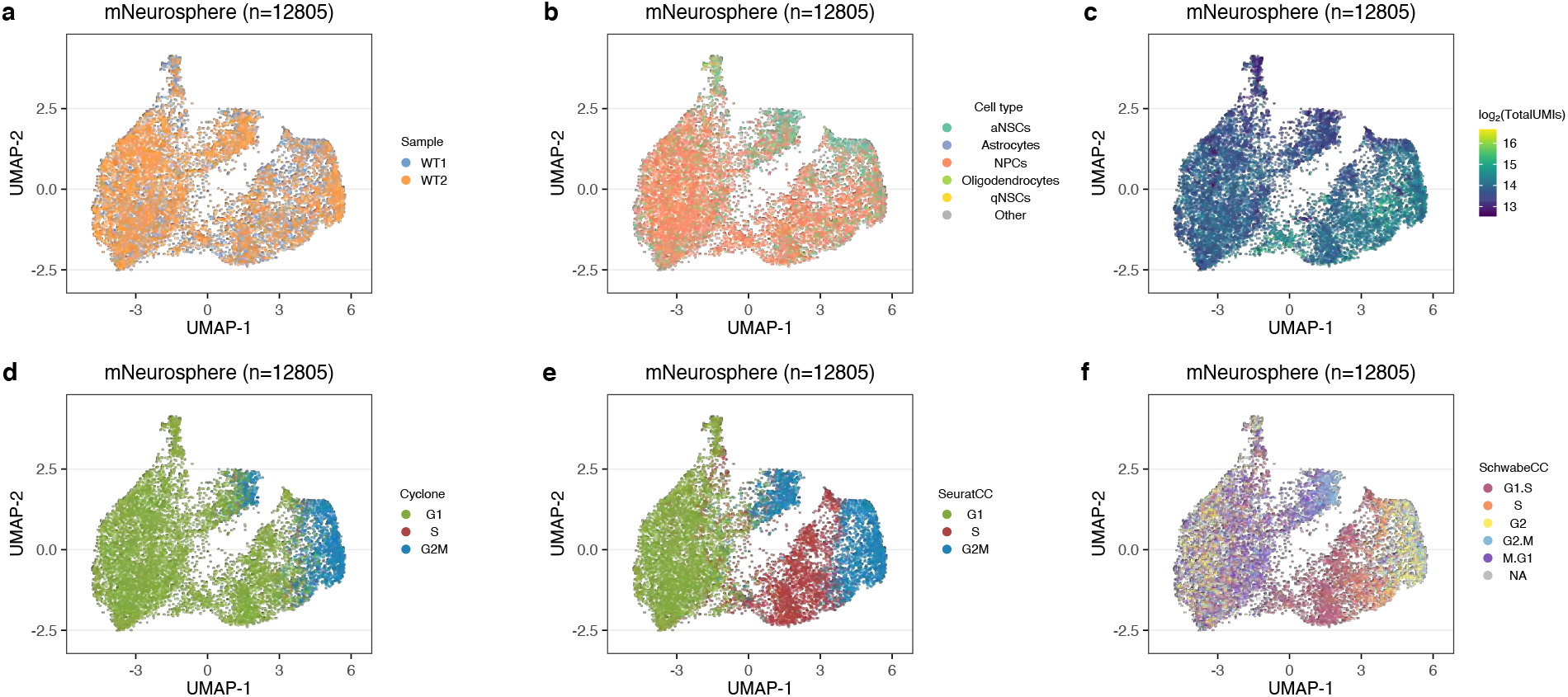
UMAPs of the mouse cortical Neurosphere dataset. Scatter plots show the UMAPs of Seurat3 mergerd Neurosphere data colored by **(a)** sample, **(b)** cell type inferred by SingleR, **(c)** log_2_(TotalUMIs), **(d)** inferred cell cycle stage by cyclone, **(e)** inferred cell cycle stage by Seurat, **(f)** inferred cell cycle stage by the SchwabeCC method (Schwabe et al., 2020) (See Methods). The UMAP coordinates were computed using the PCA on top 2000 highly variable genes after integreation by Seurat3.

**Supplementary Figure S5.**
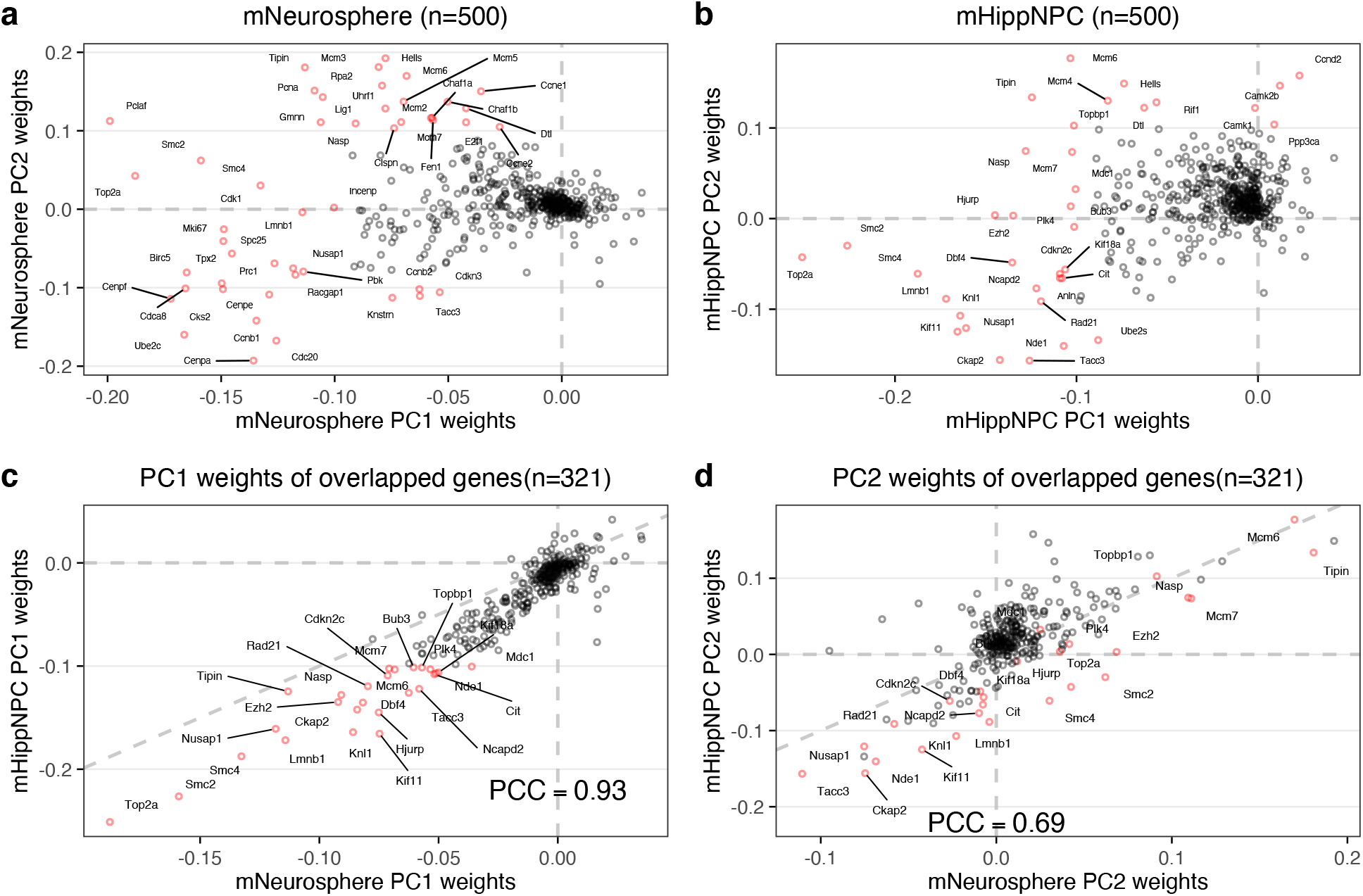
Weights of PCA on GO cell cycle genes. **(a)** The weights of top 2 PCs learned from doing PCA on GO cell cycle genes of cortical Neurosphere data. **(b)** The weights of top 2 PCs learned from doing PCA on GO cell cycle genes of mouse primary hippocampal NPC data. **(c)** A comparison of the weights on principal component 1 between the cortical neurosphere and hippocampal progenitor datasets. **(d)** As (c), but for PC2. Genes with high weights (|score| > 0.1 for either vector) are highlighted in red. PCC: Pearson’s Correlation Coefficient.

**Supplementary Figure S6.**
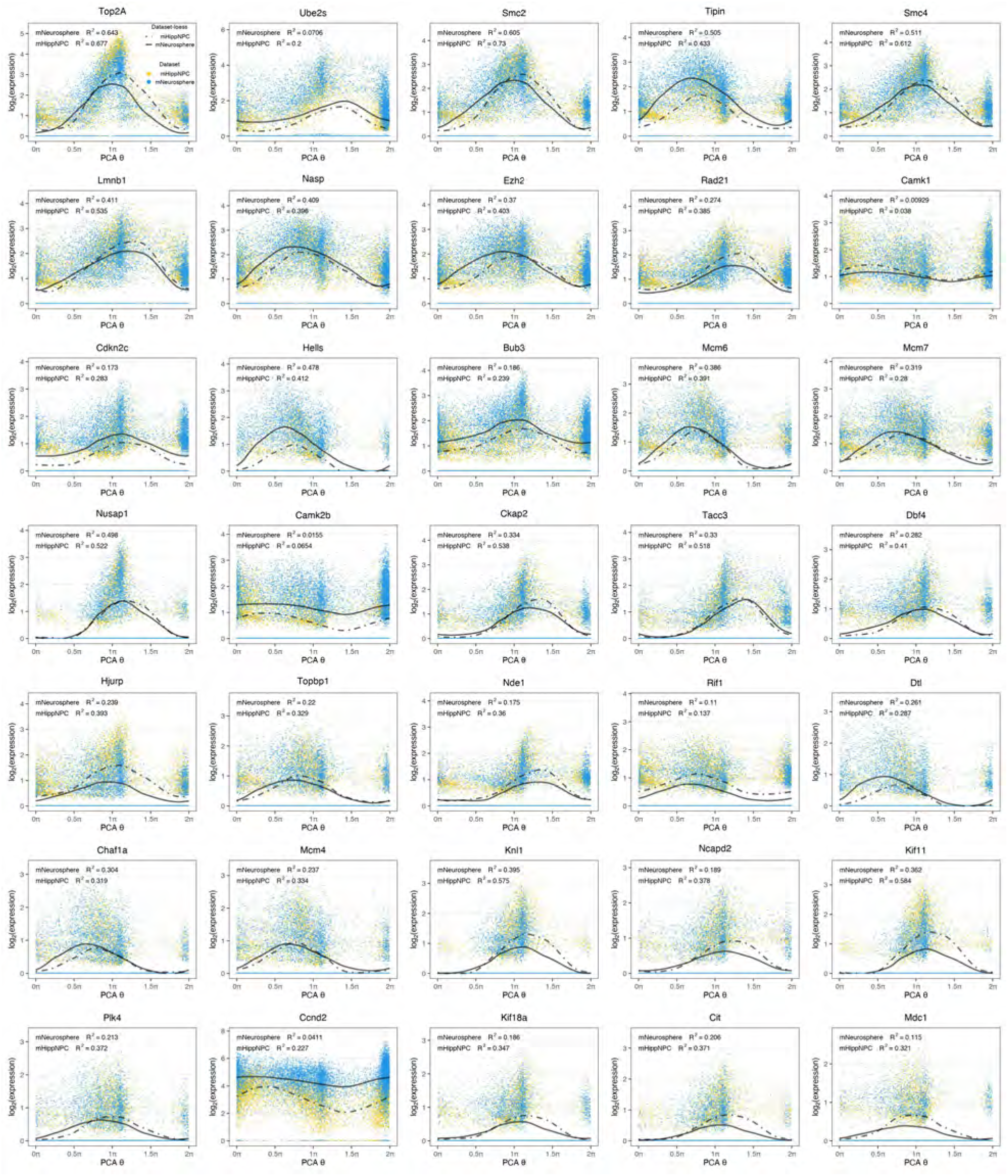
Expression dynamics of top ranked genes. Similar to Figure 2d and e, but now showing all overlapped projection genes with absolute weights greater than 0.1 in either PC1 or PC2 of either dataset. Yellow points are cells of mHippNPC data, while blue points are cells of mNeurosphere data. Two loess lines were fitted for two dataset respectively. There is high agreement of the dynamics between datasets.

**Supplementary Figure S7.**
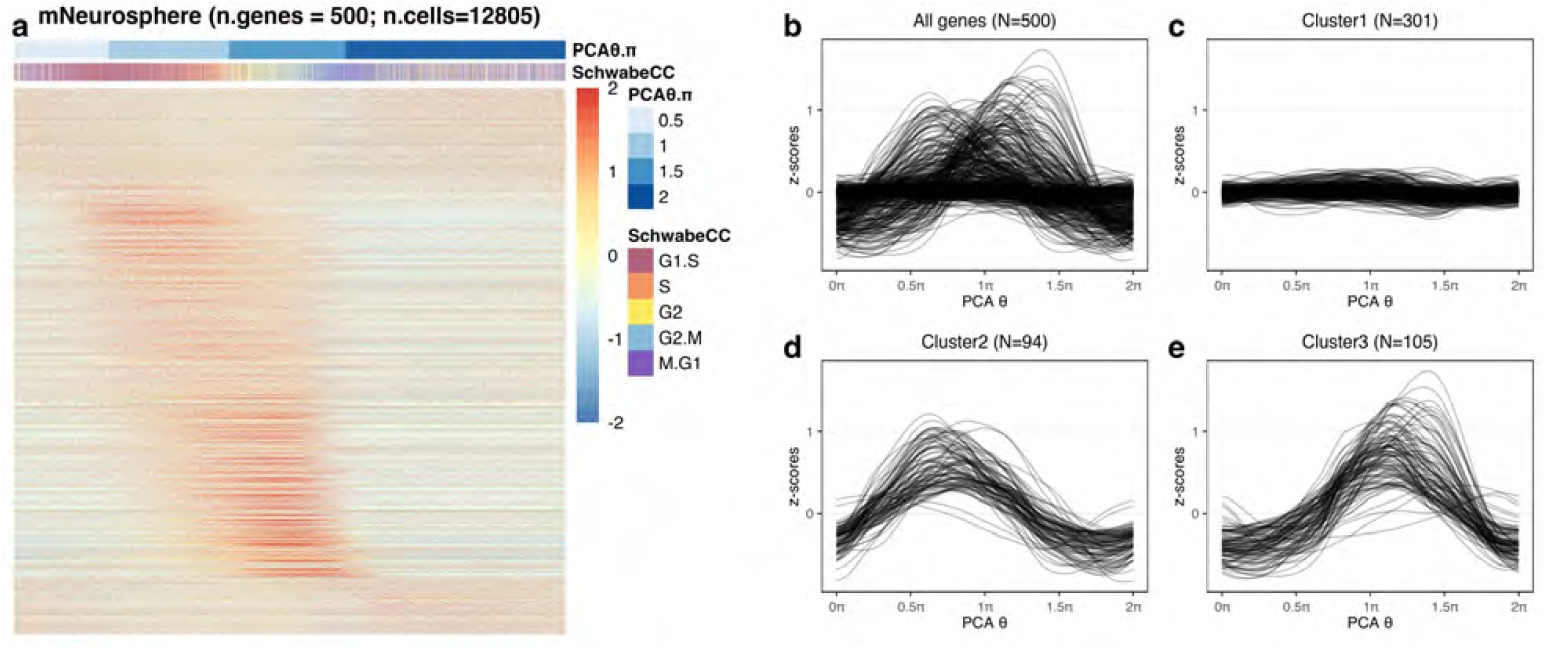
Characteristics of expression patterns of the mNeurosphere reference. **(a)** Heatmap shows the z-scores of 500 projections genes in the mNeurosphere data. Each row represents a gene and each column represents a cell, ordered by the cell cycle position *θ* from PCA. We also annotate the position of half *π* as the cells are not uniformed distributed along 0 to 2*π*. **(b)** The fitted loess line of z-scores over cell cycle position *θ* for all 500 projection genes. **(c-e)** The three different clusters in (b). **(c)** The cluster of genes with highest z-scores less than 0.5. **(d)** The cluster of genes with highest z-scores greater than 0.5 and peak position before *π* This cluster corresponds to high expression genes at G1/S stage. **(e)** The cluster of genes with highest z-scores greater than 0.5 and peak position after *π*. This cluster corresponds to high expression genes at G2/M stage.

**Supplementary Figure S8.**
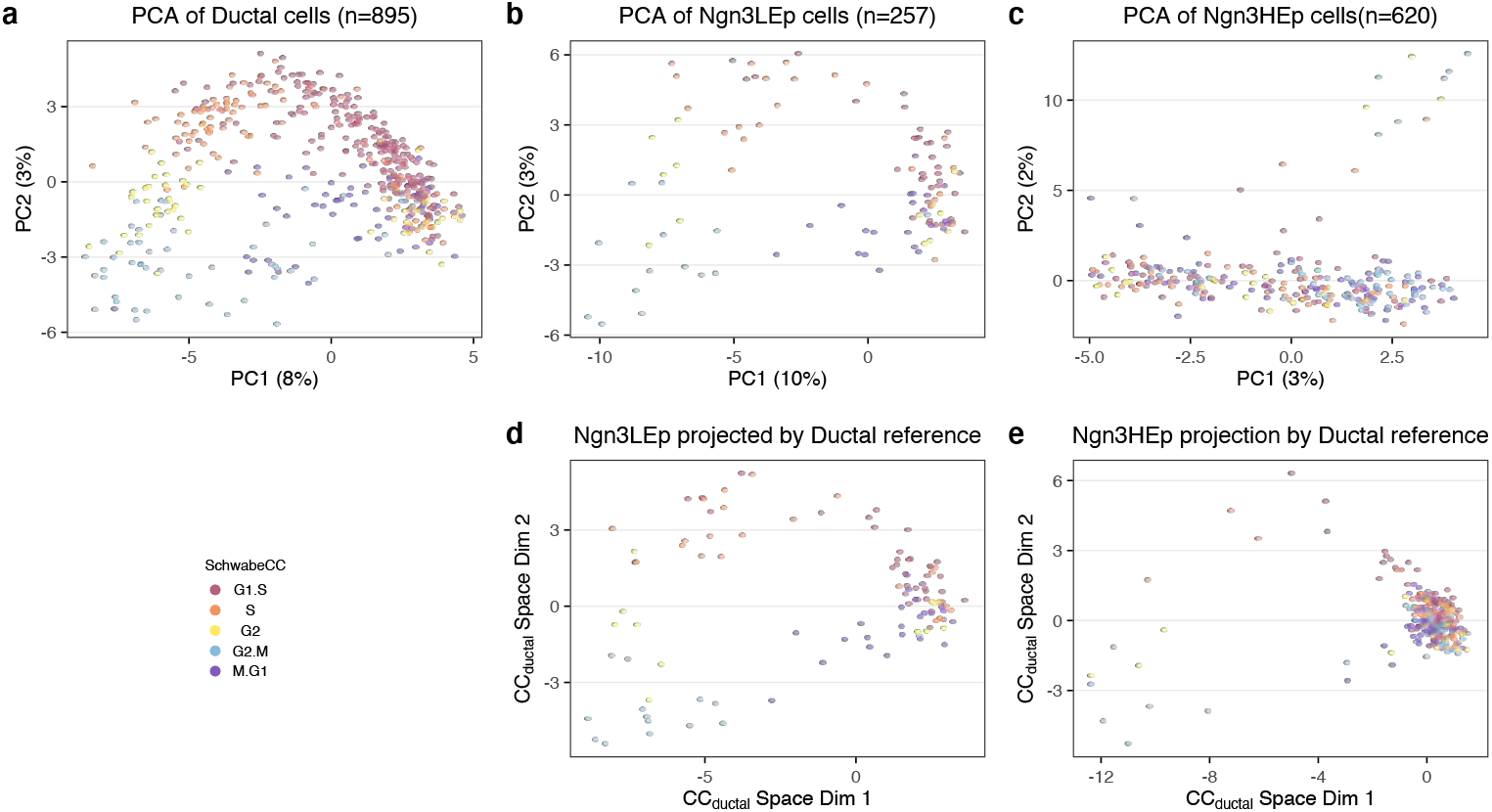
PCA and projections of the mouse developing pancreas data. **(a-c)** The top 2 PCs of GO cell cycle genes of the the three most multipotent cell types in the mouse developing pancreas data. PCA was performed independently for each cell type. Note that panel (a) reproduces Figure 3c. **(d)** Projection of allNgn3LEP cells of mPancreas data using the learned top 2 PCs weights on GO cell cycle genes of Ductal cells. **(e)** Projection of allNgn3HEP cells of mPancreas data using the learned top 2 PCs weights on GO cell cycle genes of Ductal cells.

**Supplementary Figure S9.**
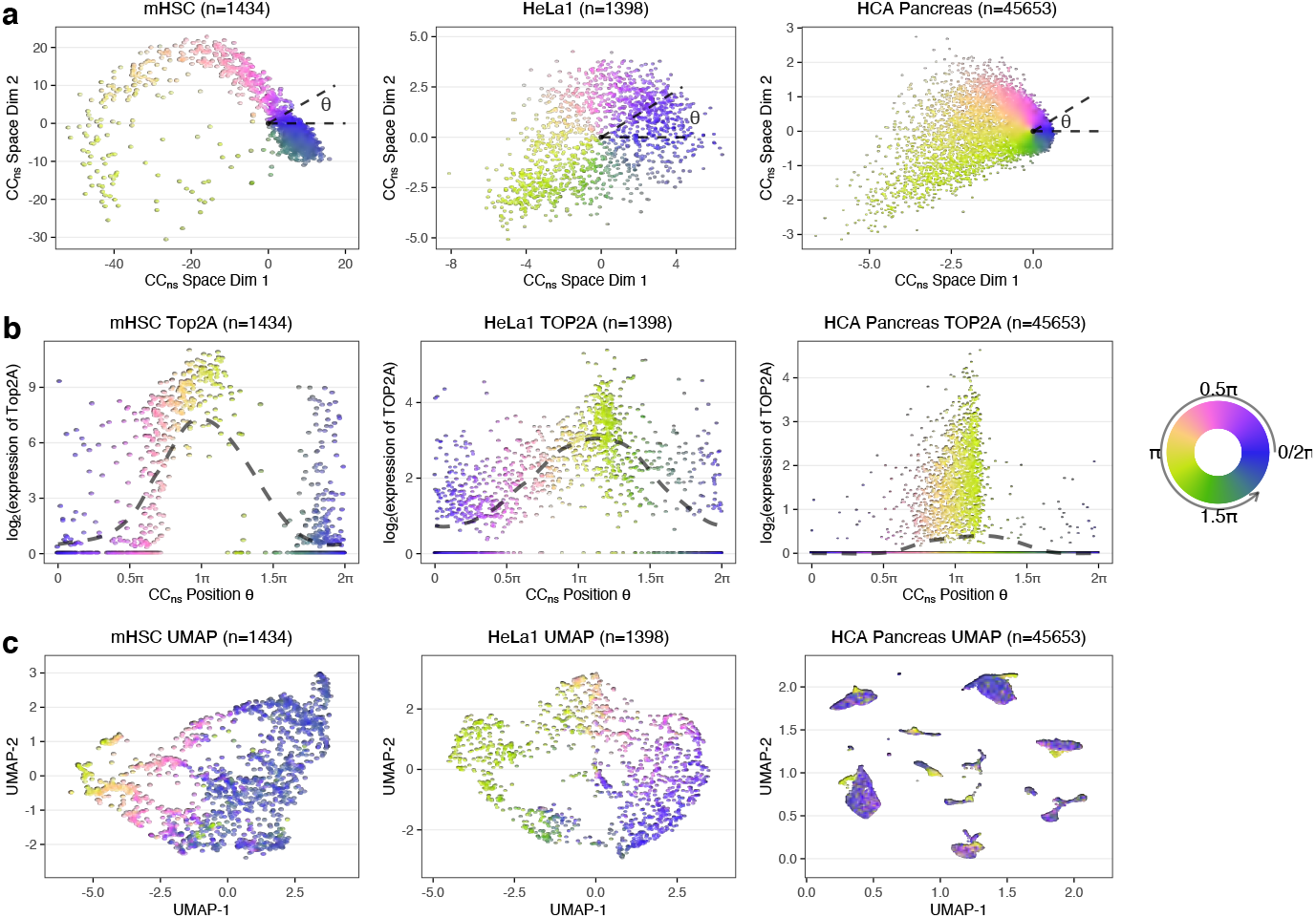
A pre-learned rotation matrix learned from proliferating cortical neurospheres enables cell cycle position estimation in other proliferating datasets. This figure include three other datasets in addition to the four datasets in Figure 4. **(a)** Different datasets (mouse hematopoietic stem cell, Hela set 1 and human fetal pancreas dataset.) projected into the cell cycle embedding defined by the cortical neurosphere dataset. Cell cycle position *θ* is estimated using polar angle. **(b)** Inferred expression dynamics of Top2A(or TOP2A for human), with a periodic loess line (Methods). **(c)** UMAP embeddings of top variable genes. All the cells are colored by cell cycle position using a circular color scale.

**Supplementary Figure S10.**
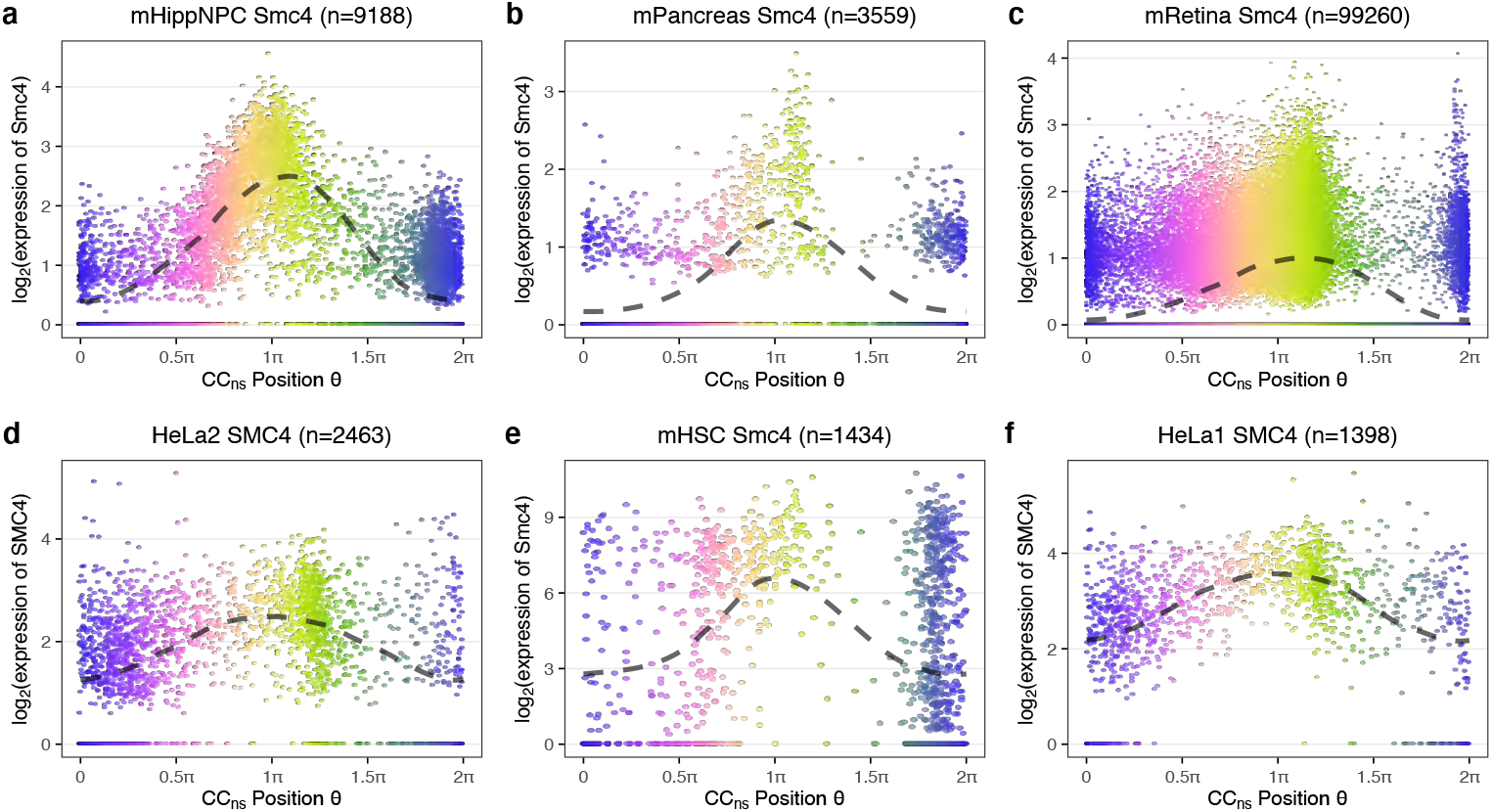
The dynamics of Smc4 expression over cell cycle position *θ*. Inferred expression dynamics of Smc4(or SMC4 for human) over cell cycle position inferred using cortical neurospheres reference, with a periodic loess line (Methods) for **(a)** hippocampal NPCs, **(b)** mouse pancreas, **(c)** mouse retina, **(d)** Hela set 2, **(e)** mouse hematopoietic stem cell, and **(f)** Hela set 1 data. These data are the same data used in Figure 4 and Supplementary Figure S9.

**Supplementary Figure S11.**
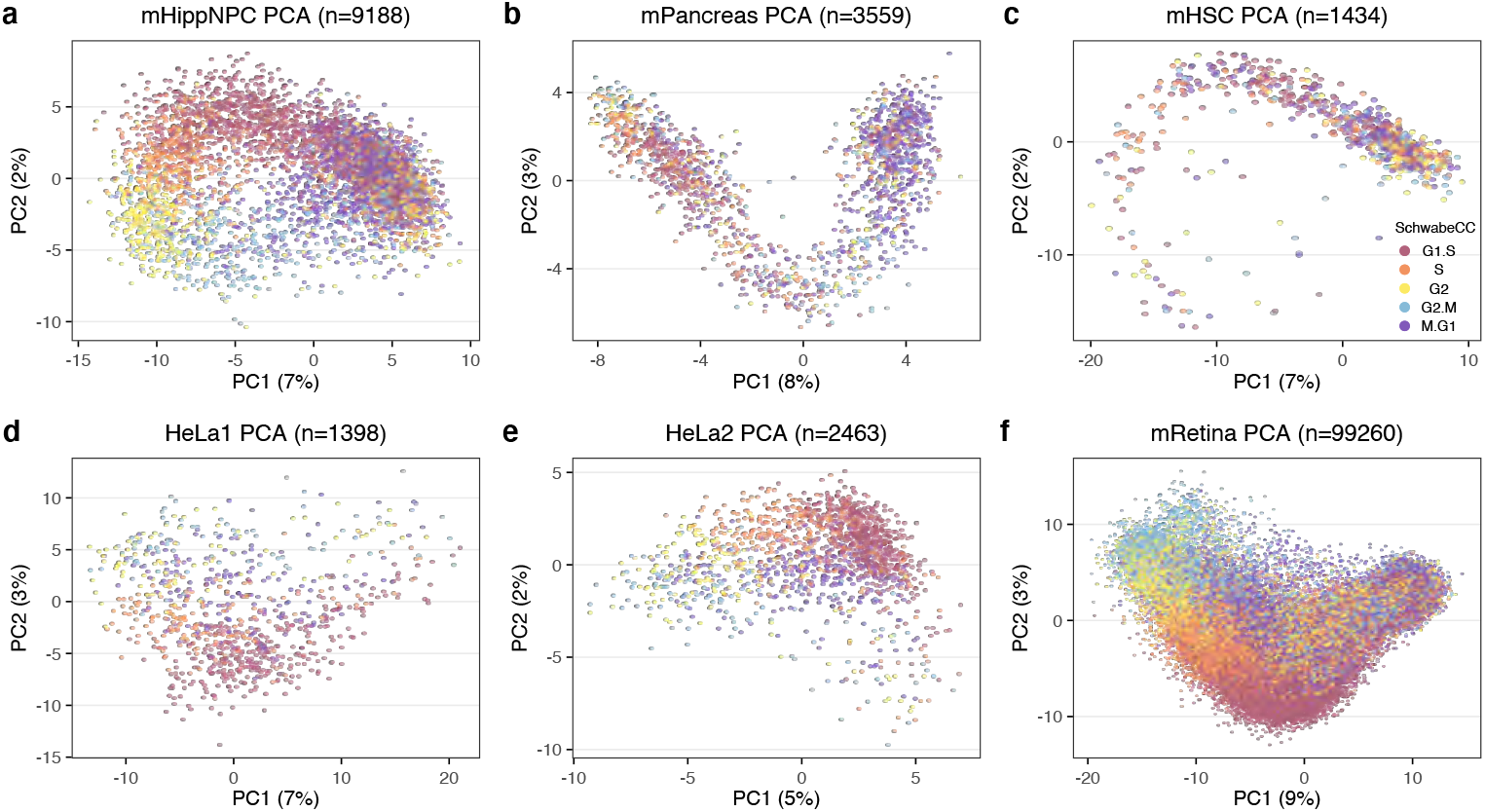
The top 2 PCs of GO cell cycle genes. The figure consists top 2 PCs of PCA performed on GO cell cycle genes of each dataset. They serve as companion figures to Figure 4 and Supplementary Figure S9. Note that the cell cycle progression is hidden by direct PCA on datasets with higher heterogeneity, such as mPancreas and mRetina dataset, while cell cycle progression is visible in other datasets.

**Supplementary Figure S12.**
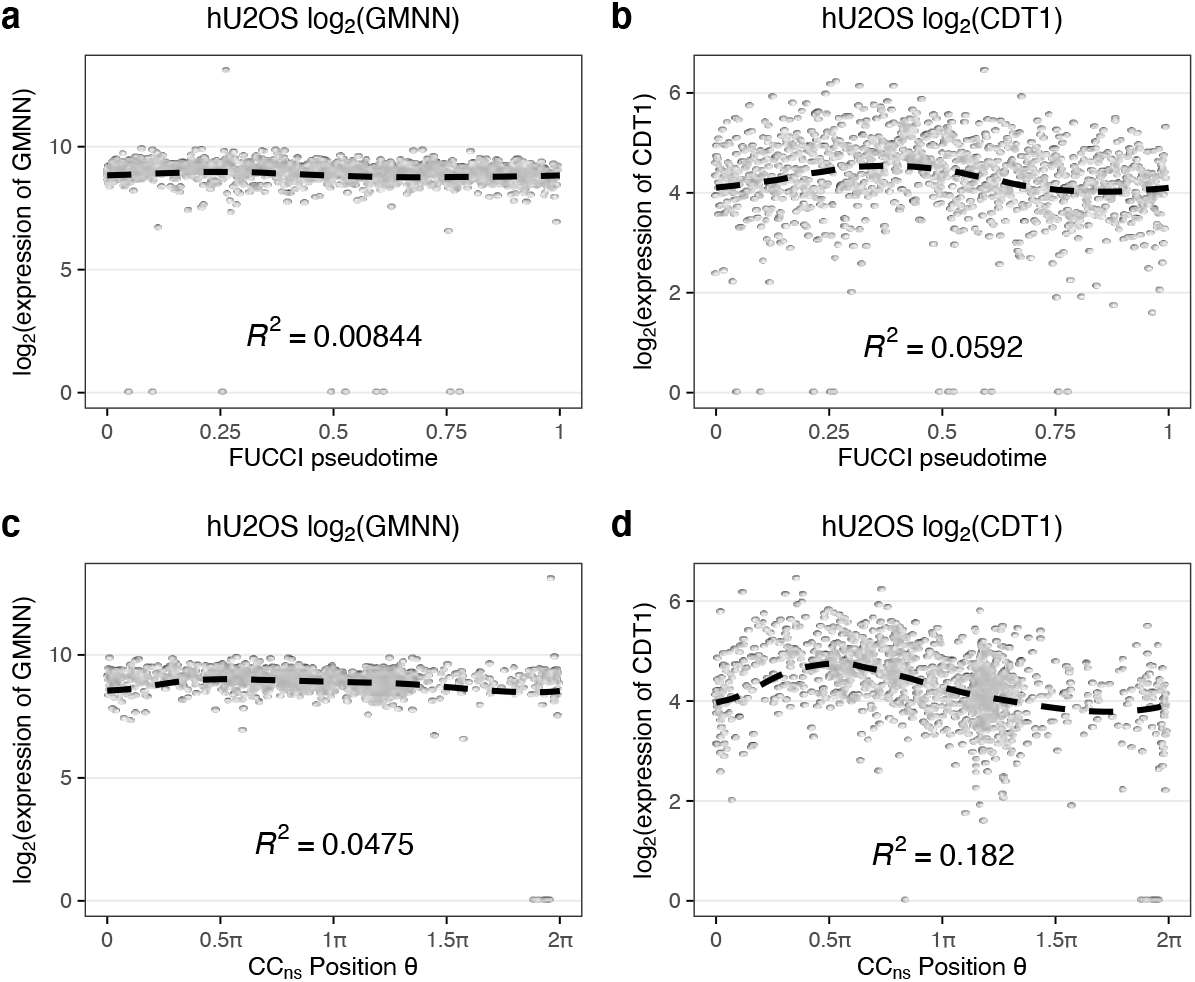
Expression dynamics of GMNN and CDT1 on FUCCI pseudotime and tricycle position of hU2OS data. All data is from the hu2OS dataset. **(a,b)** The gene expression of (a) GMNN and (b) CDT1 as a function of FUCCI pseudotime derived using imaging of the protein levels. **(c,d)** Like (a,b) but as a function of tricycle cell cycle position inferred using scRNA data. Note that GMNN has constant expression over the cell cycle while CDT1 oscillates. This strongly suggests that the protein level of GMNN is regulated post transcriptionally.

**Supplementary Figure S13.**
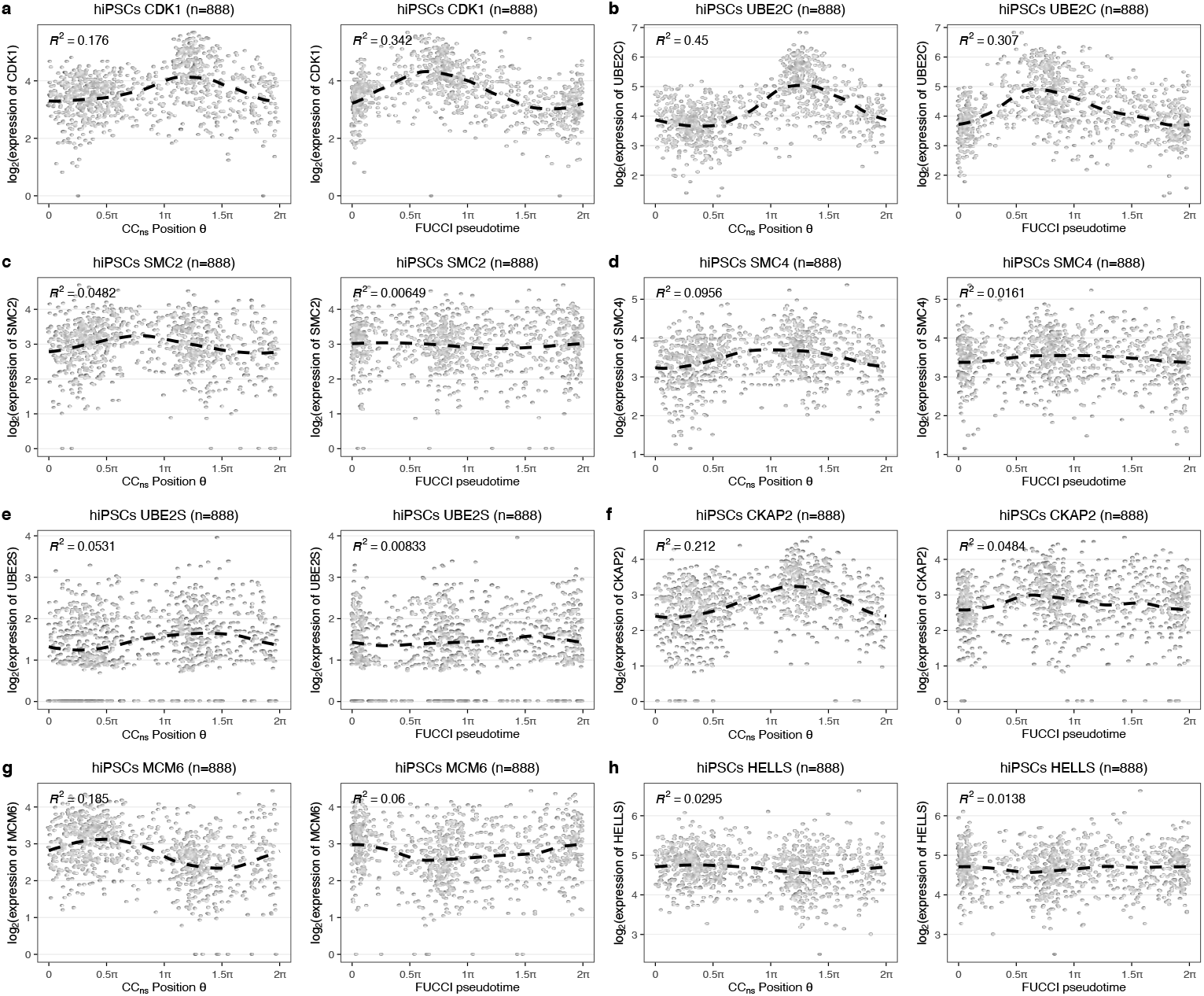
Expression dynamics of selected cell cycle genes of hiPSCs dataset. Similar to Figure 5e,f, but now we show more cell cycle related genes. In each panel, the left sub-panel shows the expression of the gene over tricycle cell cycle position *θ* using mNeurosphere reference, and the right sub-panel over the FUCCI pseudotime inferred by Hsiao et al. (2020). Periodic loess lines and *R^2^* are added for each sub-panel (Methods).

**Supplementary Figure S14.**
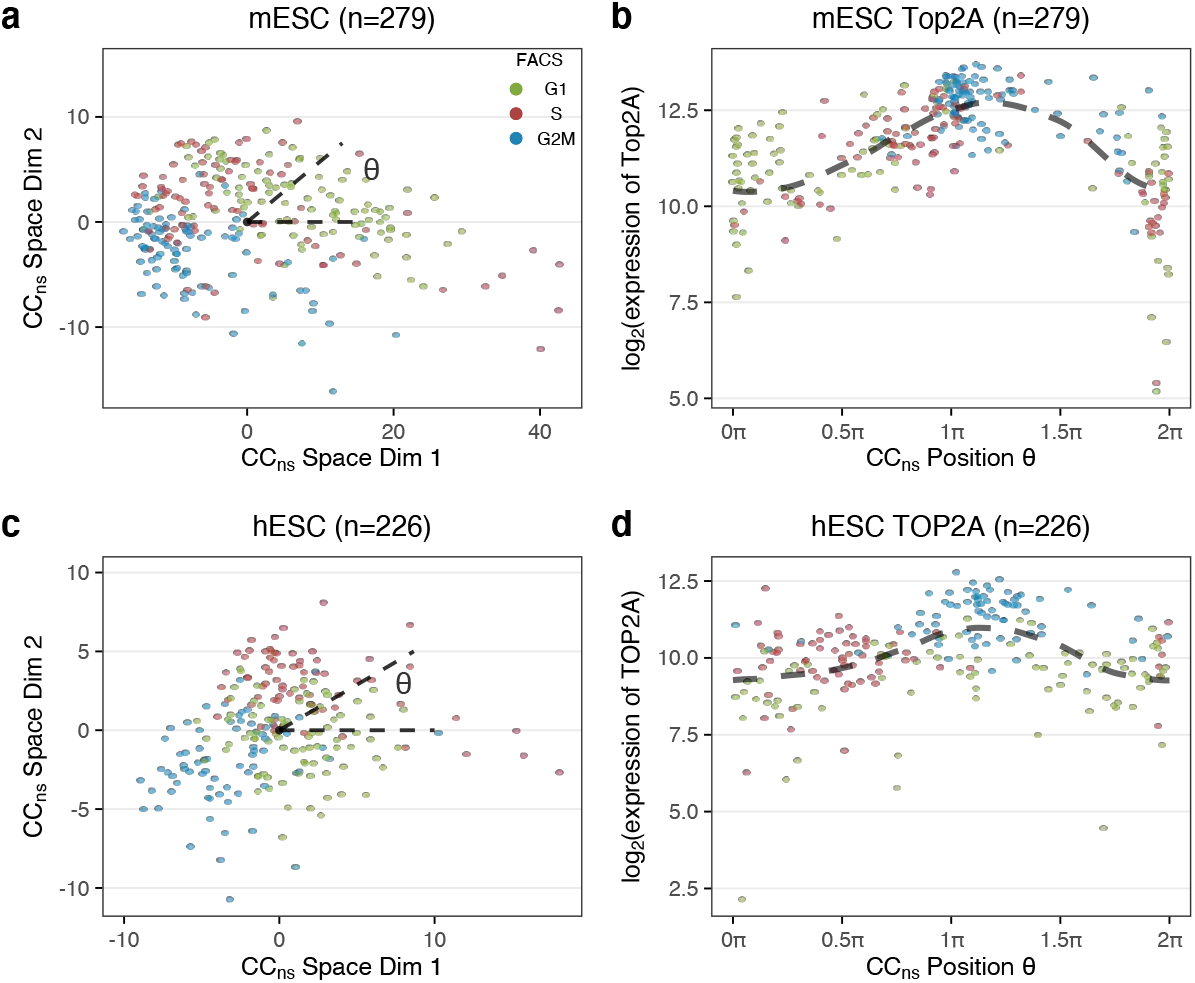
Evaluation of tricycle on FACS datasets. **(a-b)** Data from Buettner et al. (2015). **(a)** The data is projected to the cell cycle embedding defined by the cortical neurosphere dataset. Cells are colored by FACS labels. **(b)** Expression dynamics of Top2A with a periodic loess line using tricycle cell cycle position estimated by projection in (a). **(c-d)** Similar to (a,b), but for data from Leng et al. (2015).

**Supplementary Figure S15.**
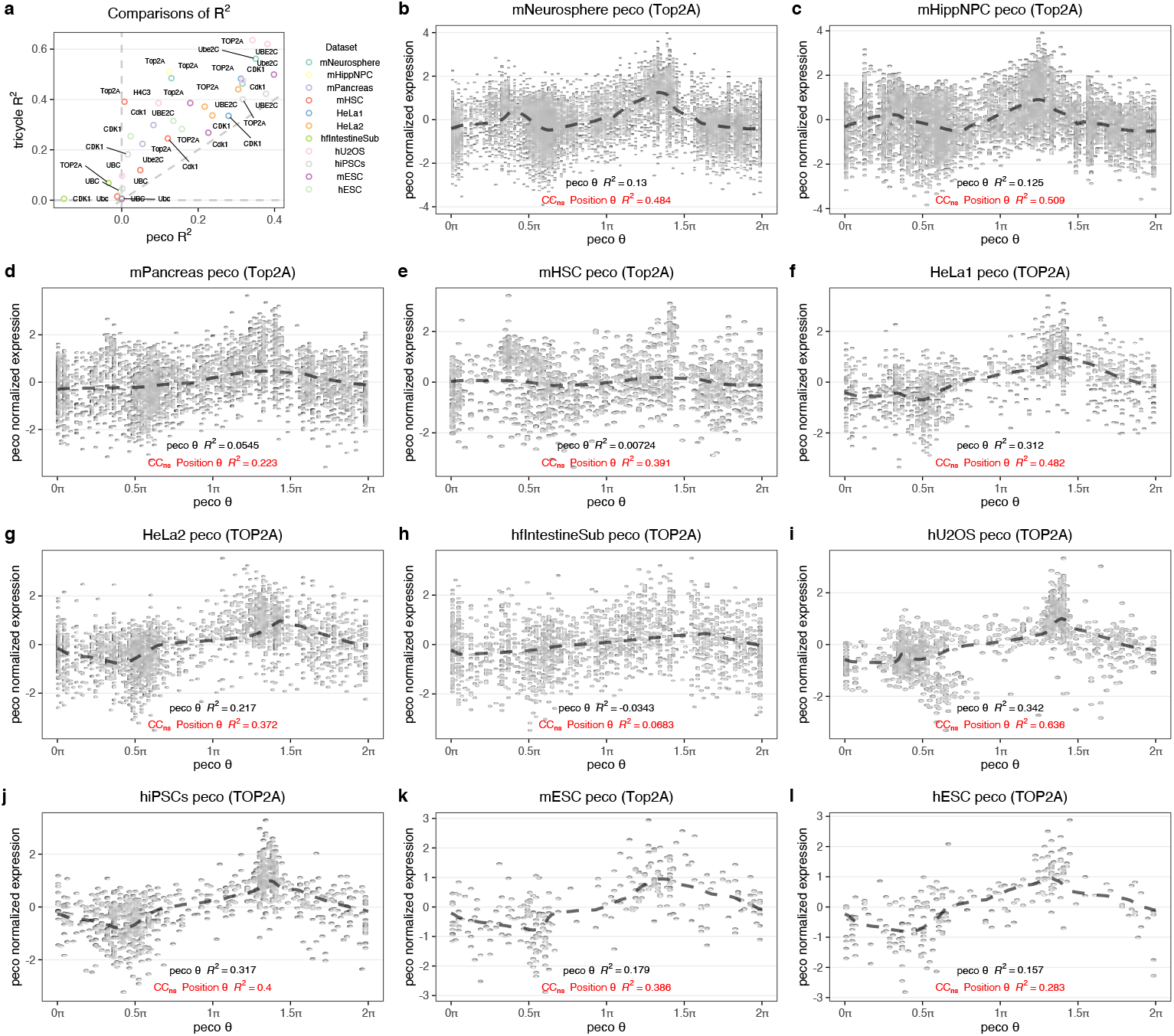
Expression dynamics of cell cycle genes on peco cell cycle position. We run peco on all dataset described in Table 1, except mRetina and human fetal tissues. mRetina data has too many cells, and for human fetal tissues, we only use a subset of random 2000 cells from intestine data (hfIntestineSub). For each data, the *R*^2^ of expression dynamics of Cdk1, Top2A, Ube2C and H4c3, as long as they exist in the target dataset, over peco inferred *θ* are computed. **(a)** Comparison of *R*^2^ of peco inferred *θ* and *R*^2^ of loess line on tricycle inferred *θ* using mNeurosphere reference for all datasets and genes mentioned above. **(b-l)** Expression dynamics of Top2A (TOP2A for human) over peco inferred *θ* for each dataset. Note that in these panels, the y-axis represents the peco normalized expression values, as peco has its own normalization requirement. We annotate the each panel with *R*^2^ of loess line calculated on peco inferred *θ* and *R*^2^ of loess line on tricycle inferred *θ* using mNeurosphere reference (although we have not plotted out the expression dynamics over tricycle inferred *θ*). Across all datasets and genes, the tricycle inferred *θ*s have greater *R*^2^ to peco *θ*, and are highlighted as red.

**Supplementary Figure S16.**
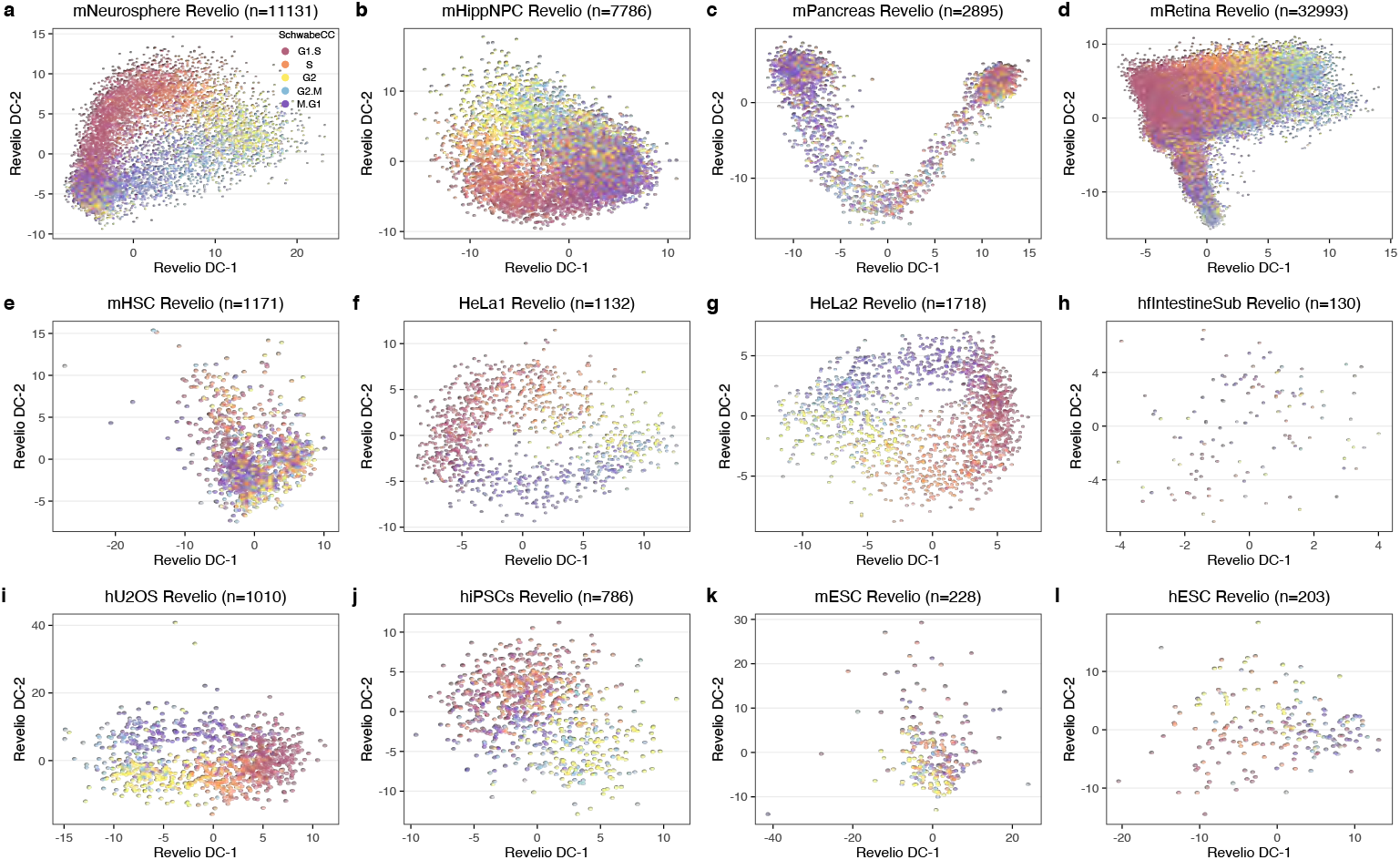
Cell cycle embeddings by Revelio. The cell cycle embedding produced by Revelio for each data. Cells are colored by 5 stage cell cycle representation, inferred using the original Schwabe method (Schwabe et al., 2020) as implemented in the Revelio package. Note that all cells without a valid stage assignment (assigned to “NA”) are removed by the functions in Revelio package.

**Supplementary Figure S17.**
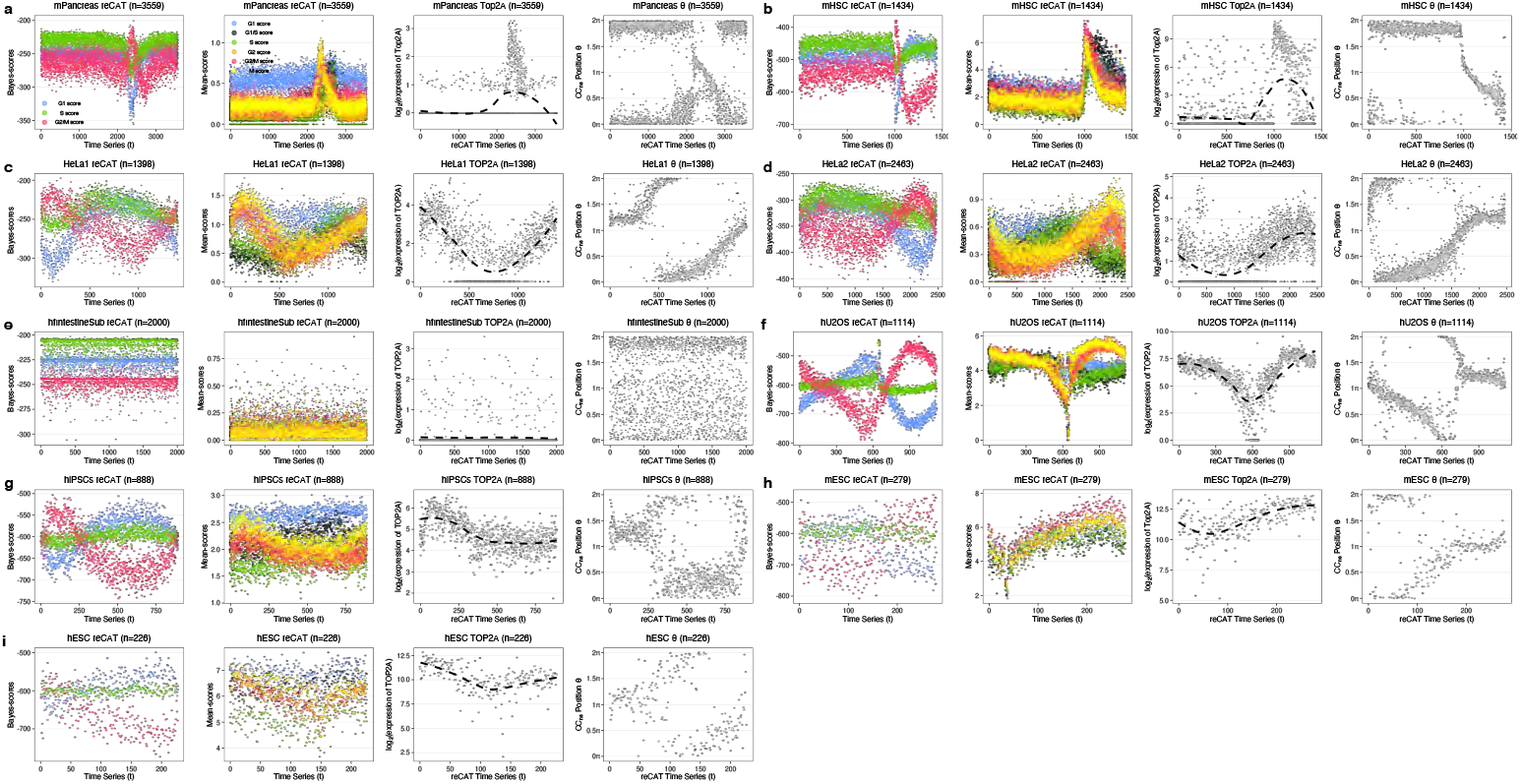
Cell cycle stage and order estimations by reCAT. Panels show the cell cycle stage scores and cell orders estimated by reCAT for **(a)** mPancreas, **(b)** mHSC, **(c)** HeLa set 1, **(d)** HeLa set 2, **(e)** hfIntestineSub, **(f)** hU2OS, **(g)** hiPSCs, **(h)** mESC, and **(i)** hESC data. Each data point is a cell. For each data, the first sub-panel shows the Bayes scores for G1, S, and G2/M stage over the estimated cell orders(time series t). For each cell, there will be three scores (data points) colored by stage. The second sub-panel shows the mean scores for G1, G1/S, S, G2, G2/M, and M stage over the estimated cell orders. For each cell, there will be six scores (data points) colored by stage. The third sub-panel is the expression dynamic of Top2A(or TOP2A for human) over reCAT estimated cell orders. Note that although reCAT package provide function to assign cell cycle stage, it requires manual input cutoff for Bayes scores. It is unrealistic for us to pick some appropriate cutoffs for most of the datasets presented here. For example, for mPancreas data in (a), we cannot decide which region has the consistent G1 scores. The last sub-panels compares tricycle cell cycle position using mNeurosphere reference and reCAT cell orders.

**Supplementary Figure S18.**
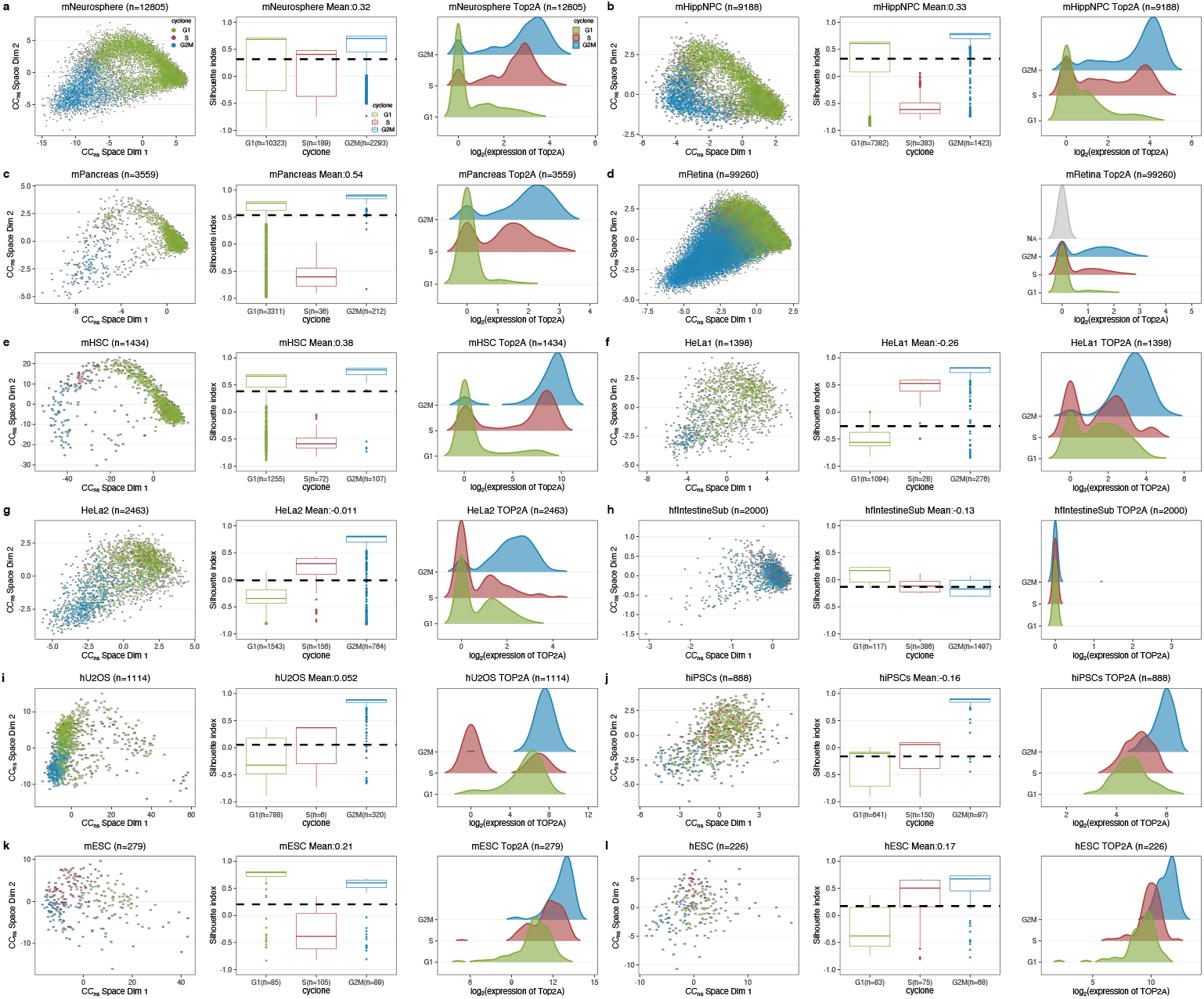
Comparison between cyclone assigned stages and tricycle cell cycle position using mNeurosphere reference. Each panel describe one data, specifically for **(a)** mNeurosphere, **(c)** mHippNPC, **(c)** mPancreas, **(d)** mRetina, **(e)** mHSC, **(f)** HeLa set 1, **(g)** HeLa set 2, **(h)** hfIntestineSub, **(i)** hU2OS, **(j)** hiPSCs, **(k)** mESC, **(l)** hESC data. For each data, the first sub-panel shows the cell cycle embedding projection by mNeurosphere reference, and each point is a cell, colored by cyclone inferred cell cycle stage. The second sub-panel shows silhouette index computed using angular separation distance of tricycle cell cycle position *θ* estimated using mNeurosphere reference (Methods), stratified by cyclone inferred cell cycle stage. The mean silhouette index across all cells is given in the title. Boxes indicate 25th and 75th percentiles. Whiskers extend to the largest values no further than 1.5 × interquartile range (IQR) from these percentiles. For mRetina data, the pairwise distance matrix is too big to substantiate, so we could not compute silhouette index. The third sub-panel shows the marginal density of Top2A(or TOP2A for human) expression conditioned on cyclone cell cycle stage.

**Supplementary Figure S19.**
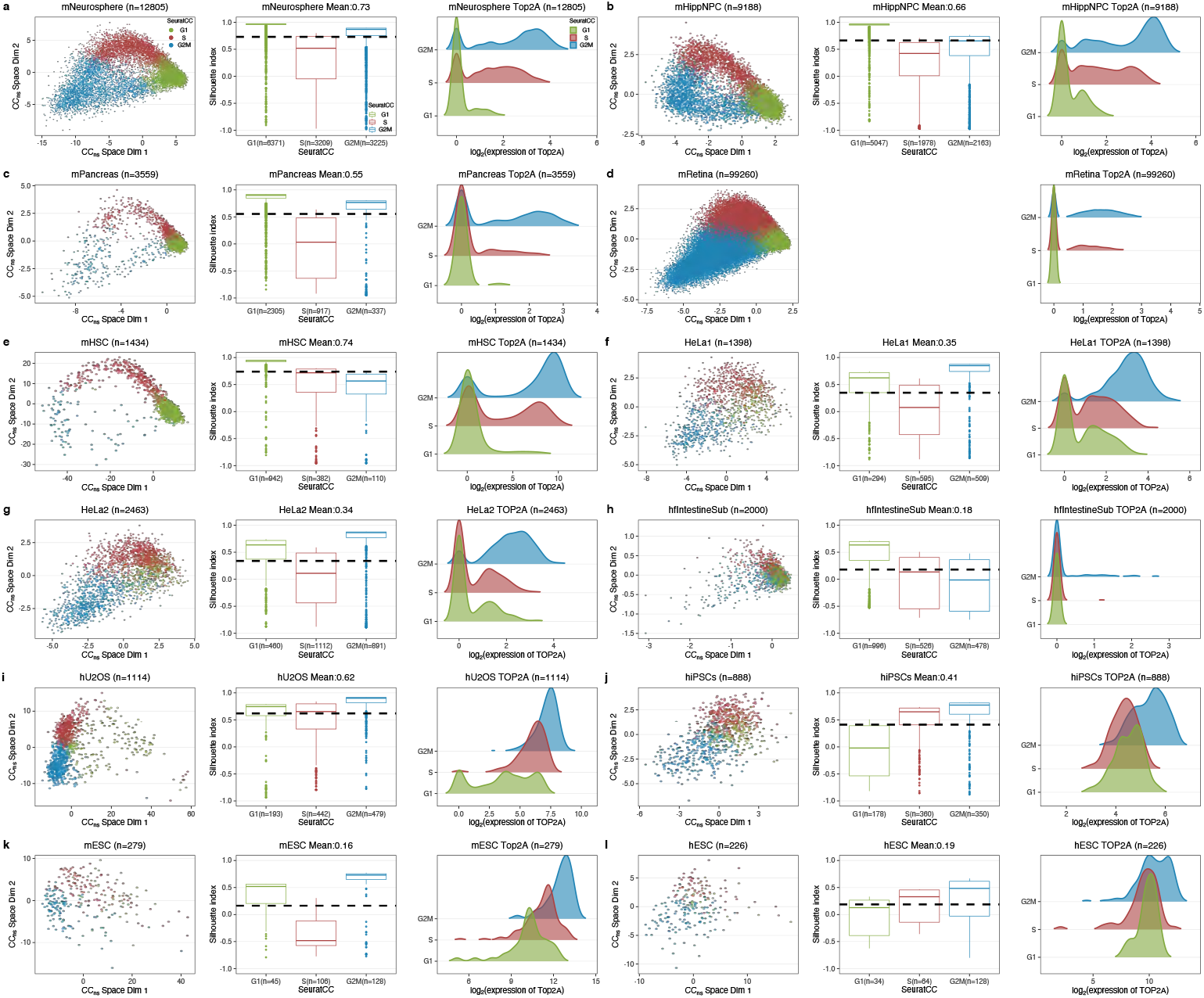
Comparison between Seurat assigned stages and tricycle cell cycle position using mNeurosphere reference. Each panel describe one data, specifically for **(a)** mNeurosphere, **(c)** mHippNPC, **(c)** mPancreas, **(d)** mRetina, **(e)** mHSC, **(f)** HeLa set 1, **(g)** HeLa set 2, **(h)** hfIntestineSub, **(i)** hU2OS, **(j)** hiPSCs, **(k)** mESC, **(l)** hESC data. For each data, the first sub-panel shows the cell cycle embedding projection by mNeurosphere reference, and each point is a cell, colored by Seurat inferred cell cycle stage. The second sub-panel shows silhouette index computed using angular separation distance of tricycle cell cycle position *θ* estimated using mNeurosphere reference (Methods), stratified by Seurat inferred cell cycle stage. The mean silhouette index across all cells is given in the title. Boxes indicate 25th and 75th percentiles. Whiskers extend to the largest values no further than 1.5·IQR from these percentiles. For mRetina data, the pairwise distance matrix is too big to substantiate, so we could not compute silhouette index. The third sub-panel shows the marginal density of Top2A(or TOP2A for human) expression conditioned on Seurat cell cycle stage.

**Supplementary Figure S20.**
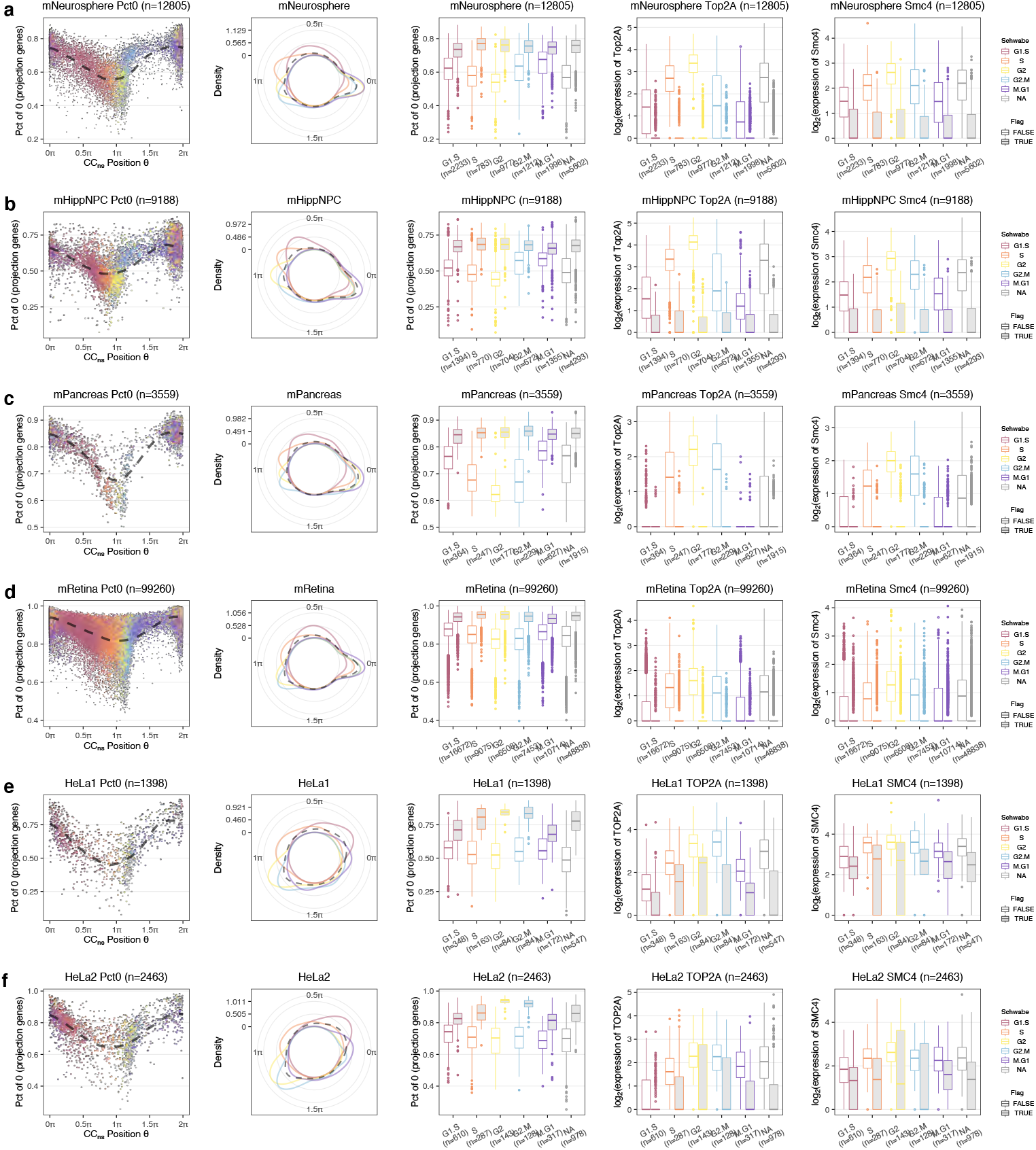
Comparison between SchwabeCC 5 stage assignments and tricycle cell cycle position using mNeurosphere reference. Each row or panel contains analysis for a dataset, specifically **(a)** for mNeurosphere, **(b)** for mHippNPC, **(c)** for mPancreas, **(d)** for mRetina, **(e)** for HeLa set 1, **(f)** for HeLa set 2 data. For each data, the first sub-panel shows the dynamics of percentage of non-expressed genes over all overlapped genes with mNeurosphere projection matrix (number of genes with 0 expression divided by the number overlapped genes with mNeurosphere projection matrix) w.r.t. tricycle cell cycle position *θ* using mNeurosphere reference. Cells are colored by 5 stage assignment. The second panel shows the marginal density of tricycle cell cycle position *θ* conditioned on 5 stage assignments using von Mises kernel on polar coordinate system. The third sub-panel shows the percentage of non-expressed genes over all overlapped genes with mNeurosphere projection matrix conditioned on 5 stages assignment and whether cells appear in the G1/G0 cluster - *θ* < 0.25 *π* or *θ* > 1.5*π* as boxplots. The forth and the last sub-panel show the expression of Top2A and Smc4 conditioned on 5 stages assignment and whether cells appear in the G1/G0 cluster. Boxes indicate 25th and 75th percentiles. Whiskers extend to the largest values no further than 1.5·IQR from these percentiles.

**Supplementary Figure S21.**
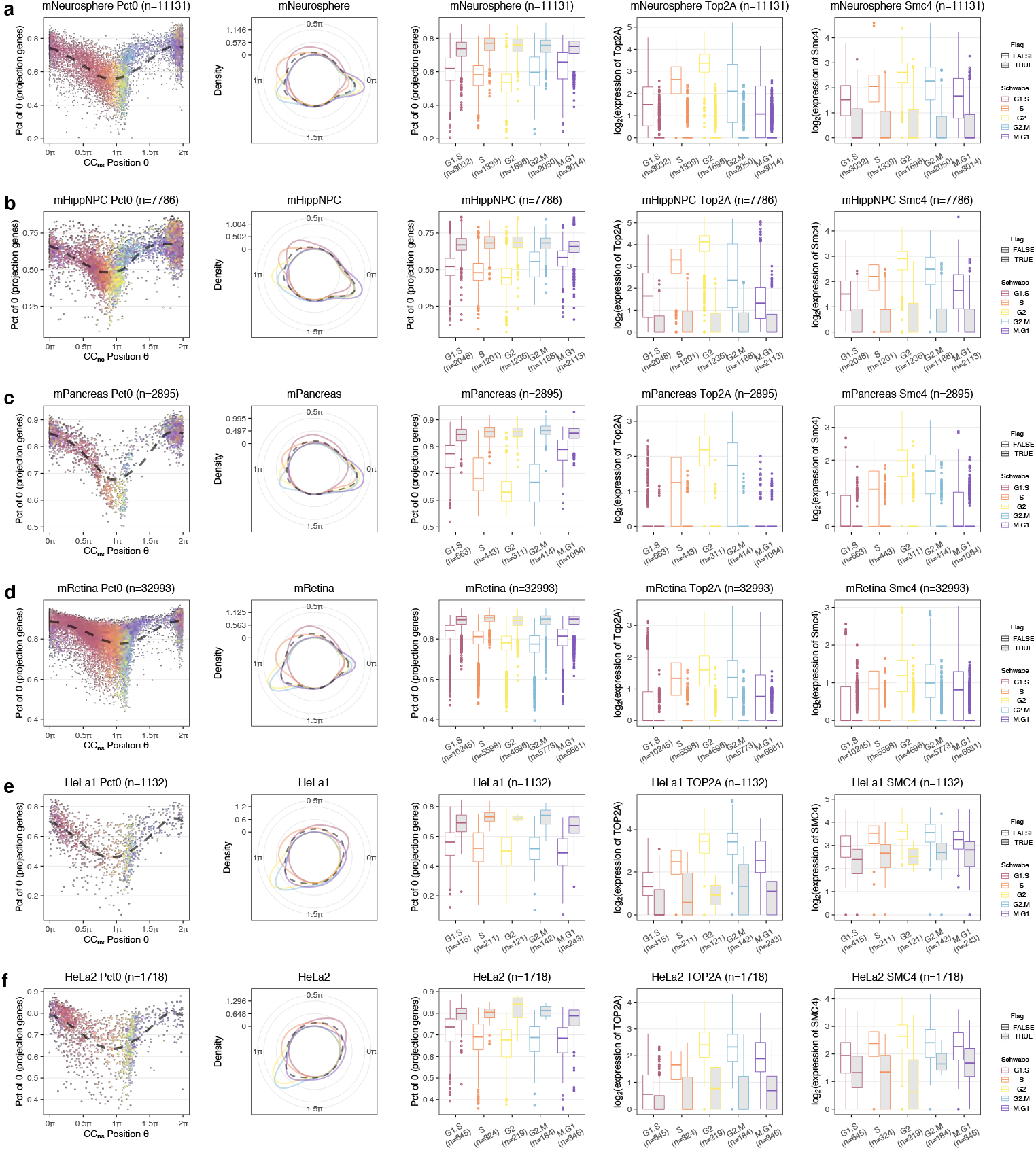
Comparison between original SchwabeCC 5 stage assignments and tricycle cell cycle position using mNeurosphere reference. This figure shows the exact same data and comparisons as in Supplementary Figure S20, but now we use the original SchwabeCC method as implemented in the Revelio package (Schwabe et al., 2020). Note that the number of cells in each dataset is decreased as any cell without a valid stage assignment (assigned to “NA”) is removed by the functions in Revelio package.

**Supplementary Figure S22.**
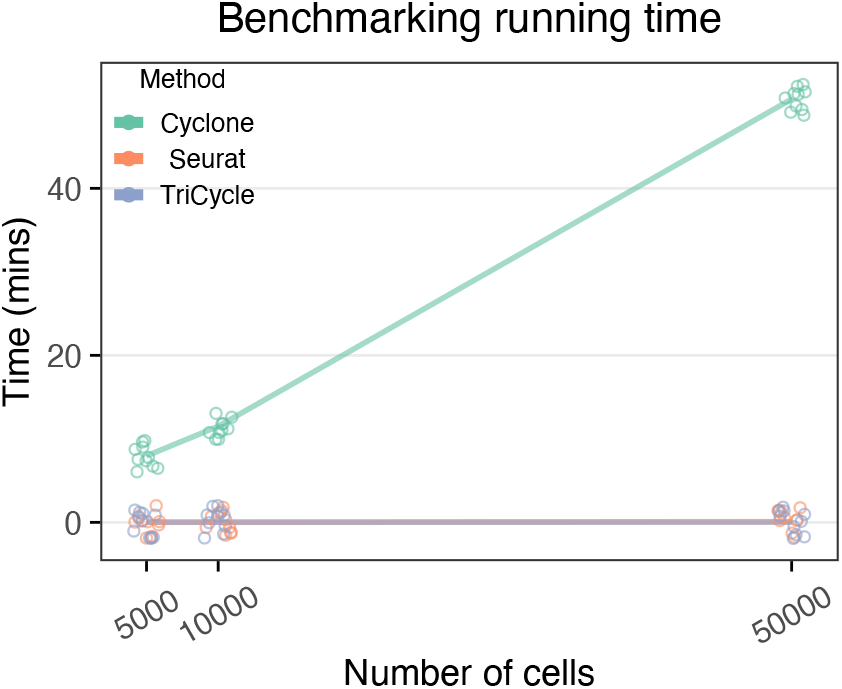
Running time comparisons between cyclone, Seurat, and tricycle cell cycle inference. We record the elapsing time for each method when running them on 10 random subsets of mRetina data with 5000, 10000, and 50000 cells. For cyclone and Seurat, the time is recorded for the cell cycle stage assignment function. For tricycle, the time is recorded for cell cycle position estimation using mNeurosphere reference. Note that we add jitters to the data points to avoid excessive overlaps.

**Supplementary Figure S23.**
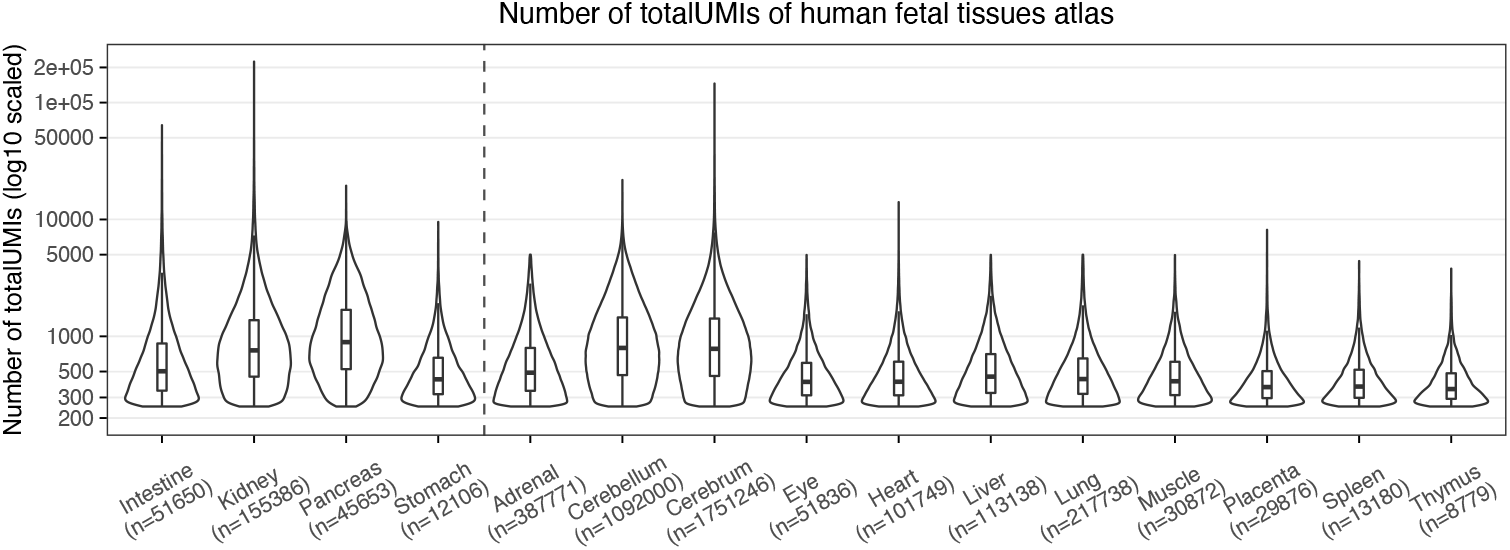
TotalUMIs of human fetal atlas. For each tissue type of the human fetal atlas data (Cao et al., 2020), we show the total UMIs of a cell. The dashed line separates 4 single-cell profiled tissues with 11 single-nuclei profiled tissues. Boxes indicate 25th and 75th percentiles. Whiskers extend to the largest values no further than 1.5 × interquartile range (IQR) from these percentiles.

**Supplementary Figure S24.**
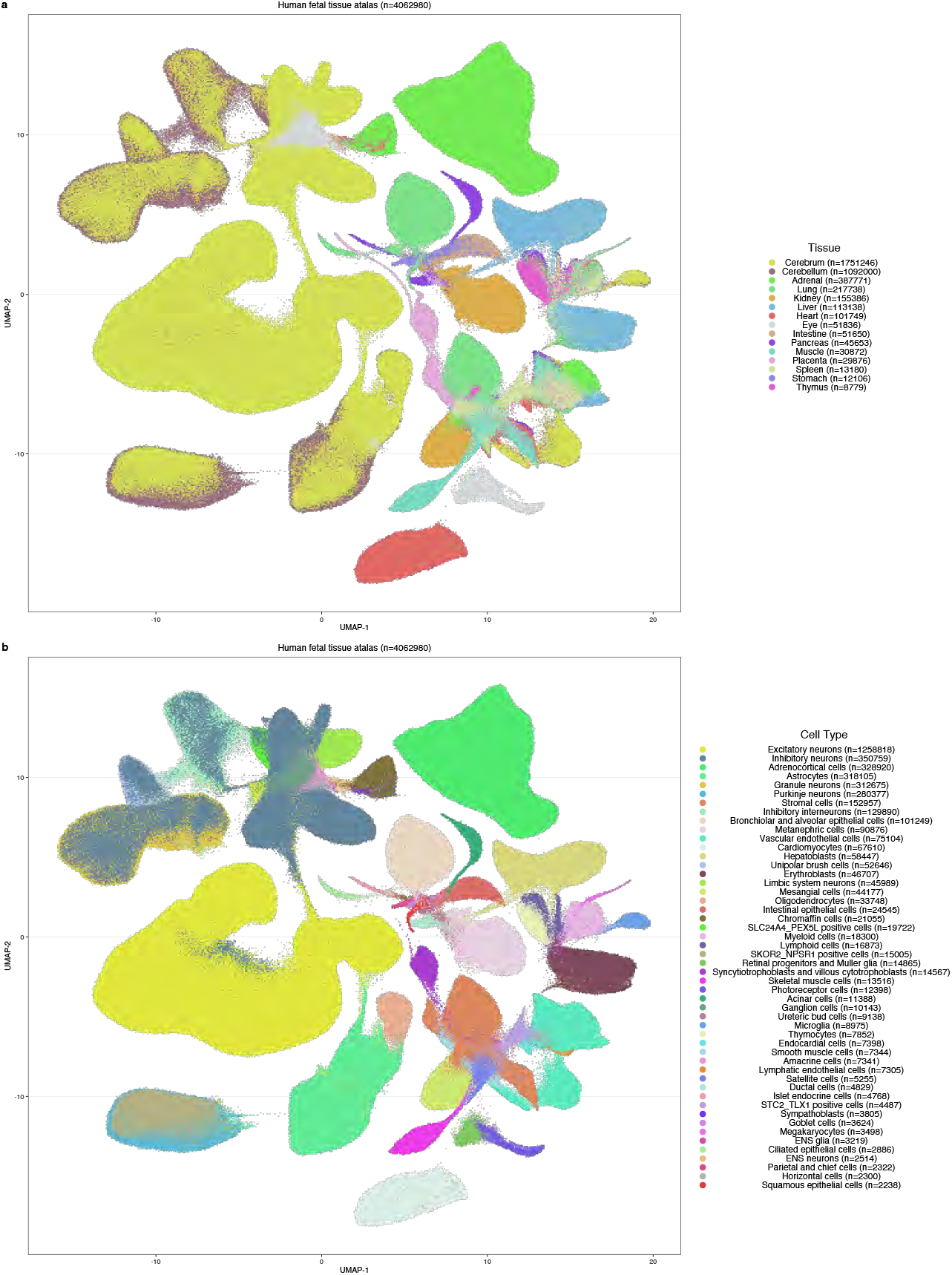
Human fetal tissue atlas UMAP embeddings with all tissues. Human fetal tissue atlas UMAP embeddings with all tissues, colored by **(a)** tissue and **(b)** cell type.

**Supplementary Figure S25.**
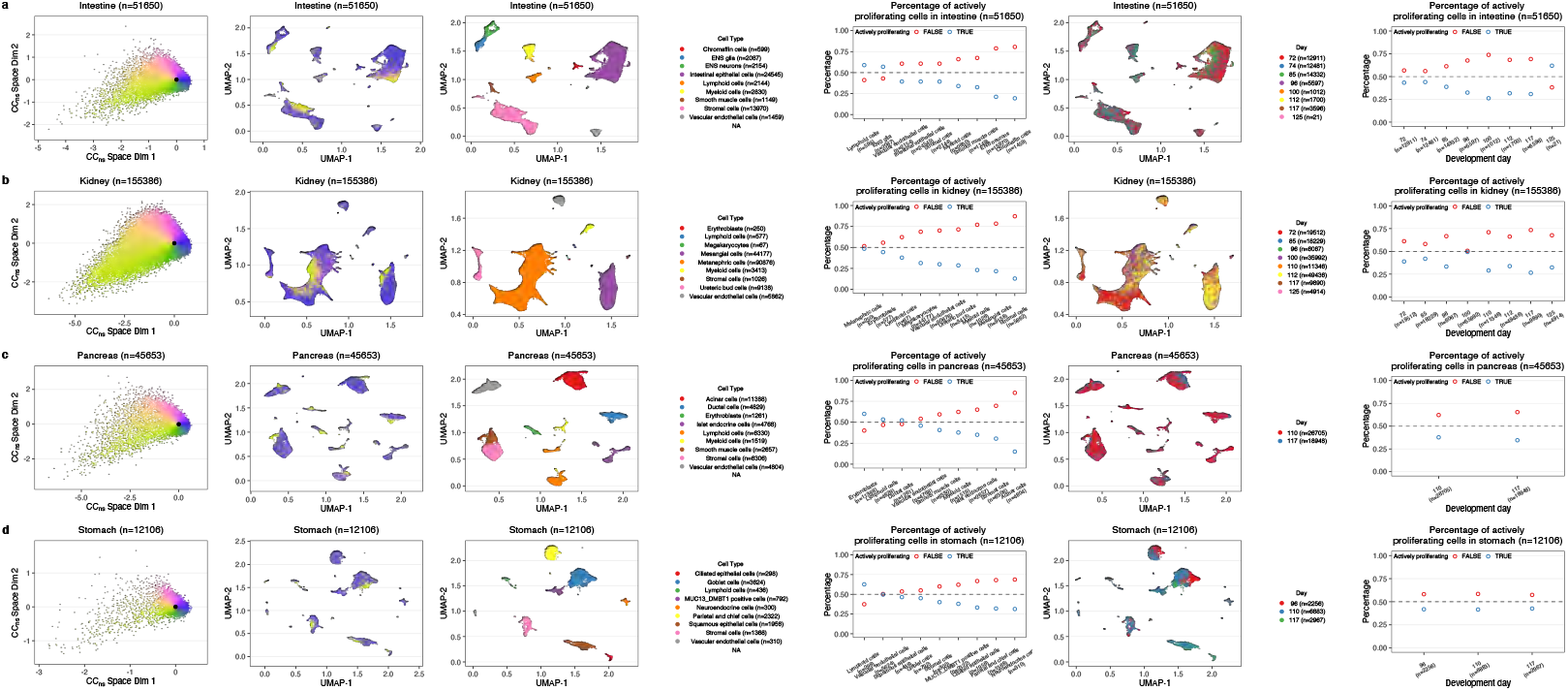
Application of tricycle on 4 single-cell profiled human tissues. We show one tissue type in each row/panel **(a)** intestine, **(b)** kidney, **(c)** pancreas, and **(d)** stomach. For each tissue, the cell cycle embedding using mNeurosphere reference is given in the first sub-panel, tissue-level UMAPs from Cao et al. (2020) colored by cell cycle position *θ* in the second sub-panel, tissue-level UMAPs from Cao et al. (2020) colored by cell type in the third sub-panel, percentage of actively proliferating cells for each cell type in decreasing order in the forth sub-panel, tissue-level UMAPs from Cao et al. (2020) colored by development days in the fifth sub-panel, and percentage of actively proliferating cells for each development day in the last sub-panel.

**Supplementary Figure S26.**
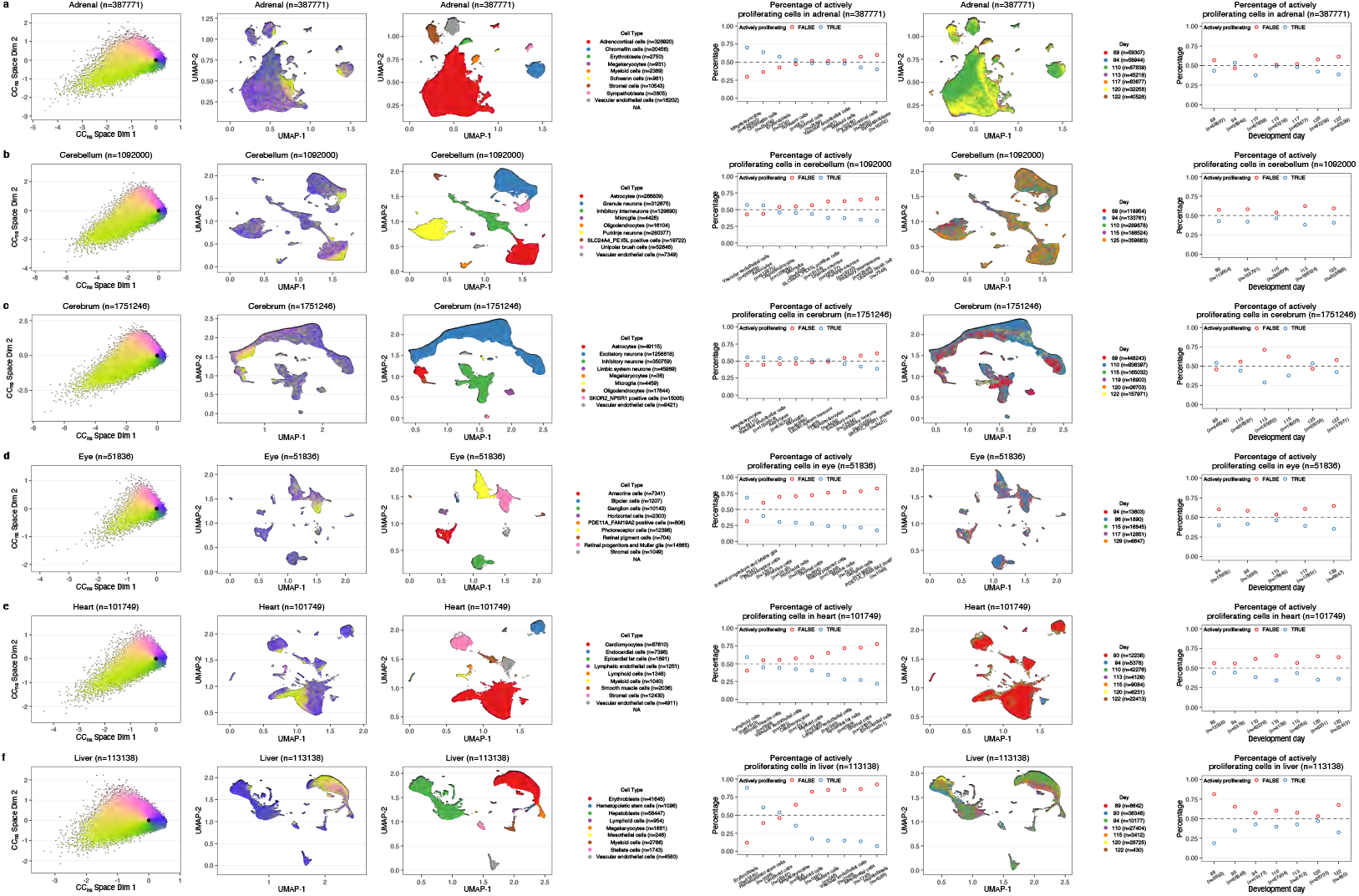
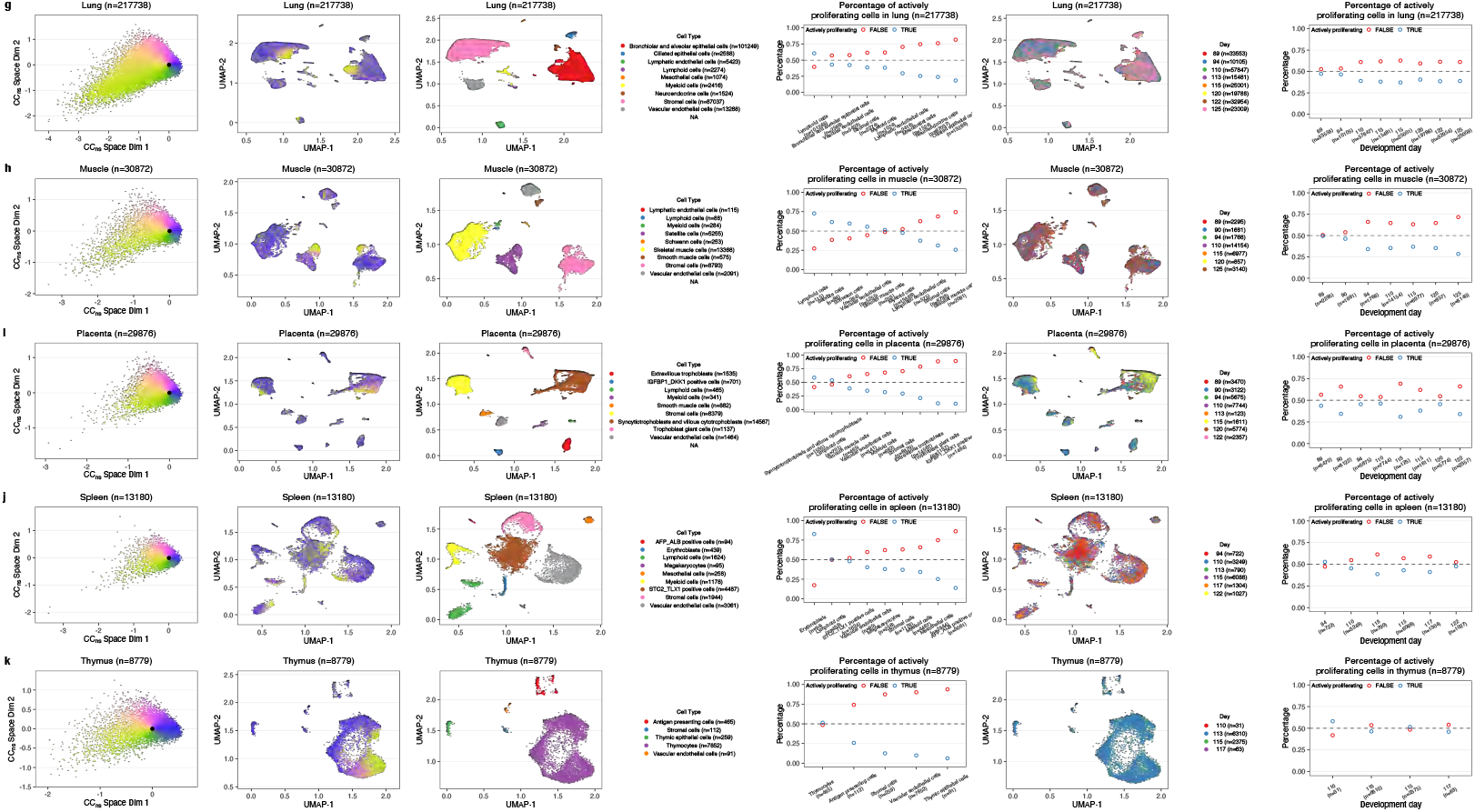
Application of tricycle on 11 single-nuclei profiled human tissues. Similar to Figure S25, but for 11 tissues with single-nuclei RNA profiled. We show one tissue type in each panel **(a)** adrenal, **(b)** cerebellum, **(c)** cerebrum, **(d)** eye, **(e)** heart, **(f)** liver, **(g)** lung, **(h)** muscle, **(i)** placenta, **(j)** spleen, and **(k)** thymus. For each tissue, the cell cycle embedding using mNeurosphere reference is given in the first sub-panel, tissue-level UMAPs from Cao et al. (2020) colored by cell cycle position *θ* in the second sub-panel, tissue-level UMAPs from Cao et al. (2020) colored by cell type in the third sub-panel, percentage of actively proliferating cells for each cell type in decreasing order in the forth sub-panel, tissue-level UMAPs from Cao et al. (2020) colored by development days in the fifth sub-panel, and percentage of actively proliferating cells for each development day in the last sub-panel. with **(g)** lung, **(h)** muscle, **(i)** placenta, **(j)** spleen, and **(k)** thymus.

**Supplementary Figure S27.**
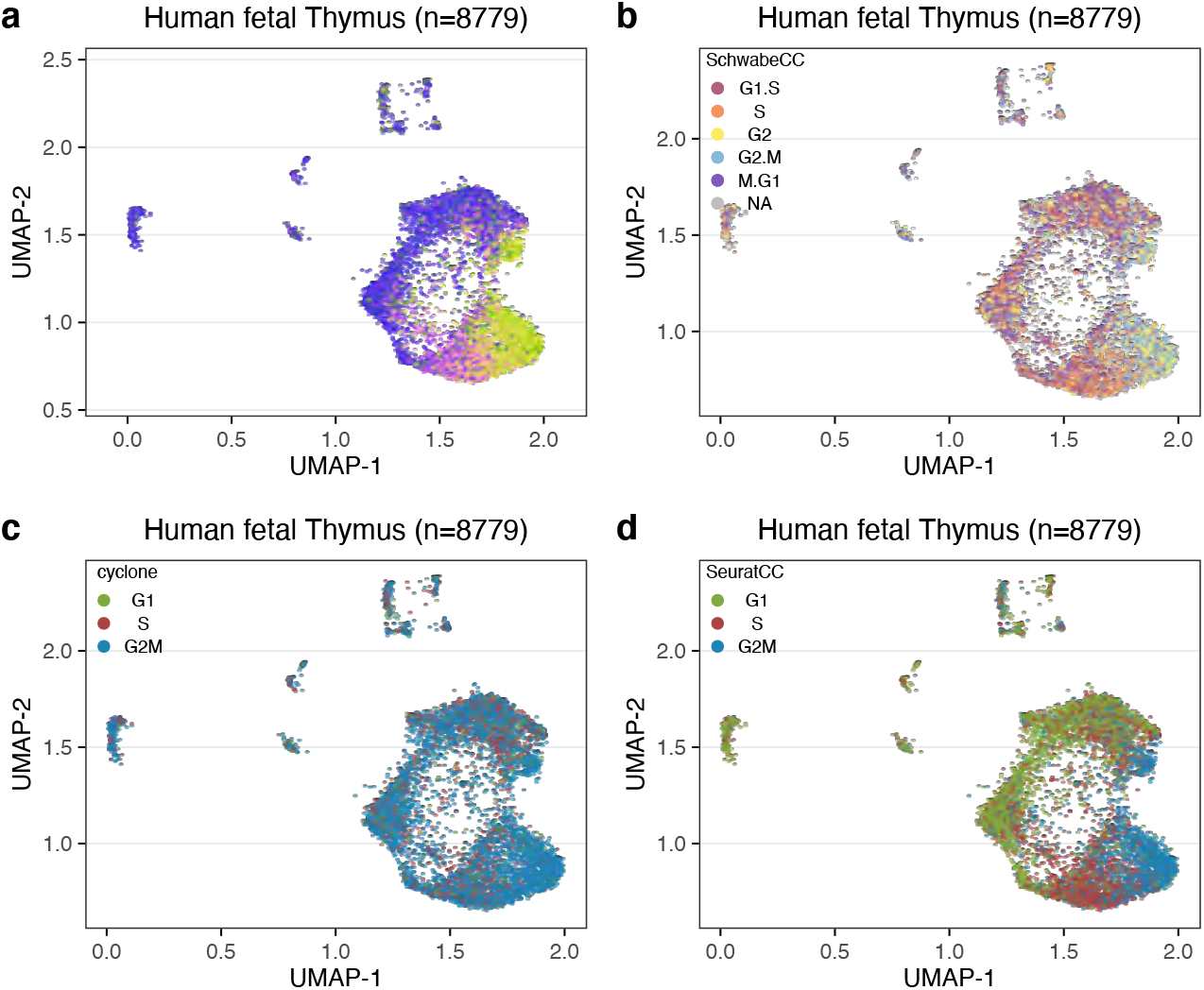
Human fetal thymus UMAPs colored by cell cycle position or stage. **a)** Same as Supplementary Figure S26k second sub-panel, which shows the UMAP embeddings of human fetal thymus, colored by cell cycle position *θ*. **(b)** Same UMAP embedding as in (a), but colored by 5 stage cell cycle representation, inferred using the SchwabeCC method from Schwabe et al. (2020). **(c)** Same UMAP embedding as in (a), but colored by 3 stage cell cycle representation, inferred by cyclone (Scialdone et al., 2015). **(d)** Same UMAP embedding as in (a), but colored by 3 stage cell cycle representation, inferred by Seurat (Stuart et al., 2019).

**Supplementary Figure S28.**
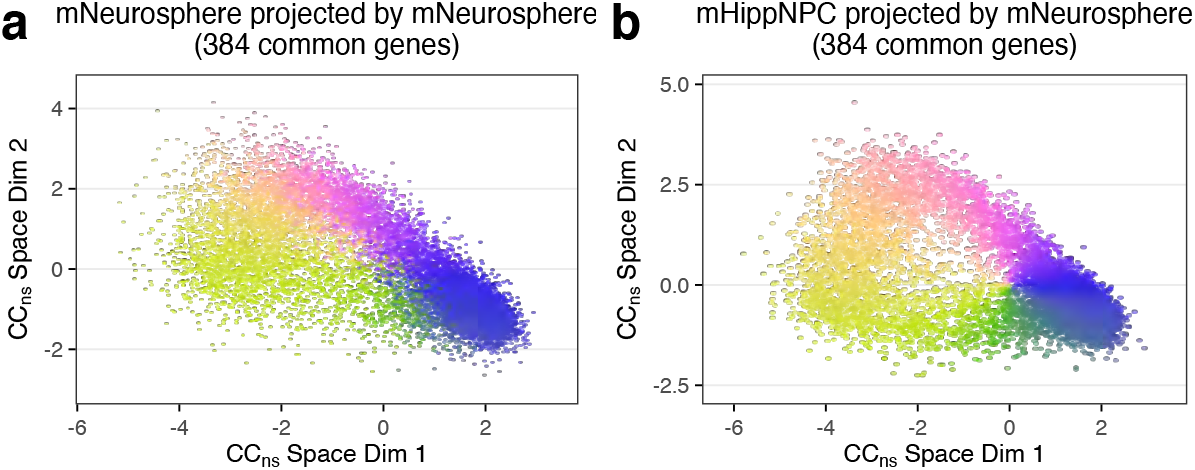
Projection using the exact same genes on two datasets. This figure shows cell cycle embeddings for **(a)** mNeurosphere and **(b)** mHippNPC dataset using the subset mNeurosphere reference restricted to 384 genes existing in both datasets. Cells are colored by the cell cycle position *θ*.

**Supplementary Figure S29.**
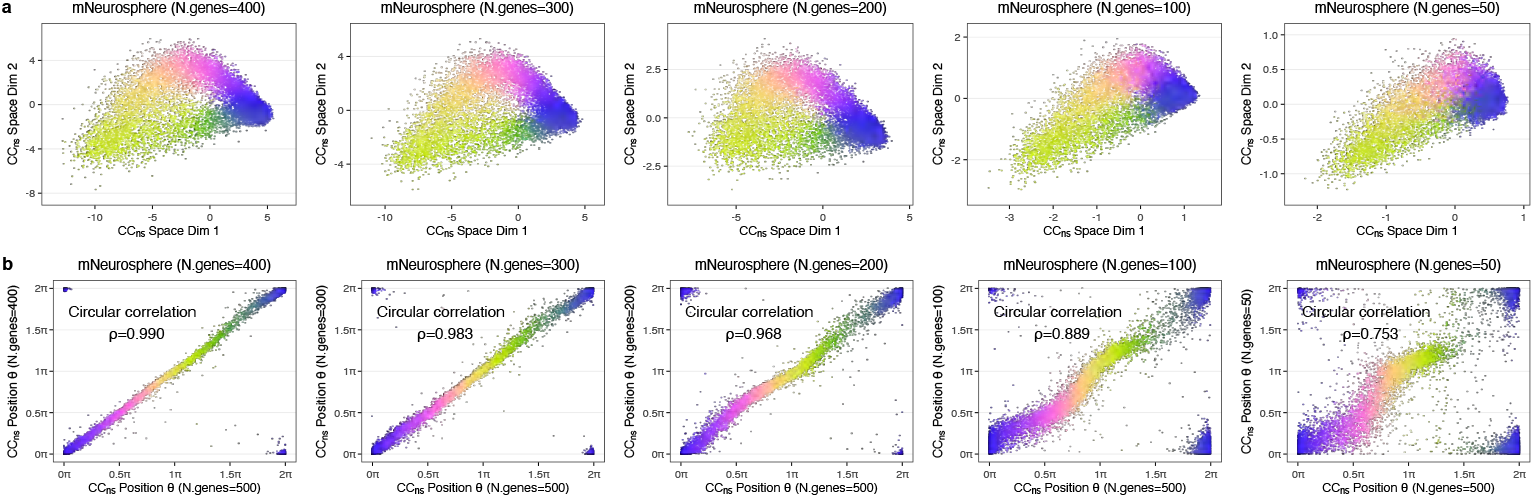
Examples of mNeurosphere dataset projections with randomly sub-sampled projection matrices. Each column represents an example of a projection using the sub-sampled genes from original 500 projection genes. From left to the right, the numbers of genes are 400, 300, 200, 100, and 50. **(a)** The projected cell cycle embedding using the sub-sampled projection matrix. Cells are colored by cell cycle position inferred using all 500 projection genes. **(b)** Comparisons of cell cycle positions *θ* estimated using the full 500 projection matrix and using the sub-sampled projection matrix. The circular correlation *ρ* is given in the figure.

**Supplementary Figure S30.**
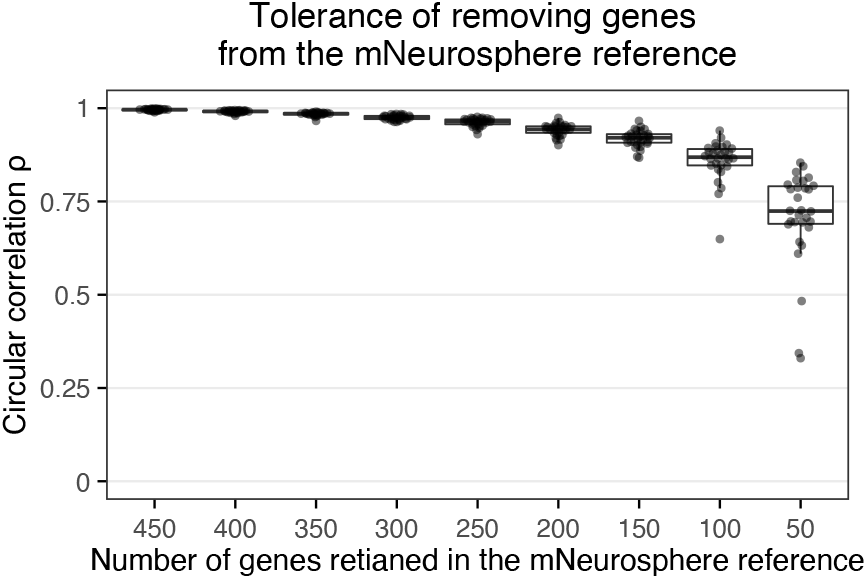
Stability assessment with projection genes missing. This figure shows comprehensive assessments as complement to Supplementary Figure S29. For each target number of genes retained in the mNeurosphere reference matrix, we randomly sampled different genes 30 times. For each run, the circular correlation coefficient *ρ* was calculated between *θ* from projection using the full reference matrix and *θ* from projection using sub-sampled reference.

**Supplementary Figure S31.**
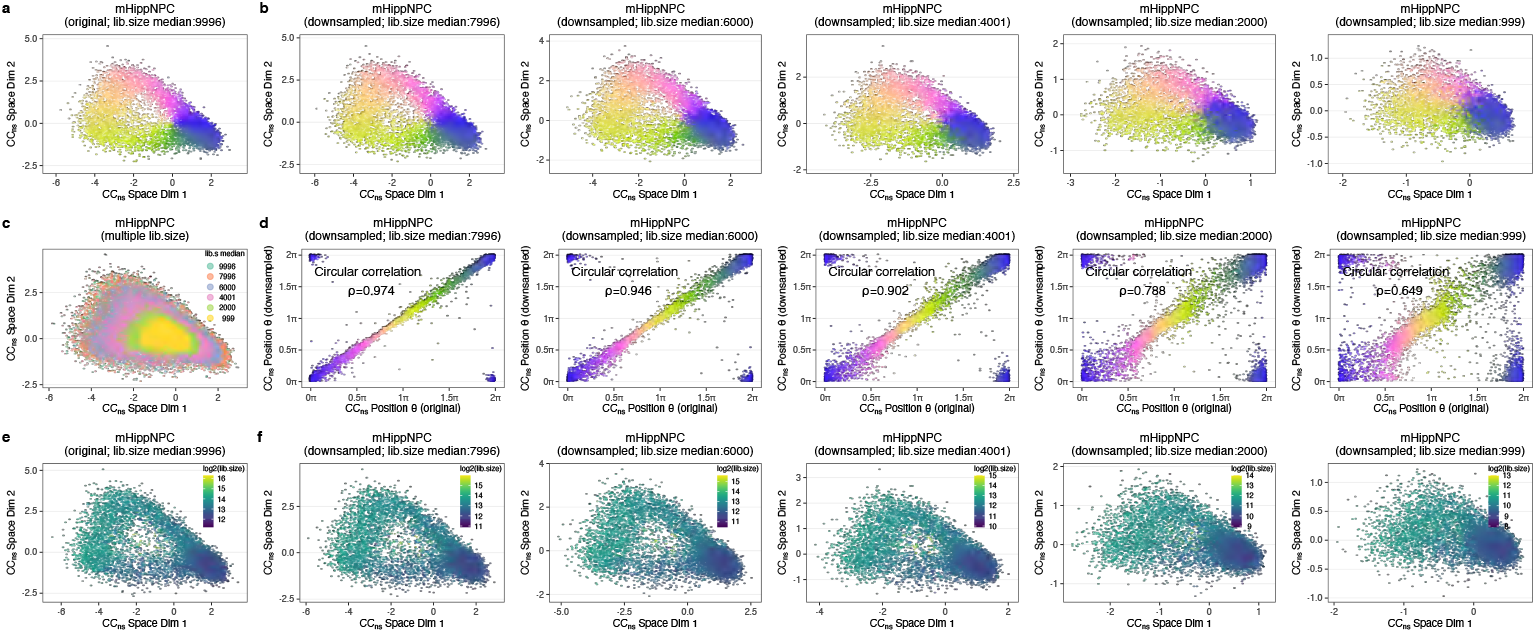
Examples of projections on downsampled mHippNPC dataset. **(a)** The cell cycle embedding projection using the mNeurosphere reference on original mHippNPC data. Each point is a cell, colored by cell cycle position (polar angle in the embeddings). **(b)** Each sub-panel represents the same projection as in (a), but the mHippNPC is downsampled to the 80%, 60%, 40%, 20%, and 10% of its original library size (corresponding to median of library size is given in the panel title). Note that the ranges of both x-axis and y-axis are different across sub-panels. Cells are colored by the cell cycle position inferred on original data. **(c)** By overlaying (a) and all sub-panels of (b), it shows the shrinkage of projections with library size decreasing. **(d)** Comparisons of cell cycle positions *θ* estimated from the original mHippNPC data and from the downsampled mHippNPC data. Cells are colored by the cell cycle position inferred on original data (x-axis). **(e)** Similar to (a), but the points are colored by log_2_ transformed library size. **(f)** Similar to (b), but the points are colored by log_2_ transformed library size.

**Supplementary Figure S32.**
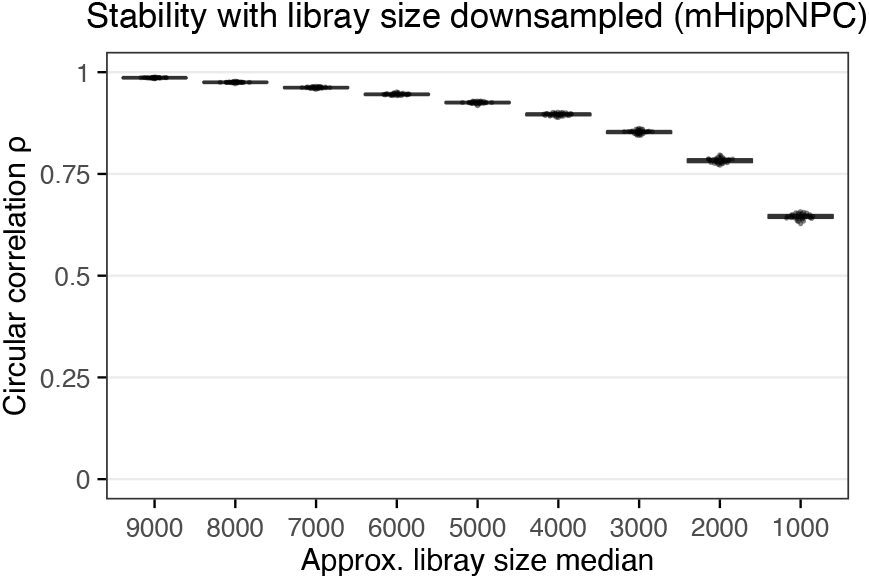
Stability assessment with decreasing sequencing depths. This figure shows comprehensive assessments as complement to Supplementary Figure S31. We repeated the downsampling processes for mHippNPC for each target downsampling percentage. For each run, the circular correlation coefficient *ρ* was calculated between *θ* estimated on the original mHippNPC data and *θ* estimated on the downsampled data.

**Supplementary Figure S33.**
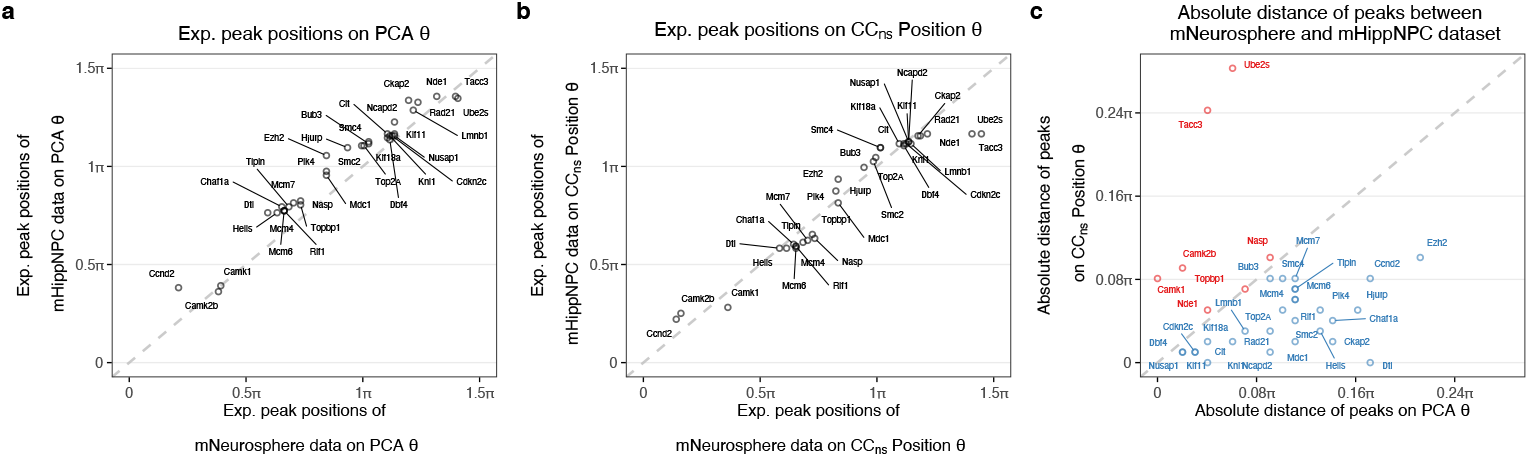
Comparison of positions of peak expression for *θ* estimated on independent PCA and projection by mNeurosphere reference. **(a)** For each gene dipicted in Supplementary Figure S6, we estimate and compare when the peak expression is reached between 0 to 2*π* for mNeurosphere and mHippNPC data. The position *θ* is based on independent PCA on GO cell cycle genes of each data. **(b)** Similar to (a), but now we use position *θ* estimated using pre-learned mNeurosphere reference. **(c)** The majority of genes are better aligned on *θ* pre-learned mNeurosphere reference. x-axis represents the absolute distance of position of peak expression on *θ* estimated on independent PCA, while y-axis represents those estimated using pre-learned mNeurosphere reference. Genes with a larger absolute distance on *θ* estimated on independent PCA compared to *θ* estimated using pre-learned mNeurosphere reference are colored as blue, and genes are colored by red if showing the opposite direction.

**Supplementary Figure S34.**
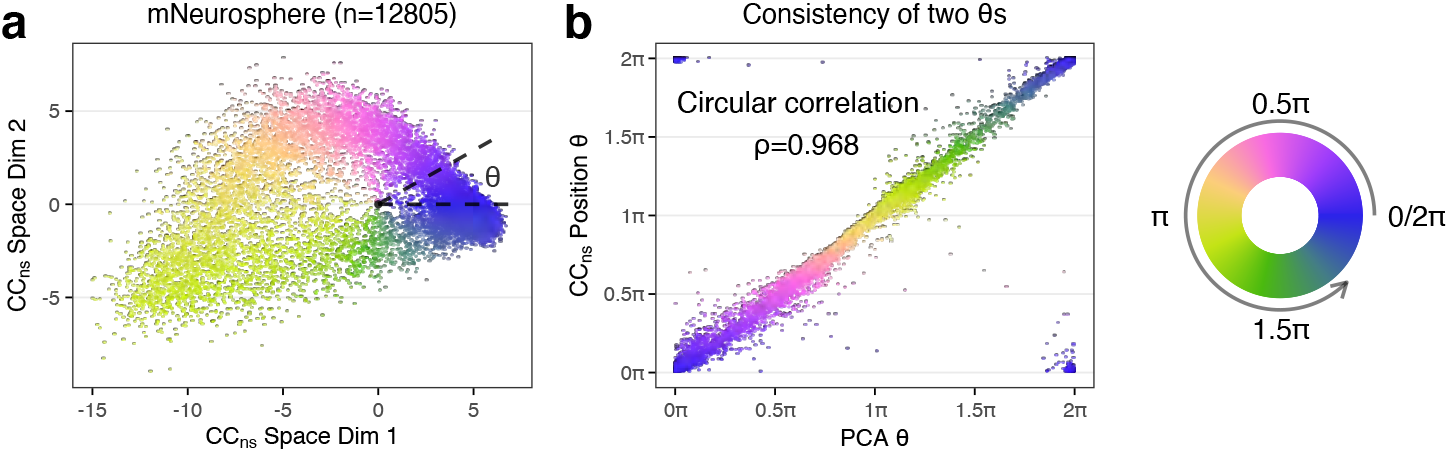
Self-projection to test method sensitivity on a positive control. **(a)** The cell cycle embedding of mNeurosphere data using the reference learned from itself. Note the projections are different from direct PCA, as the PCA is done on Seurat corrected expressions while the projection is calculated on non-corrected expressions. **(b)** Comparisons of cell cycle positions *θ* estimated from the direct PCA and from the projection. Cells are colored by cell cycle positions *θ* estimated from the projection.

**Supplementary Figure S35.**
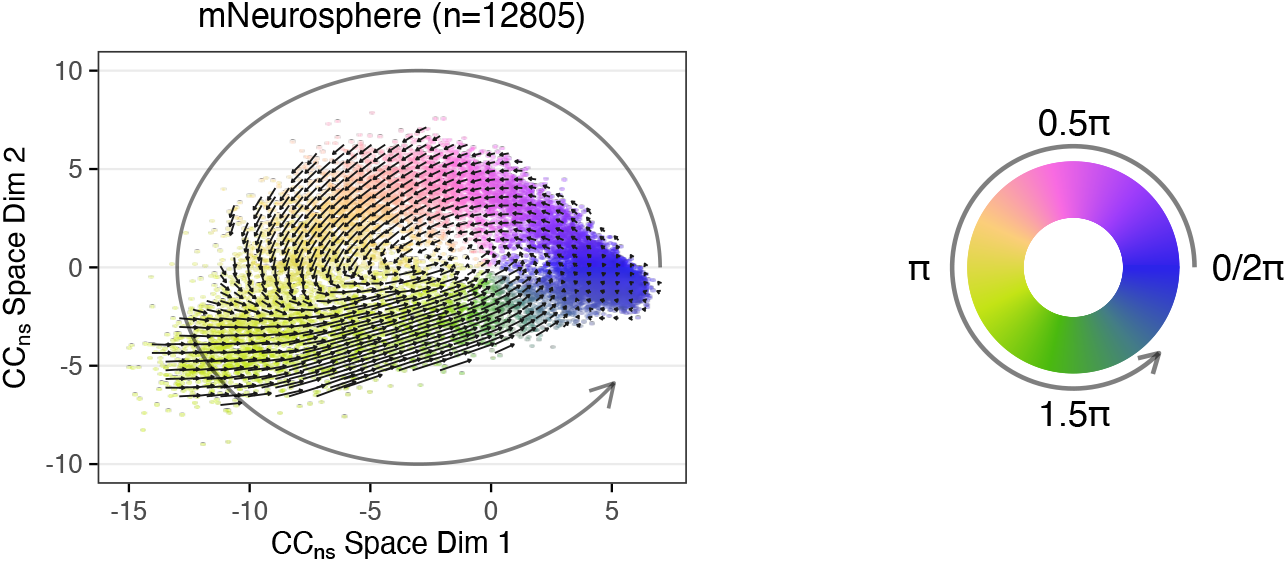
RNA velocity projection into the cell cycle embedding. The arrows are projected RNA velocity estimations from the RNA velocity dynamical model in Bergen et al. (2020). Cells are colored by our cell cycle positions *θ*. The directions of RNA velocity projections are consistent with the directions of cell cycle positions *θ*.

